# Generalism vs. specialization: Does niche breadth influence species responses to anthropogenic land-use change in Neotropical leaf-cutter ants?

**DOI:** 10.64898/2026.08.12.744409

**Authors:** Diego Garcia Castillo

## Abstract

Land-use change, such as the transformation of woody ecosystems into open pastures, acts as a strong ecological filter, favouring some species while excluding others according to differences in ecological niche breadth. Understanding how differences in niche breadth influence species responses under anthropogenic filters is crucial to anticipate their persistence or displacement. In this study, we quantified realized niche breadth in two sympatric ecosystem engineers, the Neotropical leaf-cutter ants *Atta cephalotes* and *Atta laevigata*, to test whether breadth differences are consistent with specialist and generalist ecological strategies. We characterized realized niche breadth across fine-scale environmental gradients by integrating hemispherical photography, microclimatic data, mound architecture, and edaphic profiles from 114 colonies across a regional transect in the Colombian Andes, alongside macroclimatic data from Copernicus. Principal Component Analysis (PCA) and PERMANOVA identified canopy openness and bushes- and tree-type vegetation density as the principal axes of interspecific niche partitioning. The observed differences in realized niche breadth were consistent with specialist and generalist ecological strategies. *A. laevigata* was predominantly associated with open-canopy areas, warmer micro- and macroclimatic conditions, and narrower edaphic dispersion. In contrast, *A. cephalotes* occupied a wider range of microhabitat conditions. This broader realized niche breadth is compatible with previous reports of *A. cephalotes* occurring in urban areas. Together, these findings suggest that niche breadth may influence how Neotropical leaf-cutter ants respond to habitat transformation, helping to understand the ecological consequences of land-use change.

## Introduction

Land-use change in the Anthropocene acts as a strong thermal and structural filter for biodiversity, particularly across transitions between natural and human-modified habitats [1,2]. Rapid environmental change has been triggered by the increasing demand for agricultural production that accelerates the conversion of complex woody habitats into croplands, pastures, and other simplified environments [3–5]. As a consequence, species may undergo local extirpation, respond through phenotypic plasticity or heritable genetic changes, or shift their distributions into novel or previously restricted habitats [6–8]. Understanding the mechanisms behind these disparate outcomes is of critical ecological interest; specifically, why certain taxa successfully expand into anthropogenic mosaics while others decline or become locally extinct [8–10].

Identifying the ecological mechanisms that determine which species persist in anthropogenic mosaics remains a critical challenge. Niche research has traditionally focused on its position, understood as the environmental optima or “average” preferences of a species, to predict response to habitat modification [11,12]. However, niche position alone often fails to account for the complexity of human-disturbed landscapes. Urbanized and agricultural landscapes often combine environmental conditions (e.g. temperature, vegetation structure, soil characteristics, and resource availability) in ways that species may not have previously experienced [13]. In contrast to the relative stability of natural environments, human-modified landscapes are often highly fragmented, constantly fluctuating, and perturbed [14,15]. Recent studies increasingly recognize niche breadth as an important determinant of species persistence under environmental change because it defines the range of abiotic conditions and biotic resources that species can tolerate and exploit [16–21]. Broader niche breadth is generally associated with greater ecological generalism, whereas a narrower niche breadth is associated with greater ecological specialization [19,22]. Consequently, species with relatively narrow niche breadths are expected to be more vulnerable to the environmental filtering associated with land use change [9,18]. On the other hand, broader niche breadth may facilitate species persistence by enabling more tolerance across novel combinations of environmental conditions [18,19].

The greater persistence of species with broader realized niche breadths in human-modified landscapes has been documented across a wide range of taxa [9,18,19]. For instance, avian generalists are more likely to maintain stable populations in urban centres than specialists [23–25]. Similar patterns have been observed in rodents, bats, reptiles, amphibians, and insects, where a positive relationship exists between niche breadth and occupancy in landscapes altered by humans [26,27]. Recent work on butterflies further supports this pattern, showing that species with broader environmental niches are more likely to persist across agricultural landscapes than habitat specialists [28]. However, many previous studies infer niche breadth from species’ presence–absence records or broad habitat categories, providing limited insight into how fine-scale environmental gradients shape realized niche breadth and species persistence [19,28,29]. Consequently, whether differences in realized niche breadth measured across fine-scale environmental gradients explain the contrasting persistence of ecosystem engineers in human-modified landscapes remains largely unknown. Ecosystem engineers are taxa that physically modify their environment [30,31]. Understanding how they respond to novel environmental conditions arising from rapid land-use change is therefore critical, as their persistence or exclusion can have cascading effects on soil structure, nutrient cycling, and ecosystem functioning [32,33].

The Neotropical leaf-cutting ants *Atta cephalotes* and *Atta laevigata* (Figs 1A and 1B) provide an ideal system to investigate these dynamics. These closely related ecosystem engineers occur sympatrically across much of northern South America [34–36]. *Atta* colonies excavate large subterranean nests that alter soil structure and influence local microclimatic conditions [32,37]. At broader spatial scales, *Atta* species are generally considered generalists because they forage on a wide variety of plant species and occupy broad geographic ranges [38,39]. However, field studies suggest they may diverge in their resilience to disturbance; for instance, while both species have been documented persisting within heavily degraded matrices [40,41], *A. cephalotes* has been more frequently observed in urban centres [42,43]. Because these two species coexist sympatrically, contrasting patterns of persistence are less likely explained by broad macro-climatic variation or dispersal limitations. Instead, this system provides an opportunity to test whether differences in realized niche breadth measured across fine-scale environmental gradients are associated with contrasting responses to land-use change. We therefore hypothesize that *A. cephalotes* occupies a broader realized niche than *A. laevigata* across fine-scale abiotic and structural gradients.

**Fig 1.**
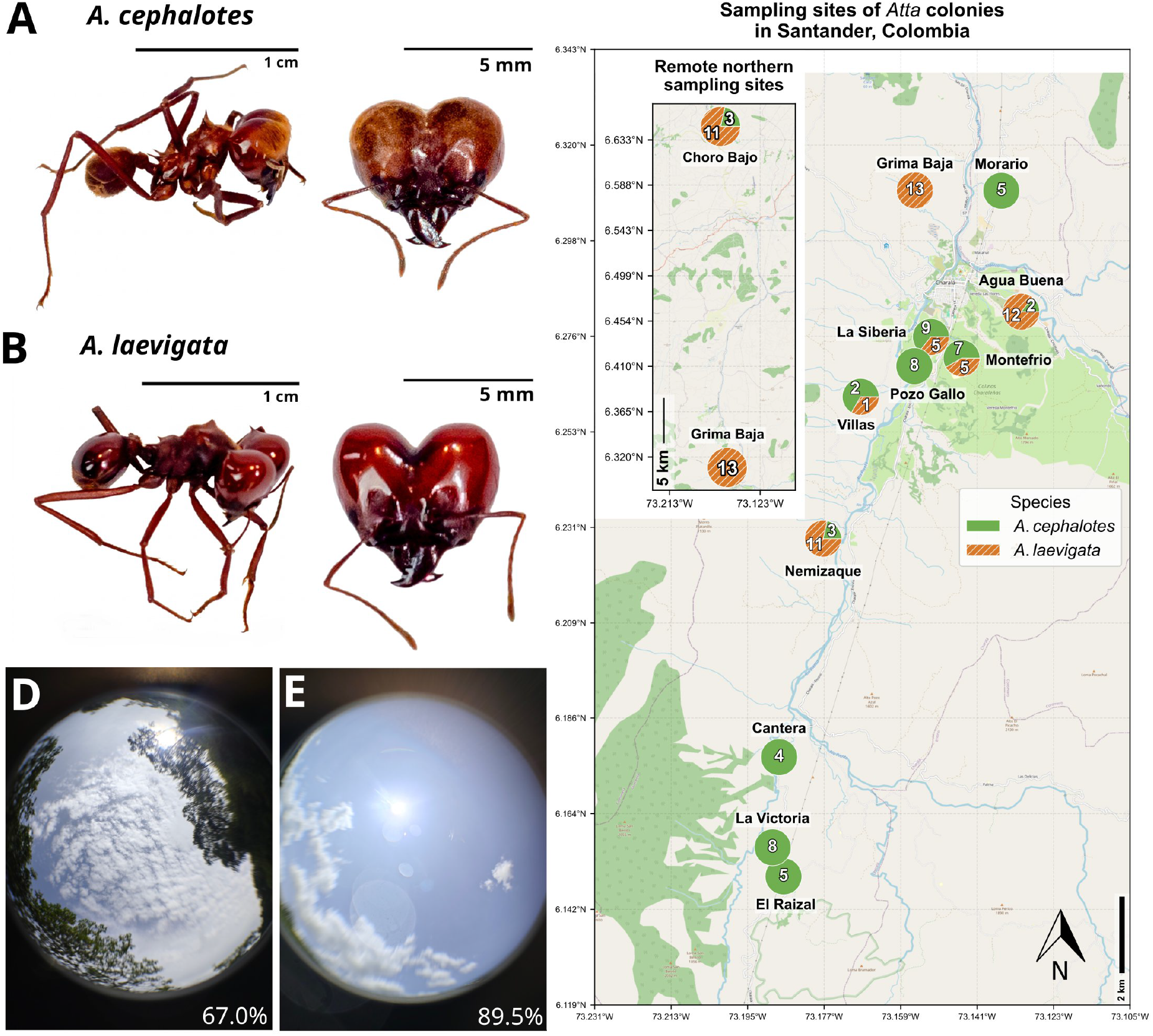
Sampling design and microhabitat characterization of *Atta* colonies in Santander, Colombia. (A) Morphological detail of an *A. cephalotes* soldier. (B) Morphological detail of an *A. laevigata* soldier. Both depicted individuals were collected from colonies situated in the “La Siberia” location (geographic coordinates detailed in S1 Table). (C) Topographic distribution of sampled colonies; markers represent individual nest locations with proportional pie charts indicating species composition at each site, where green represents *A. cephalotes* and orange with a diagonal hatch pattern represents *A. laevigata*. (D) Representative canopy cover of a typical *A. cephalotes* microhabitat; the selected image corresponds to a site with a canopy openness percentage (indicated in panel) approximating the mean estimate for the species. (E) Representative canopy cover of a typical *A. laevigata* microhabitat; the selected image corresponds to a site with a canopy openness percentage (indicated in panel) approximating the mean estimate for the species. Base map © OpenStreetMap contributors. Map generated using Python 3.12.3 with the Matplotlib, Geopandas, and Contextily libraries.

In this study, we characterized the fine-scale realized niche breadth and microhabitat associations of *A. cephalotes* and *A. laevigata* occurring sympatrically within the Andean region of Santander, Colombia. Our primary objective was to determine whether differences in realized niche breadth measured across fine-scale environmental gradients are associated with the contrasting persistence of these two species in human-modified landscapes. To characterize the realized niche of each species, we quantified the microhabitat parameters of 114 active colonies (*n* = 56 for *A. cephalotes*; *n* = 58 for *A. laevigata*; Figs 1A-C; S1 Table). For each nest, we measured canopy openness (Figs 1D and 1E), soil pH, mound surface conditions, and macro-vegetation type associations, mound surface temperature, while also quantifying mound architecture to explore whether nest characteristics differ between species. Utilizing multivariate dispersion metrics and affinity indices, we tested the hypotheses that *(i) A. laevigata* occupies a narrower realized niche than *A. cephalotes* and *(ii)* differences in realized niche breadth between the two species are associated with their contrasting occurrence in human-modified habitats. By testing these hypotheses, we help to clarify how differences in realized niche breadth may contribute to the contrasting responses of closely related ecosystem engineers to land-use change.

## Materials and methods

### Study area

The study area encompasses multiple locations in the department of Santander, Colombia across a south-to-north geographic transect spanning approximately 59 km. Specifically, 12 locations were sampled: 10 in the rural area of the municipality of Charalá, 1 in Ocamonte, and 1 in Barichara (S1 Table, Fig 1C). These locations, separated by several kilometres, were selected to capture variation in the regional distribution of both species in this part of the Colombian Andes.

### Sampling design

Within each location, we identified *Atta* colonies by systematically surveying field transects. We selected the first active nest that met the minimum distance criterion of 100 m from previously sampled colonies. We established this threshold based on the maximum foraging and architectural boundaries of both species; mature *A. cephalotes* surface foraging networks regularly extend up to 65–78 m [38], while subterranean excavations of mature *A. laevigata* nests demonstrate that underground foraging tunnels can reach up to 70 m radially [44]. Enforcing a minimum 100 m buffer zone minimized the likelihood of territorial overlap and increased the independence of all localized microclimatic and edaphic samples. Only active colonies were included, defined by the presence of at least two active entrance holes (confirmed by worker activity or freshly excavated soil). Sampling was conducted in open grasslands and adjacent woody areas, where both species are known to occur; deep forest interiors were not explored.

Species identification followed the diagnostic characters described by Mackay and Mackay [45]. During field surveys, *A. cephalotes* workers consistently exhibited a lighter and more opaque tegument with a distinct head-thorax contrast, whereas *A. laevigata* workers appeared darker with a distinctly shinier tegument. In addition, *A. laevigata* soldiers consistently displayed a more intense reddish colouration than those of *A. cephalotes* (Figs 1A and 1B). We used mound architecture as a secondary field indicator; during the sampling season, *A. cephalotes* typically presented more robust mounds with abundant freshly excavated soil, while *A. laevigata* mounds were flatter.

Three colonies in one specific location (Villas) were classified as “Unknown” (U). Although the individuals matched the morphological criteria for *A. laevigata*, their mounds were constructed of masticated leaf fragments rather than excavated soil. We excluded these three colonies from the main analyses but we fully reported them in S2 Table.

Most sampling was conducted in publicly accessible areas intersected by public roads and footpaths, where no access permits were required. For colonies located within privately owned properties (including Choro Bajo, Ocamonte, Grima Baja, El Raizal, and Montefrío), field access was granted with verbal permission from the respective landowners or property managers, several of whom accompanied the sampling. The study was based on in situ measurements of active colonies. Two soldier ants were collected solely for photographic documentation (Fig. 1A–B) and euthanised by freezing prior to photography. We did not preserve or use individuals for tissue sampling or other biological analyses. Consequently, no scientific collecting permit was required.

### Edaphic assessment

We measured soil pH by collecting a sample from a major active nest entrance. All samples were stored separately in plastic bags and air-dried for 48 h before the analysis. We prepared a 1:5 (w/v) soil-to-water suspension by mixing 2 g of soil with 10 ml of distilled water. After stirring, we allowed the mixture to settle for 10 minutes. We then determined pH using non-bleeding indicator strips (pH 0–14, MColorpHast™). Each strip was submerged for one minute, and the resulting colour was immediately compared against the reference scale of the manufacturer. All colour assessments were performed by the same observer under standardized lighting conditions.

### Temperature recording

Internal and external temperatures in degrees Celsius (°C) were measured *in situ* using an HTC-2 digital thermometer positioned immediately above the mound surface. To record internal colony temperature, the device’s remote sensor probe was inserted approximately 10 cm into a major active nest entrance. The device remained in place for approximately five minutes until the temperature reading stabilized. We then recorded the internal colony temperature together with the ambient air temperature measured by the device’s integrated sensor.

### Canopy openness and image processing

We characterized canopy openness using hemispherical photography with a 198° fisheye lens. Images were captured using a Redmi Note 14 Pro+ 5G smartphone (3:4 aspect ratio; 3060 x 4080 px resolution; no flash). The device was mounted on a tripod, levelled parallel to the mound surface at its centre, and oriented upwards. Photographs were taken using a remote shutter to avoid camera shake.

During fieldwork, GPS watermarks were initially embedded in the images. However, we identified that these watermarks were unreliable due to satellite signal inconsistencies. We then removed all watermarks during post-processing to ensure data integrity and avoid interference with pixel analysis.

We developed an image processing pipeline in Python 3.12 using the OpenCV library. To exclude lens barrel artifacts, we defined a circular Region of Interest (ROI) using a binary calibration mask derived from a reference photograph of a uniform white surface (’*Mask_White_FishEye.jpeg*’). To standardize lighting variations, a differential gamma correction was applied based on sky conditions recorded during sampling (Clear sky: γ = 1.0; Partially cloudy: γ = 1.5; Overcast: γ = 2.0; S2 Table). Images were converted to greyscale and binarized using Otsu’s thresholding algorithm. Canopy openness was calculated as the ratio of white pixels (sky) to total pixels within the ROI, see *S1 Supplementary Dataset* for the photographs and associated light percentage estimates.

### Vegetation assessment

We characterized the vegetation surrounding each colony within a 10 m radius from the mound edges. We recorded the presence of three general categories: Grasses (G), Bushes (B), and Trees (T). These categories were not mutually exclusive, allowing for the characterization of mixed vegetation structures at each site.

### Mound characterization

We recorded external mound dimensions in centimetres (cm) using a standard measuring tape. Maximum length (*L*) was defined as the distance between the two most distant active entrance holes, while width (*W*) was measured as the maximum distance perpendicular to the length axis. We used these measurements to characterize the surface footprint of the colonies rather than for detailed architectural analysis. Assuming an elliptical shape, we estimated mound area (cm^2^) using the semi-major (*L*/2) and semi-minor (*W*/2) axes. We also calculated the elongation index as the *L:W* ratio. An elongation index of 1.0 indicates a circular mound and values > 1.0 indicate increasing ellipticity.

### Statistical analysis

All statistical analyses were performed in Python 3.12.3 using the libraries scikit-learn version 1.8.0 [46], scipi version 1.17.1 [47], and scikit-bio version 0.7.2 [48]. Prior to statistical analyses, we assessed the distribution of all continuous variables using Shapiro–Wilk tests as well as the homogeneity of variances using Levene’s tests (S3 Table). Because most variables deviated from normality, species comparisons were performed using Permutational Multivariate Analysis of Variance (PERMANOVA), a permutation-based approach that does not rely on multivariate normality.

#### Analysis of environmental and architectural variables

To assess species differences in continuous environmental (soil pH, temperature, canopy openness) and architectural (mound area, shape ratio) variables, we performed separate one-factor PERMANOVA analyses based on Euclidean distance matrices following Anderson (2001) [49]. Euclidean distance matrices were calculated from standardized continuous measurements after *z*-score transformation. These matrices represented dissimilarities between colonies. Significance was assessed using 999 permutations directly on the Euclidean dissimilarity matrices. Species comparisons involved two groups (114 colonies), whereas geographic location analyses included 12 localities. Group dispersions were subsequently assessed using PERMDISP [50] to determine whether significant PERMANOVA results reflected differences in group centroids rather than differences in multivariate dispersion.

We calculated effect sizes (*R*²) only for species comparisons. The scikit-bio implementation of PERMANOVA reports pseudo-*F* statistics and permutation-based P-values but does not provide effect sizes. Thus, we algebraically derived *R*² values from the pseudo-*F* statistic for one-factor PERMANOVA models. We used the relationship between the pseudo-*F* ratio and the proportion of explained sums of squares as follows:

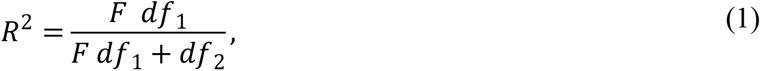

where *F* is the pseudo-*F* statistic, *df*_1_ is the numerator degrees of freedom *g* -1, and *df*_2_ is the denominator degrees of freedom *N* - *g*. We reported effect sizes only for species because these represented the primary explanatory factor of this study. Geographic location was evaluated separately to assess spatial heterogeneity rather than as a competing explanatory model.

#### Multivariate niche characterization

We performed a Principal Component Analysis (PCA) of the standardized environmental matrix to visualize the realized niche overlap and summarize the main environmental gradients separating the two species. We used the PCA implementation of scikit-learn (version 1.8.0) following the classical formulation described by Jolliffe (2002) [51]. Categorical vegetation data (Trees, Bushes, Grasses) were integrated into the PCA as binary presence/absence flags. A multivariate PERMANOVA [49] based on the Euclidean distance matrix of dissimilarity derived from these variables (999 permutations) was used to test for significant realized niche partitioning between species. We used the PERMANOVA implementation of the scikit-bio library as described before.

#### Habitat affinity and niche breadth

We evaluated habitat preferences using Indicator Value (IndVal) analysis to identify significant associations between each species and vegetation categories based on specificity (A) and fidelity (B). This analysis was implemented using NumPy (v2.4.4) and Pandas (v3.0.3), following the formulations of Dufrêne & Legendre (1997) and De Cáceres & Legendre (2009) [52,53]. Finally, realized niche breadth was quantified using three dispersion metrics: Standard Deviation (SD), Coefficient of Variation (CV), and Interquartile Range (IQR). All results were considered significant at *p* < 0.05.

### Regional climatic characterization

To characterize the regional climatic context of the sampling sites, we extracted regional climatic variables from the ERA5-Land reanalysis dataset [54]. We retrieved data via the Copernicus Climate Data Store (CDS) API using the Python library cdsapi version 0.7.7. The study area was defined by a bounding box spanning 8.0° to 6.0° N and 74.5° to 72.0° W, covering a ∼150 km × 150 km region included the sampled locations.

We processed the downloaded climatic datasets using the Python libraries xarray and rioxarray as follows:

#### Average temperature

The *2m_temperature* (*t2m*) data correspond to the month of February 2024, 2025, and 2026. We chose those three years to assess whether the regional thermal pattern during the sampling month was consistent across years. For each year, the hourly values were averaged over the 24-hour cycle and across the entire month to generate a monthly mean temperature. Units were converted from Kelvin to Celsius (°C).

#### Total annual precipitation

We used the *total_precipitation* (*tp*) variable for the full years 2024 and 2025 for January–July 2026. Because 2026 was still in progress at the time of analysis, we treated precipitation data for that year as a partial-year accumulation and we did not directly compare with the annual totals for 2024 and 2025. As this variable is provided as an accumulated value within each forecast day, only the final hourly recording (24:00) of each day was extracted to represent the daily total. We then summed such daily values to calculate the total annual accumulation in meters (m).

#### Total annual evaporation

Data for *evaporation_from_bare_soil* (*evabs*) were retrieved for 2024 and 2025 and for January–July 2026. We treated 2026 data as a partial-year accumulation because 2026 was still in progress at the time of analysis, and therefore this year was not directly compared with the annual totals for 2024 and 2025. Similar to precipitation, this is an accumulated variable; therefore, we extracted only the final hourly record of each day. To reflect total water loss from the surface, we calculated the absolute values (converting the model’s negative flux convention to positive) and multiplied them by 1,000 to report the final values in millimetres (mm).

We reprojected the final processed rasters from their native geographic coordinate system (WGS84, EPSG:4326) to Web Mercator (EPSG:3857) to ensure spatial alignment with the topographical base maps used for visualization. Sampling locations were overlaid on the resulting climatic maps to facilitate visual comparison with regional environmental gradients.

### Use of AI-assisted technologies

During the preparation of this work, we used AI-powered technologies to assist with language editing and programming support. Specifically, we used Gemini (Gemini 3 Flash, Google) and ChatGPT-5.6 to improve writing clarity, correct grammatical errors, and reduce wordiness. We used these tools through targeted prompts (e.g., “Please correct wordiness and grammar of the following statement…”). Gemini was further used to assist in drafting and debugging Python code developed by the authors for image processing, data handling, and statistical analyses. In all cases, AI-assisted outputs were critically reviewed, verified, and revised by the authors before incorporation into the manuscript or analytical workflow. We did not use AI tools to generate the study hypotheses, experimental design, field data, statistical analyses, ecological interpretations, or scientific conclusions. All source data were generated by the authors, and all analytical decisions and interpretations remain entirely the responsibility of the authors.

## Results

### Realized niche characterization reveals fine-scale ecological differences between sympatric *Atta* species

To characterize the realized niches of both species, we quantified architectural (mound area and mound shape ratio), edaphic, and thermal profiles, along with canopy openness and vegetation affinity, for 114 *Atta* colonies across a heterogeneous landscape (Fig 1; S1 Fig; S1 and S2 Tables). Normality tests were conducted prior to analysis (S3 Table). Univariate PERMANOVA revealed that species identity significantly influenced most continuous variables, including mound area, soil pH, canopy openness, internal mound temperature, and thermal regulation capacity (*p* < 0.05; Table 1). Mounds of *A. laevigata* (69.1 ± 61.9 m²) were significantly larger than those of *A. cephalotes* (45.0 ± 37.5 m²). In contrast, mound shape ratio remained comparable between species (*A. laevigata*: 2.0 ± 1.0; *A. cephalotes*: 1.7 ± 0.9). Mean shape ratio for both species exceeded 1.0, indicating an elongated rather than circular mound geometry. Soil pH was slightly higher at *A. laevigata* (4.8 ± 0.2) than at *A. cephalotes* (4.6 ± 0.3) nests. Canopy openness showed the largest species effect with *A. laevigata* occurring in significantly more exposed sites (89.5 ± 6.9%) compared to *A. cephalotes* (67.6 ± 18.5%). Internal mound temperature also differed between species, with *A. laevigata* maintaining higher internal temperatures (25.1 ± 1.3°C) than *A. cephalotes* (24.0 ± 2.2°C; Table 1; S2 Fig). Thermal regulation capacity also differed between species (*A. laevigata*: -10.3 ± 4.7°C; *A. cephalotes*: -7.0 ± 4.3°C). PERMDISP revealed significant differences in within-species dispersion (interpreted here as realized niche breadth) for canopy openness, soil pH, and mound area, whereas thermal regulation capacity, internal temperature, and mound shape ratio did not differ in within-species dispersion. Geographic location explained additional variation in most environmental variables, although species identity remained significant across the principal ecological traits.

**Table 1.**
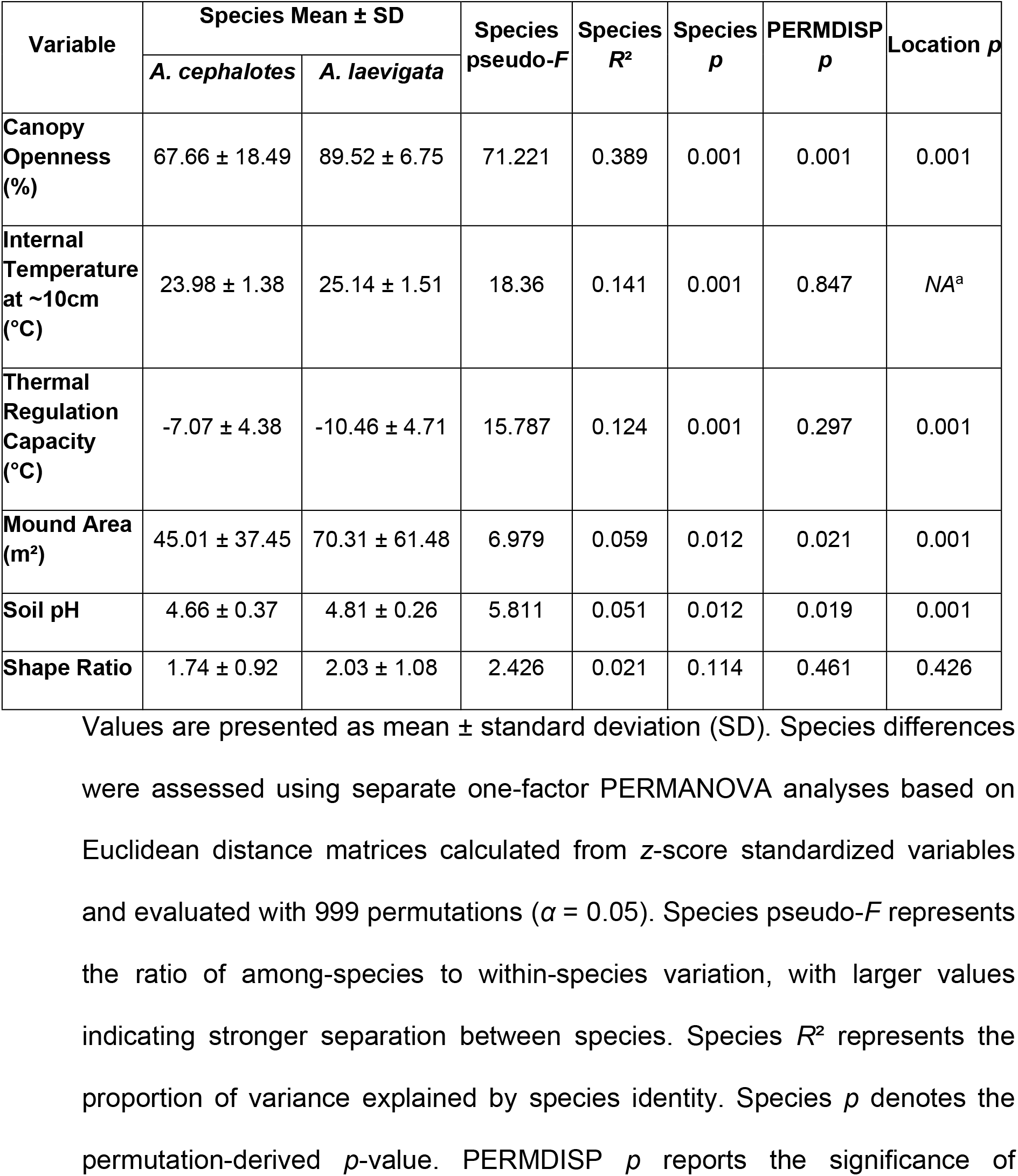
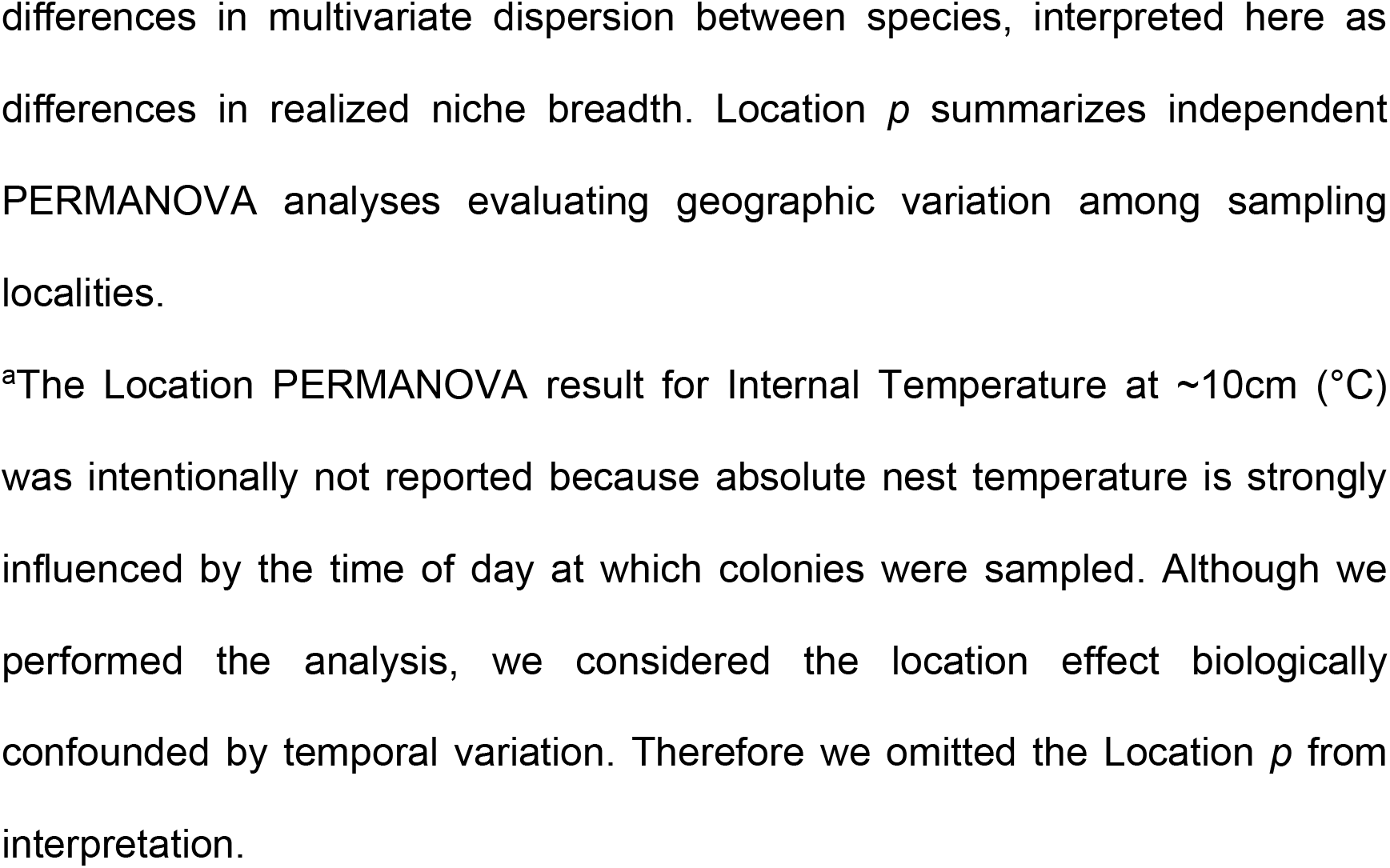
Comparison of environmental and architectural variables between *A. cephalotes* and *A. laevigata*.

| Variable | Species Mean ± SD | | Species pseudo-*F* | Species *R*<sup>2</sup> | Species *p* | PERMDISP *p* | Location *p* |
| --- | --- | --- | --- | --- | --- | --- | --- |
|  | *A. cephalotes* | *A. laevigata* |  |  |  |  |  |
| Canopy Openness (%) | 67.66 ± 18.49 | 89.52 ± 6.75 | 71.221 | 0.389 | 0.001 | 0.001 | 0.001 |
| Internal Temperature at ~10cm (°C) | 23.98 ± 1.38 | 25.14 ± 1.51 | 18.36 | 0.141 | 0.001 | 0.847 | NA<sup>a</sup> |
| Thermal Regulation Capacity (°C) | -7.07 ± 4.38 | -10.46 ± 4.71 | 15.787 | 0.124 | 0.001 | 0.297 | 0.001 |
| Mound Area (m<sup>2</sup>) | 45.01 ± 37.45 | 70.31 ± 61.48 | 6.979 | 0.059 | 0.012 | 0.021 | 0.001 |
| Soil pH | 4.66 ± 0.37 | 4.81 ± 0.26 | 5.811 | 0.051 | 0.012 | 0.019 | 0.001 |
| Shape Ratio | 1.74 ± 0.92 | 2.03 ± 1.08 | 2.426 | 0.021 | 0.114 | 0.461 | 0.426 |

Values are presented as mean ± standard deviation (SD). Species differences were assessed using separate one-factor PERMANOVA analyses based on Euclidean distance matrices calculated from z-score standardized variables and evaluated with 999 permutations ( $\alpha = 0.05$ ). Species pseudo-*F* represents the ratio of among-species to within-species variation, with larger values indicating stronger separation between species. Species *R*<sup>2</sup> represents the proportion of variance explained by species identity. Species *p* denotes the permutation-derived *p*-value. PERMDISP *p* reports the significance of
differences in multivariate dispersion between species, interpreted here as differences in realized niche breadth. Location $p$ summarizes independent PERMANOVA analyses evaluating geographic variation among sampling localities.
<sup>a</sup>The Location PERMANOVA result for Internal Temperature at ~10cm (°C) was intentionally not reported because absolute nest temperature is strongly influenced by the time of day at which colonies were sampled. Although we performed the analysis, we considered the location effect biologically confounded by temporal variation. Therefore we omitted the Location $p$ from interpretation.

To assess the extent of realized niche separation, we integrated environmental and architectural data with binary-transformed vegetation presence (trees, bushes, and grasses) into a Principal Component Analysis (PCA; Fig 2A). The first two components accounted for 47.9% of the total variance. PC1 explaining 34.2% of the variance largely separated the two species. Along this axis, species tended to occupy two regions of the multivariate space although, the 95% confidence ellipses overlapped near the multivariate centroids. Eigenvector analysis identified tree-type vegetation, bush-type vegetation, and canopy openness as the primary drivers of partitioning on PC1 (S5 Table). Thermal regulation capacity, mound area, and grass presence contributed to a lesser extent along this primary axis. Although soil pH differed significantly between species in the univariate analyses (Table 1), it contributed little to separation along PC1. Instead, soil pH loaded primarily on PC2 together with mound shape ratio (S5 Table). Further generalized linear modelling confirmed that this second multivariate axis reflects site-specific edaphic baselines rather than direct colony-mediated soil modifications (see *S1 Supplementary Text* for full model outputs). Multivariate PERMANOVA confirmed significant realized niche differentiation between species (pseudo-*F* = 27.01, *p* = 0.001), indicating that between-species multivariate differences greatly exceeded within-species variation.

**Fig 2.**
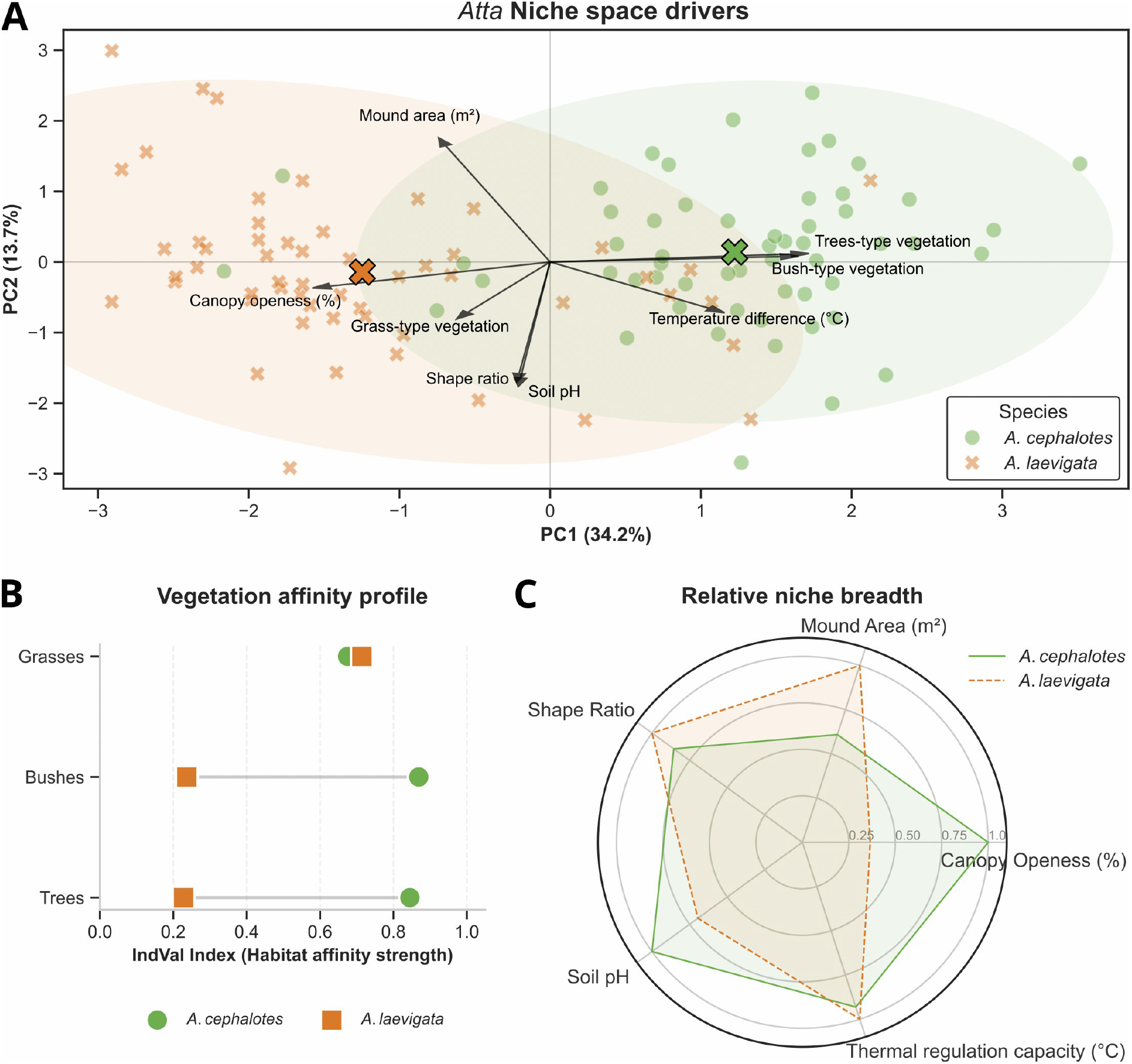
Fine-scale realized niche characterization and habitat affinity. (A) Principal Component Analysis (PCA) of environmental and architectural niche space for *Atta* species. Points represent individual colonies of *A. cephalotes* (green; n = 56) and *A. laevigata* (orange; n = 55). Crosses (X) indicate group centroids, and shaded areas represent 95% confidence ellipses for each species. The axes (PC1 and PC2) together explained 47.9% of the total variance. Loading vectors (arrows) indicate the direction and relative magnitude of the contribution of each environmental and architectural variable to the principal components. Vector direction indicates the relationship of each variable with the principal components, whereas vector length reflects its relative contribution to the species separation. Significant multivariate differentiation between species was confirmed in realized niche space via PERMANOVA based on Euclidean distances (pseudo-*F* = 27.01, *df* = 1, 109, *p* = 0.001, 999 permutations). (B) Vegetation affinity profile based on Indicator Value (IndVal) analysis for Trees, Bushes, and Grasses. The horizontal lines (dumbbells) represent the ecological distance in habitat association between the two species, where higher IndVal scores denote stronger specificity and fidelity to a vegetation category. (C) Radar plot comparing the relative multivariate dispersion (used here as a proxy for realized niche breadth) across five architectural and environmental variables for *A. cephalotes* (green) and *A. laevigata* (orange). For each variable, niche breadth was calculated from species-specific standard deviations (SD) and scaled relative to the maximum SD between species. Thus, a value of 1.0 represents the species with the greatest dispersion for that variable, whereas values below 1.0 represent proportional dispersion relative to that maximum. Values are therefore comparable between species within each variable, but not as absolute measures across variables. We evaluated differences in dispersion using PERMDISP (Table 1).

### Contrasting vegetation associations contribute to realized niche differentiation

Indicator Value (IndVal) analysis supported the PCA by revealing the same pattern of microhabitat differentiation between species (Fig 2B; S6 Table). Bushes and trees were highly specific to *A. cephalotes* (A = 0.783 and 0.785), whereas neither vegetation type had a high specificity for *A. laevigata* (A < 0.22). Grasses demonstrated high fidelity for both species (B > 0.94). However, woody vegetation showed much higher fidelity to *A. cephalotes* colonies (B = 0.964 and 0.911 for bushes and trees, respectively) but was infrequently present in the vicinity of *A. laevigata* colonies (B < 0.28). Together, these patterns indicate that *A. cephalotes* colonies were found across a wider range of vegetation structures, while *A. laevigata* was largely restricted to open grass-dominated environments. Consequently, bushes and trees emerged as strong indicators for *A. cephalotes* (IndVal > 0.84), whereas no vegetation category was uniquely associated with *A. laevigata,* largely because grasses occurred with high fidelity in both species.

### Realized niche breadth differed across environmental dimensions

We used two complementary metrics of within-species dispersion [SD and Interquartile Range (IQR)] to summarize differences in niche breadth for each environmental and architectural variable (Fig 2C; S4 Table). *A. cephalotes* exhibited substantially greater dispersion for canopy openness than *A. laevigata*, displaying an approximate 2.7-fold increase in SD (18.49 vs. 6.76) and a 6.1-fold increase in IQR (30.33 vs. 4.99). This pattern was consistent with the significant PERMDISP results (Table 1). For soil pH we observed a similar pattern of elevated niche breadth for *A. cephalotes* with a 1.4-fold higher SD (0.38 vs. 0.26) and a 2-fold higher IQR (0.60 vs. 0.30). Conversely, within-species dispersion in mound area was greater in *A. laevigata*, showing an approximate 1.6-fold increase in SD (61.49 vs. 37.45 m²) and a 1.5-fold increase in IQR (70.14 vs. 46.11 m²). Dispersion profiles of internal thermal regulation capacity and mound shape ratios were statistically comparable between both congeners (Table 1).

### Regional environmental context

ERA5-Land reanalysis data indicated that the study area spans a pronounced regional climatic gradient, providing the climatic context within which we evaluated realized niche breadth (S3–S5 Figs). The northernmost locality, Choro Bajo, was consistently warmer and drier than the remaining sampling localities. The Choro Bajo location showed average February temperatures of 22.92°C, 22.07°C, and 20.85°C in 2024, 2025, and 2026, respectively, and annual precipitation of approximately 2.98 m in 2024 and 2.84 m in 2025. In contrast, localities along the southern portion of the transect experienced lower temperatures and generally higher precipitation. February temperatures at El Raizal reached minima of 16.46°C, 15.36°C, and 14.76°C in 2024, 2025, and 2026, respectively. Annual precipitation across the southern and central sampling region ranged from approximately 3.72 to 4.38 m in 2024 and 4.30 to 5.03 m in 2025. Bare-soil evaporation also varied spatially, although its pattern was less clearly associated with the north–south climatic gradient (S5 Fig).Within the Andean transect evaluated here, this gradient coincided with the apparent southern occurrence limit of *A. laevigata*, whereas *A. cephalotes* remained present throughout the study area.

## Discussion

In this study, we characterized and compared the realized niche breadth of two sympatric ecosystem engineers across fine-scale environmental gradients, the leaf-cutting ants *A. cephalotes* and *A. laevigata*, in a region of the Colombian Andes. While both species overlapped across broad landscape categories within the zone of co-occurrence, they diverged most strongly at the fine-scale (Table 1; Fig 2A). This divergence was particularly pronounced for canopy openness, vegetation structure, and soil pH, in line with previous observations that sympatric *Atta* congeners segregate locally along structural and edaphic axes [34,36]. The greater within-species dispersion in *A. cephalotes* (Table 1; Fig 2C) is consistent with a broader realized niche and a generalist ecological strategy capable of exploiting a broader range of environmental conditions [17,43]. Conversely, the lower within-species dispersion in *A. laevigata* suggests ecological specialization.

*Atta* species are traditionally characterized as functional generalist herbivores because of their broad foraging habits and regional distribution [38,39,55]. However, this macro-scale classification often masks significant niche partitioning and specialization at the microhabitat level [35,36]. Our results indicate that while *A. cephalotes* closely matches the generalist archetype [56], *A. laevigata* exhibits greater specificity for multiple features of its microhabitat (Fig 2A). A near 3-fold higher dispersion for canopy openness in *A. cephalotes* compared to *A. laevigata*, coupled with higher SD and IQR values for soil pH (Table 1, S4 Table), indicates that while the former maintains a high capacity to thrive in a variety of microhabitats [42,43,57], the latter remains restricted to a narrower range of environmental axes (Fig 2C). Importantly, both canopy openness and soil conditions are directly modified by land-use change [15,17]. Some temperate studies suggest soil pH stability across land-use types [58]. However, evidence from the Neotropics indicates that forest-to-pasture conversion significantly increases soil pH and nutrient availability [59] and cattle urine deposition further intensifies this effect [60]. Such localized increases in pH fall within the relatively narrow edaphic range occupied by *A. laevigata* [40]. In contrast, the broader edaphic range of *A. cephalotes* may facilitate its occurrence in forest edges and other partially shaded habitats, where soil conditions are more heterogeneous [61].

The architectural and thermal profiles of the colonies suggest that a conserved nest architecture can accommodate distinct microclimatic strategies. On the one hand, mound shape ratios remained comparable across species suggesting a conserved architectural blueprint associated with efficient gas exchange across several species within the *Atta* genus [62–64]. Variation in mound shape ratios (Table 1) suggests flexibility in mound geometry, as previously suggested for colonies growing around roots or sloping terrain [40]. The reduced mound area in *A. cephalotes* may reflect spatial constraints imposed by dense woody vegetation [36]. On the other hand, the resulting internal environments diverged sharply. *A. laevigata* maintained a significantly higher nest temperatures, coupled with its occupation of highly exposed sites (∼90% canopy openness). Together, these findings suggest a strategy in which solar exposure contributes to maintaining favourable nest temperatures in open environments (S2 Fig; S3 Table) [43,62]. In contrast, the lower and more variable internal temperatures of *A. cephalotes* reflect its affinity for shaded micro-refuges (Fig 2B; S6 Table). This pattern is consistent with previous studies showing that colonies can modify local forest structure and microclimatic conditions [65]. Nest interiors in *A. laevigata* remained, on average, more than 10 °C cooler than the surrounding air, compared with approximately 7 °C in *A. cephalotes* (Table 1; S2 Fig). Although ambient temperatures varied with sampling time, this larger temperature difference indicates that *A. laevigata* maintained stronger thermal buffering under the more exposed conditions. Such buffering is likely an advantage in cleared pastures where solar exposure is maximal [15,40]. While sampling location denoted a significant influence on these variables (Table 1), the stronger influence of species identity in canopy openness and thermal regulation is consistent with distinct micro-environmental preferences [34,35] that contribute to niche partitioning.

Soil pH differed significantly between *A. cephalotes* and *A. laevigata* sampling sites (Table 1). Ant biological activity, such as organic waste accumulation, has often been proposed as a driver of localized soil modification [32]. However, our regression analyses did not support direct colony-mediated modification of surface soil pH, as neither mound area nor mound shape predicted pH variation (see *S1 Supplementary Text*). Instead, the separation along PC2 (Fig 2A; S5 Table) is more consistent with underlying differences in soil conditions and land-use history than with direct soil engineering by the colonies. The relatively higher and more stable pH values observed in *A. laevigata* sites is consistent with its strong affinity for cattle pastures, where soil chemistry is frequently modified by livestock urine and agricultural inputs [59,60]. Conversely, the lower and more variable pH levels in *A. cephalotes* sites (4.6 vs. 4.8; S3 Table, S1C Fig) align with the organic-rich profiles typical of forest-edge soils in this region [61]. Although the mean difference between *A. cephalotes* and *A. laevigata* may appear numerically small, the logarithmic nature of the pH scale means this represents a substantial difference in hydrogen ion concentration. Such a shift may influence the symbiotic fungus, *Leucoagaricus gongylophorus* [12]. Notably, the pH values observed for both species align closely with the known physiological optima for *L. gongylophorus*; fungus gardens are typically maintained at a pH of approximately 5.0 to optimize growth and suppress the proliferation of competitor microbes [66]. Although *A. laevigata* occupies a wide range of open anthropogenic habitats [40], it does so within a comparatively narrow range of soil pH (Table 1; Fig 2C). Whether this narrow range reflects a deliberate colonization preference or is a secondary consequence of its strict requirement for open-canopy environments remains unresolved.

Vegetation associations differed markedly between the two species (Fig 2B; S6 Table). *A. cephalotes* exhibited a high affinity for all three vegetation categories (trees, bushes, and grasses). This association with multiple vegetation strata reflects the species’ ability to occupy diverse microhabitats [38,39], likely contributing to its success in heterogeneous forest edges and urban landscapes [42]. In contrast, *A. laevigata* demonstrated a marked association with grass-dominated environments. *A. laevigata* is considered a mixed dicot–grass cutter rather than a strict grass specialist, yet in our study it showed a strong association with grass-dominated habitats, consistent with previous evidence linking shifts toward grass exploitation with the expansion of open landscapes [35]. Together, these patterns indicate that the two species partition the landscape not only along microclimatic and edaphic gradients but also according to vegetation structure (Fig. 2A). This vegetation segregation further supports the narrower realized niche of *A. laevigata* inferred from the multivariate analyses.

The differences in realized niche breadth suggest that *A. cephalotes* and *A. laevigata* may respond differently to land-use change. For example, as forests are transformed into open landscapes, environmental conditions increasingly resemble those occupied by *A. laevigata*. This interpretation is consistent with previous observations showing significantly higher colony survival and growth of *A. laevigata* in man-made clearings than in undisturbed vegetation [40]. While specialists often face range contractions [67], the loss of primary forest habitat appears to be pushing *A. cephalotes* into urban mosaics [42,68], where soils are often chemically heterogeneous because of anthropogenic disturbance [69,70]. Furthermore, urban refuges like residential gardens provide a light-and-shadow matrix in heat islands that resemble the natural forest edge (Fig 2) [42]. This relationship between environmental tolerance and habitat occupancy aligns with a general ecological pattern in which niche breadth is a key predictor of geographical range and species’ vulnerability to environmental change [17,67,71–73]. Our findings are also consistent with recent studies in butterflies and terrestrial snails showing that broader niche breadths are associated with greater persistence across human-modified landscapes, whereas species occupying narrower niche space are more vulnerable to land-use change [18,21,28]. We complement such recent studies conducted across broader geographic scales by demonstrating that the relationship between niche breadth and persistence is also evident when realized niche breadth is quantified directly from fine-scale environmental gradients measured in the field.

Beyond the environmental gradients that we evaluated here, predation may also contribute to the contrasting persistence of both species in human-modified landscapes. Leaf-cutting ants are attacked by a variety of natural enemies, including phorid flies, armadillos, beetles, birds, and spiders, which can influence colony establishment, foraging activity, and colony survival [74,75]. However, little is known about whether changes in predator communities across forests, pastures, and urban environments differentially affect *A. cephalotes* and *A. laevigata*. An additional anthropogenic pressure may arise from the traditional harvest of reproductive females of *A. laevigata* for human consumption in the Santander region, whereas comparable exploitation of *A. cephalotes* is disproportionately lower [76]. Since the demographic consequences of this harvest remain unknown, both natural predation and human harvesting deserve further attention as potential ecological filters acting alongside realized niche breadth in shaping species persistence under land-use change.

Differences between species also extended to mound architecture. Colonies of *A. cephalotes* produced overall smaller and less variable mounds than *A. laevigata* (Table 1; S4 Table). This pattern probably reflects structural constraints imposed by tree roots [36]. In urban environments, where space is often limited, the same characteristic could facilitate colony persistence. Artificial lighting may further contribute to differences in urban colonization. Because *A. cephalotes* performs nocturnal nuptial flights [77], its alates may be more susceptible to light-induced disruption of orientation than the diurnal flights of *A. laevigata* [78]. Similar disruption of nocturnal nuptial flights by light pollution has been reported in *A. texana* [78], a pattern consistent with disruption of the dorsal-light response described for flying insects [79–81]. Together, these observations suggest that different forms of human-modified landscapes may favour different ecological strategies. Consistent with the contrasting realized niche breadths documented here, *A. laevigata* may be favoured by the expansion of agropastoral landscapes, whereas the broader environmental tolerance of *A. cephalotes* may facilitate persistence in heterogeneous urban environments.

Field observations across the study area complemented the quantitative analyses of realized niche breadth (S3-S5 Figs). At the northernmost site (Choro Bajo), both species coexisted under comparatively warmer and drier regional climatic conditions, characterized by higher February temperatures, substantially lower annual precipitation, and relatively high bare-soil evaporation compared with most localities farther south. Conversely, at the southernmost sites (La Victoria and El Raizal) characterized by lower temperatures and generally higher annual precipitation than Choro Bajo, only *A. cephalotes* was present. The absence of *A. laevigata* at these sites, despite available open pasture, suggests its distribution is constrained by regional climatic conditions beyond those occupied elsewhere in the study area. Lower temperatures combined with greater precipitation are therefore plausible environmental factors associated with the absence of *A. laevigata* from this region. While *A. cephalotes* appears capable of colonizing these humid, cold enclaves, the narrower realized niche of *A. laevigata* likely restricts its distribution to warmer, better-drained landscapes [34,40]. Additional field observations also suggested that mound architecture may vary toward the climatic limits of the species’ distribution. We noted shifts in mound architecture at the arid northern limit, such as apparent reductions in entrance hole diameters in *A. laevigata* [62,63]. Representative photographs were archived in the ground-level photographic repository (*S2 Supplementary Dataset*) for future comparisons. While we did not quantify these observations, they may motivate future investigation into potential behavioural responses under high desiccation risk at the species’ range margins [43]. Together, these qualitative observations complement the quantitative analyses by suggesting that *A. laevigata* is constrained by a relatively narrow set of environmental conditions at a regional and local scale, whereas *A. cephalotes* occupies a broader range of climates and microhabitats.

We also identified several methodological aspects that deserve further attention. First, some sampling locations contained substantially more colonies of one species than the other because of their natural distribution across the landscape. Future designs should aim for a more homogeneous species distribution across sites to control for potential site-specific biases. Second, while we maintained a minimum distance of 100 meters between colonies to ensure independence, the future integration of precise GPS coordinates would allow for a more granular analysis of micro-climatic gradients and spatial autocorrelation [38,39]. Third, although the variables that we selected for this study (pH, canopy openness, and mound architecture) captured key dimensions of the realized niche, they represent only a subset of a multivariate space [12]. To further refine our understanding of niche breadth, future research should incorporate additional environmental and biological axes, such as soil moisture, mound entrance dimensions, and specific foraging preferences. Fourth, our inference that differences in realized niche breadth contribute to contrasting persistence in human-modified landscapes is based on direct measurements of niche breadth combined with previously reported presence of *A. cephalotes* in urban areas. Future studies should evaluate colony survival, growth, and persistence across human-modified environments within the same study area to directly test this hypothesis. Finally, as these ecosystem engineers exist in an obligate symbiosis with the fungus *L. gongylophorus* [34,36], focusing solely on the ant provides an incomplete picture. Because colony performance depends on both partners, future work should evaluate how the responses of both the ants and *L. gongylophorus* to environmental conditions contribute to differences in realized niche breadth.

Land-use change in the Anthropocene is reshaping species distributions across natural and human-modified landscapes [15]. As our analyses suggest, agropastoral expansion could favour the open-canopy specialist *A. laevigata*, while simultaneously displacing *A. cephalotes* from its typical woody habitats. This shift involves more than species turnover; it may alter nutrient cycling and other ecosystem processes within shaded forest patches. While *A. cephalotes* nests are known to create localized nutrient-depleted zones [32] and significantly elevate CO_2_ emissions [82], they also drive essential forest heterogeneity by altering canopy structure and light infiltration [65]. The decline of this engineer may trigger ecological cascades in forest remnants [36,39]. At the same time, the increasing occurrence of *A. cephalotes* in urban environments and the expansion of *A. laevigata* across agricultural landscapes may increase the frequency of human–ant interactions, potentially amplifying their impact as pests [42,55]. Understanding the environmental filters associated with land-use change is therefore important for improving the management of human-modified landscapes by helping predict where *A. laevigata* is likely to expand and where *A. cephalotes* may increase occurrence in urban areas.

## Conclusions

We investigated whether the leaf-cutter ants *A. cephalotes* and *A. laevigata* in an area of the Colombian Andes differed in realized niche breadth across fine-scale environmental gradients, and whether those patterns are consistent with ecological strategies of generalism and specialization. Such strategies may shape species response to the ecological filter of ongoing land-use change. *A. cephalotes* presented a broader niche breadth for canopy openness and soil pH, along with a high association with woody-type vegetation. In contrast, *A. laevigata* showed a narrower niche for these environmental variables and an affinity for grasses in open areas, with a larger heat buffering capacity. Mound architecture characterization revealed a comparable elliptical shape, while mounds of *A. laevigata* were significantly larger than those of *A. cephalotes*. Our results support the hypothesis of generalism in *A. cephalotes* capable of occupying a broader range of microhabitats at local and regional scale, while also suggesting specialization of *A. laevigata*, better suited for cleared agro-pastoral landscapes. The findings suggest that a broader realized niche breadth is consistent with the documented occurrence of *A. cephalotes* in urban areas. This study contributes to the understanding of species responses to the ecological filter created by land-use change in view of the growing demand for agricultural lands and may support the management of human-modified landscapes.

## Data availability

All relevant data are fully available without restriction. The original environmental measurements for the 114 colonies analysed in this study are provided in the Supporting Information (S2 Table). Ground-level photographs documenting colony mound architecture are publicly archived in Zenodo (https://doi.org/10.5281/zenodo.20342937). The hemispherical photographs used to estimate canopy openness are publicly archived in Zenodo (https://doi.org/10.5281/zenodo.20344008). The ERA5-Land reanalysis data used for the regional climatic characterization are archived in Zenodo (https://doi.org/10.5281/zenodo.21877589). The Python scripts and Jupyter notebooks used for image processing, statistical analyses, climatic data downloading and processing, and figure generation are archived in Zenodo (https://doi.org/10.5281/zenodo.21886892), with the corresponding development repository available on GitHub.

## Supporting information

S1 Dataset. Characterization of canopy openness across sampled colonies via hemispherical photography

S2 Table. Comprehensive raw dataset of architectural, edaphic, and micro-climatic variables for Atta colonies

## Acknowledgments

I am profoundly grateful to my parents, María Cristina Castillo Sandoval and Víctor Ángel García Aguillón, whose invaluable support made this research possible. Their endurance and dedication during the fieldwork were essential for the successful collection of the colony samples. Furthermore, our endless discussions regarding the natural history and behaviour of these ecosystem engineers deeply shaped the conceptualization of this study; this manuscript is a testament to their lifelong support of my academic journey. I am also sincerely grateful to my sister, Francy Alejandra García Castillo, for generously providing measurement equipment essential for our field data collection. I sincerely thank Alejandro Mejía for his invaluable local knowledge and field assistance in locating *Atta* colonies in Grima Baja and for providing information that guided us to the sampling area in Choro Bajo. I also thank Rosina Soler for her expert professional advice in curating the environmental variable matrix during the fieldwork planning phase, as well as for her insightful feedback during the elaboration of this manuscript.

## Supporting information

**S1 Text. Disentangling the drivers of Principal Component 2 (PC2).** Additional analysis examining the relative contribution of environmental variables to PC2.

**S1 Table. Representative geographic coordinates and corresponding visualization codes for the study locations.** For each of the 12 sampled sites, a single representative GPS coordinate (decimal degrees, WGS 84 datum) is provided. The standardized two-letter abbreviation codes (Two-letter code) are explicitly defined here to facilitate spatial cross-referencing and improve graphical visualization in the box-plot distributions (S1 Fig).

**S2 Table. Comprehensive raw dataset of architectural, edaphic, and micro-climatic variables for *Atta* colonies.** This external CSV dataset contains the microhabitat profiles and structural measurements recorded for 117 active sympatric colonies across a latitudinal and elevational transect in Santander, Colombia. Species identity is specified under the *Species* column, which comprises *A. cephalotes* (n = 56), *A. laevigata* (n = 58), and three colonies designated as U (n = 3), denoting individuals of ambiguous micro-habitat x morphology that could not be confidently classified to the species level in the field; these three unclassified colonies were included in the raw metadata for transparency but were systematically excluded from downstream species-specific statistical analyses. Column headers are defined as follows: *Num_Col* represents the unique numerical identifier assigned to each sampled nest; *Location* denotes the specific geographic site of collection (spanning 12 distinct sampling localities); *Date_Fieldwork* indicates the exact date of data collection; and *Species* identifies the target organism. Structural parameters include mound *Length* (cm), *Width* (cm), and *Number_Active_Holes*. Although *Number_Active_Holes* was ultimately excluded from downstream statistical modelling. Micro-climatic and edaphic axes consist of *Out_Temp* (°C; ambient external temperature), *In_Temp* (°C; internal mound temperature at ∼10 cm depth), and *Soil_pH*. Canopy structural metrics are represented by *Time_Photo* (the exact timestamp of canopy image capture) and *Photo_File* (the high-resolution hemispherical photograph filename used to calculate canopy openness). In all main text analyses and figures, the calculated values derived from these hemispherical photographs are referred to as Canopy Openness (%), corresponding to the raw *Light_Percentage* values presented herein. Environmental background states are qualified under *Sky* (visual assessment of cloud cover parameters during sampling).

**S3 Table. Descriptive statistics and Shapiro–Wilk normality tests for ecological variables measured in *A. cephalotes* (C) and *A. laevigata* (L).** Means, standard deviations (SD), and *p*-values are presented for each species. Variables were classified as normally distributed when both species showed *p* > 0.05. The departure from normality observed for internal temperature in *A. laevigata* is likely attributable to the truncated sampling period (ending at approximately 14:00), which captured only the warming phase of the daily thermal cycle.

**S4 Table. Dispersion metrics for architectural and environmental variables.** Intraspecific variation for *A. cephalotes* (C) and *A. laevigata* (L) is summarized using Standard Deviation (SD) and Interquartile Range (IQR). Larger values indicate greater within-species dispersion for the corresponding variable. Mound Area is reported in m² to facilitate ecological interpretation.

**S5 Table. Percentage contribution of ecological and structural variables to the first two axes of the Principal Component Analysis (PCA).** Variables are ordered by their contribution to PC1. High contributions on PC1 reflect local vegetation density and canopy architecture, whereas high loadings on PC2 differentiate colony dimensions (mound area and shape ratio) and chemical soil properties (pH).

**S6 Table. Indicator Value (IndVal) analysis of vegetation categories (Grasses, Bushes, and Trees) within a 10 m radius of *A. cephalotes* and *A. laevigata mounds*.** Because vegetation categories were not mutually exclusive, indices were calculated based on the independent presence of each structural component. Specificity (A) indicates the exclusivity of a species to a vegetation category, Fidelity (B) indicates the frequency of the category’s occurrence with the species, and the IndVal index represents the combined indicative strength.

**S7 Table. Generalized Linear Model (GLM) regression results for the additive effects of colony architecture and species identity on soil pH.** The model was fitted using a Gaussian error distribution and an identity link function (*N* = 111 colonies, reflecting the subset of the 114 total downstream colonies with complete soil pH records). Geographic localities were incorporated as categorical dummy variables to serve as spatial blocks, isolating landscape-level edaphic baselines. The reference baseline (Intercept) reflects a baseline geographic control site. Coefficients (β), standard errors (SE), Wald *z*-scores, and corresponding *p*-values are provided for each predictor variable.

**S8 Table. Generalized Linear Model (GLM) regression results evaluating the conditional effects and species-specific interactions of colony traits on soil pH.** The model utilizes a Gaussian error distribution and an identity link function (*N* = 111 colonies, reflecting the subset of the 114 total downstream colonies with complete soil pH records) to test the biological hypothesis that soil engineering capacity varies as a function of species identity (represented by the multiplicative interaction term Mound_Area:Species[T.L]). Geographic localities are controlled via spatial blocking parameters. Coefficients (β), standard errors (SE), Wald *z*-scores, and *p*-values are reported for all fixed effects and spatial block contrasts.

**S1 Fig. Multivariate assessment of mound architecture, edaphic properties, and thermal regulation across sampling locations.** Box-and-whisker plots **(A–E)** illustrate the intraspecific and interspecific variation for A. *cephalotes* (green) and *A. laevigata* (orange) across twelve distinct geographic sites. (A) *Mound Area*, presented as Log_10_-transformed values (cm^2^) to normalize the highly skewed distribution of colony sizes. (B) *Shape Ratio*, defined as the degree of surface symmetry. **(C) Soil pH**, representing the chemical conditions of the mound substrate. (D) *Light Percentage*, indicating the level of canopy openness at the nest site. (E) *Temperature Difference*, representing the thermal gradient between the internal colony core at ∼10 cm and the external environment (°C). Shaded boxes denote the interquartile range (IQR), horizontal lines represent the median, and whiskers extend to 1.5 times the IQR. Two-letter codes at the base of each distribution correspond to specific sampling locations (e.g., MF: Montefrio, NZ: Nemizaque, etc.; see S1 Table for full location metadata).

**S2 Fig. Diel thermal profiles of *Atta* colony interiors.** Points represent individual temperature measurements (n = 114) recorded at approximately 10 cm depth for *A. cephalotes* (green circles) and *A. laevigata* (orange crosses). Trend lines represent a second-order polynomial regression (*R*^2^ fit), with shaded ribbons indicating the 95% confidence intervals. *A. laevigata* colonies exhibited significantly higher internal temperatures compared to *A. cephalotes* (PERMANOVA test: ΔT = 1.2°C, *p* = 0.001; S3 and S4 Tables). The widening of the confidence interval for *A. laevigata* after 14:00 reflects the increased statistical uncertainty at the upper bound of the sampling window for this species.

**S3 Fig. Regional average February temperature across the study area in Santander, Colombia, derived from ERA5-Land reanalysis data.** Panels **(A–C)** show the average 2 m air temperature (°C) for February 2024, 2025, and 2026, respectively, using a shared colour scale ranging from 16.74°C (yellow; lower temperatures) to 22.92°C (deep red; higher temperatures). All climatic heatmaps are georeferenced and overlaid on an OpenStreetMap basemap of the sampling region. Black points indicate the fieldwork locations where *Atta* colonies were surveyed, with location names provided for geographic reference across the latitudinal and elevational gradient of the study area. Base map © OpenStreetMap contributors. Map generated in Python 3.12 using Matplotlib, GeoPandas, Contextily, xarray, and rioxarray.

**S4 Fig. Regional total precipitation across the study area in Santander, Colombia, derived from ERA5-Land reanalysis data.** Panels **(A–B)** show the total annual accumulated precipitation (m) in 2024 and 2025, respectively, using a shared colour scale ranging from approximately 1.6 m (yellow; lower precipitation) to 2.5 m (deep blue; higher precipitation). Panel **(C)** shows cumulative precipitation from January to July 2026 and uses a separate colour scale because it represents a partial year. All climatic heatmaps are georeferenced and overlaid on an OpenStreetMap basemap of the sampling region. Black points indicate the fieldwork locations where *Atta* colonies were surveyed, with location names provided for geographic reference across the latitudinal and elevational gradient of the study area. Base map © OpenStreetMap contributors. Map generated in Python 3.12 using Matplotlib, GeoPandas, Contextily, xarray, and rioxarray.

**S5 Fig. Regional total bare-soil evaporation across the study area in Santander, Colombia, derived from ERA5-Land reanalysis data.** Panels **(A–B)** show the total annual accumulated bare-soil evaporation (mm) in 2024 and 2025, respectively, using a shared colour scale ranging from approximately 267 mm (yellow; lower evaporation) to 377 mm (deep blue; higher evaporation). Panel **(C)** shows cumulative bare-soil evaporation from January to July 2026 and uses a separate colour scale because it represents a partial year. All climatic heatmaps are georeferenced and overlaid on an OpenStreetMap basemap of the sampling region. Black points indicate the fieldwork locations where *Atta* colonies were surveyed, with location names provided for geographic reference across the latitudinal and elevational gradient of the study area. Base map © OpenStreetMap contributors. Map generated in Python 3.12 using Matplotlib, GeoPandas, Contextily, xarray, and rioxarray.

**S1 Dataset. Characterization of canopy openness across sampled colonies via hemispherical photography.** This PDF document compile the complete photographic registry for all 117 surveyed *Atta* colonies along the regional transect. Each row presents a pairwise comparison containing: (left) the raw hemispherical photograph converted to greyscale, and (right) the binarized silhouette version resulting from the interactive threshold masking process. Each pair is labelled with its unique image identifier (Photo_File), the selected binarization algorithm context (Sky: S [sunny], C [cloudy] or P [partially cloudy]), the specific mathematically applied Gamma correction factor, and the final estimated light percentage calculated over the mound. This light percentage serves as our proxy for canopy openness used in all downstream niche breadth and partitioning multivariate analyses.

**S2 Dataset. Ground-level photographic registry of colony mound architectures.** This compressed file (Supplementary_Dataset_S2.zip) contains the complete repository of fieldwork photographs of mounds for the 117 surveyed *Atta* colonies. The dataset includes wide-angle ground-level photographs of each distinct nest mound, captured utilizing the same hemispherical fish-eye lens system described for the canopy sub-canopy characterization in the *Materials and Methods* section. Additionally, some standard-format (non-distorted) landscape photographs are provided. While these visual records were not deployed as quantitative variables in downstream statistical modelling, they are archived here to provide transparent empirical evidence and an environmental record of the regional fieldwork. The .zip file was deposited at Zenodo and it is publicly available at https://doi.org/10.5281/zenodo.20342937.

