## Supplemental Figures, Tables, and Text for "Generalism vs. specialization: Does niche breadth influence species responses to anthropogenic land-use change in Neotropical leaf-cutter ants?"

[*] Corresponding author

### Supporting information

#### Supporting text

**S1 Text. Disentangling the drivers of Principal Component 2 (PC2)**

To determine whether the high loadings of mound architecture (*Mound Area*, *Shape Ratio*) and *Soil pH* onto PC2 (Fig 2A; S6 Table) represented a causal ecosystem engineering mechanism (i.e., larger nests driving local soil modification), we constructed Generalized Linear Models (GLMs) using a Gaussian error distribution.

*Model 1. Additive effects model (without interaction):*

The first model evaluated the independent, direct effects of *Mound Area*, *Shape Ratio*, and *Species* identity on *Soil pH* under the formula:

```
Soil_pH ~ Species + Location + Mound_Area + Shape_Ratio
```

*Model 2. Conditional effects Model (with species interaction):*

To test the biological hypothesis that soil engineering capacity is species-specific (i.e., that the slope of *Soil pH* as a function of *Mound Area* differs between *A. cephalotes* and *A. laevigata*), we introduced a multiplicative interaction term under the formula:

```
Soil_pH ~ Mound_Area * Species + Shape_Ratio + Location
```

After accounting for sampling *Location*, the apparent relationship between colony traits and soil chemistry disappeared. Neither *Mound Area* (*P*= 0.641) nor *Shape Ratio* (*P* = 0.532) predicted *Soil pH* variation (S7 Table). Furthermore, the interaction between *Species* identity and *Mound Area* was non-significant (*P* = 0.136; S8 Table), demonstrating that this lack of architectural effect is consistent across both *Atta* species. Instead, the model's high explanatory power (Pseudo-$R^2$ = 0.66) was driven entirely by spatial blocks (*Location* parameters such as *Location[T.Choro Bajo]*, *Location[T.El Raizal]*, *Location[T.Grima Baja]*, etc.).

We therefore conclude that the association between mound architecture and soil pH observed along PC2 does not reflect direct colony-mediated ecosystem engineering. Instead, it is more consistent with spatial differences among sampling locations, likely reflecting underlying edaphic conditions and land-use history.

### Supporting tables

**S1 Table. Representative geographic coordinates and corresponding visualization codes for the study locations.**

| Location | Two-letter code | Latitude (N) | Longitude (W) |
|---|---|---|---|
| **Agua Buena** | AB | 6.28135 | -073.13047 |
| **Cantera** | CT | 6.17719 | -073.18733 |
| **Choro Bajo** | VN | 6.64588 | -073.16210 |
| **El Raizal** | RZ | 6.14935 | -073.18629 |
| **Grima Baja** | CB | 6.30970 | -073.15544 |
| **La Siberia** | LS | 6.27547 | -073.15166 |
| **La Victoria** | LV | 6.15613 | -073.18887 |
| **Montefrio** | MF | 6.27050 | -073.14447 |
| **Morario** | MO | 6.30960 | -073.13513 |
| **Nemizaque** | NZ | 6.22833 | -073.17700 |
| **Pozo Gallo** | PG | 6.26865 | -073.15557 |
| **Villas** | VI | 6.26149 | -073.16818 |

**S3 Table. Descriptive statistics and Shapiro–Wilk normality tests for ecological variables measured in *A. cephalotes* (C) and *A. laevigata* (L).**

| Ecological Variable | Mean (C) | SD (C) | Mean (L) | SD (L) | *p*-norm (C) | *p*-norm (L) | Assumed Distribution |
|---|---|---|---|---|---|---|---|
| **Mound area ($m^2$)** | 45.015 | 37.452 | 69.059 | 61.899 | < 0.001 | < 0.001 | Non-normal |
| **Shape ratio** | 1.74 | 0.924 | 2.062 | 1.101 | < 0.001 | < 0.001 | Non-normal |
| **Soil pH** | 4.666 | 0.376 | 4.815 | 0.261 | 0.001 | 0.0008 | Non-normal |
| **Canopy openness (%)** | 67.668 | 18.493 | 89.57 | 6.919 | 0.0437 | < 0.001 | Non-normal |
| **Internal temperature ~10 cm (°C)** | 23.982 | 1.384 | 25.145 | 1.543 | 0.2191 | 0.0032 | Non-normal |
| **Thermal regulation capacity (°C)** | -7.07 | 4.388 | -10.32 | 4.7 | 0.2313 | 0.8717 | Normal |

Means, standard deviations (SD), and *p*-values are presented for each species. Variables were classified as normally distributed when both species showed $p > 0.05$. The departure from normality observed for internal temperature in *A. laevigata* is likely attributable to the truncated sampling period (ending at approximately 14:00), which captured only the warming phase of the daily thermal cycle.

**S4 Table. Dispersion metrics for architectural and environmental variables.**

| Variable | SD (C) | SD (L) | IQR (C) | IQR (L) |
|---|---|---|---|---|
| **Canopy openness (%)** | 18.493 | 6.755 | 30.325 | 4.987 |
| **Mound area (m²)** | 37.452 | 61.489 | 46.107 | 70.142 |
| **Shape ratio** | 0.924 | 1.08 | 0.74 | 0.933 |
| **Soil pH** | 0.376 | 0.261 | 0.6 | 0.3 |
| **Thermal regulation capacity (°C)** | 4.388 | 4.7 | 5.15 | 6.05 |

Intraspecific variation for *A. cephalotes* (C) and *A. laevigata* (L) is summarized using Standard Deviation (SD) and Interquartile Range (IQR). Larger values indicate greater within-species dispersion for the corresponding variable. Mound Area is reported in m² to facilitate ecological interpretation.

**S5 Table. Percentage contribution of ecological and structural variables to the first two axes of the Principal Component Analysis (PCA).**

| Variable | PC1 (%) | PC2 (%) |
|---|---|---|
| **Trees-type vegetation** | 28.4 | 0.14 |
| **Bush-type vegetation** | 25.95 | 0.06 |
| **Canopy openness (%)** | 23.7 | 1.3 |
| **Thermal regulation capacity (°C)** | 12.38 | 4.78 |
| **Mound area ($m^2$)** | 5.34 | 30.18 |
| **Grass-type vegetation** | 3.33 | 5.66 |
| **Shape ratio** | 0.47 | 27.8 |
| **Soil pH** | 0.43 | 30.08 |

Variables are ordered by their contribution to PC1. High contributions on PC1 reflect local vegetation density and canopy architecture, whereas high loadings on PC2 differentiate colony dimensions (mound area and shape ratio) and chemical soil properties (pH).

**S6 Table. Indicator Value (IndVal) analysis of vegetation categories (Grasses, Bushes, and Trees) within a 10 m radius of *A. cephalotes* and *A. laevigata* mounds.**

| Species | Vegetation Category | Specificity (A) | Fidelity (B) | IndVal Index |
|---|---|---|---|---|
| ***A. cephalotes*** | Bushes | 0.783 | 0.964 | 0.869 |
| ***A. cephalotes*** | Trees | 0.785 | 0.911 | 0.845 |
| ***A. cephalotes*** | Grasses | 0.482 | 0.946 | 0.675 |
| ***A. laevigata*** | Grasses | 0.518 | 0.983 | 0.714 |
| ***A. laevigata*** | Bushes | 0.217 | 0.259 | 0.237 |
| ***A. laevigata*** | Trees | 0.215 | 0.241 | 0.228 |

Because vegetation categories were not mutually exclusive, indices were calculated based on the independent presence of each structural component. Specificity (A) indicates the exclusivity of a species to a vegetation category, Fidelity (B) indicates the frequency of the category's occurrence with the species, and the IndVal index represents the combined indicative strength.

**S7 Table. Generalized Linear Model (GLM) regression results for the additive effects of colony architecture and species identity on soil pH.**

| | Coef. | Std.Err. | *z* | *p* > \|*z*\| | [0.025 | 0.975] |
|---|---|---|---|---|---|---|
| **Intercept** | 4.6012 | 0.115 | 40.021 | 0 | 4.3759 | 4.8265 |
| **Species[T.L]** | -0.0118 | 0.0669 | -0.177 | 0.8597 | -0.1429 | 0.1193 |
| **Location[T.Cantera]** | -0.2107 | 0.156 | -1.351 | 0.1768 | -0.5165 | 0.095 |
| **Location[T.Choro Bajo]** | 0.3801 | 0.0986 | 3.856 | 0.0001 | 0.1869 | 0.5733 |
| **Location[T.El Raizal]** | -0.6155 | 0.1416 | -4.347 | 0 | -0.893 | -0.338 |
| **Location[T.Grima Baja]** | 0.323 | 0.1047 | 3.086 | 0.002 | 0.1178 | 0.5281 |
| **Location[T.La Siberia]** | 0.2149 | 0.1046 | 2.055 | 0.0399 | 0.0099 | 0.4199 |
| **Location[T.La Victoria]** | -0.2022 | 0.1296 | -1.56 | 0.1187 | -0.4563 | 0.0518 |
| **Location[T.Montefrio]** | -0.0161 | 0.1065 | -0.152 | 0.8795 | -0.2248 | 0.1925 |
| **Location[T.Morario]** | 0.3478 | 0.1445 | 2.407 | 0.0161 | 0.0646 | 0.6311 |
| **Location[T.Nemizaque]** | 0.234 | 0.1017 | 2.302 | 0.0213 | 0.0348 | 0.4333 |
| **Location[T.Pozo Gallo]** | 0.2895 | 0.1239 | 2.337 | 0.0194 | 0.0467 | 0.5324 |
| **Location[T.Villas]** | 0.3018 | 0.1635 | 1.846 | 0.0649 | -0.0186 | 0.6223 |
| **Mound_Area** | 0 | 0 | -0.467 | 0.6405 | 0 | 0 |
| **Shape_Ratio** | 0.0149 | 0.0239 | 0.625 | 0.532 | -0.0319 | 0.0617 |
| Model Fit: No. Obs = 111 \| Df Residuals = 96 \| Pseudo R-squ. (CS) = 0.6501 \| Deviance = 5.4869 | | | | | | |

The model was fitted using a Gaussian error distribution and an identity link function (*N* = 111 colonies, reflecting the subset of the 114 total downstream colonies with complete soil pH records). Geographic localities were incorporated as categorical dummy variables to serve as spatial blocks, isolating landscape-level edaphic baselines. The reference baseline (Intercept) reflects a baseline geographic control site. Coefficients (β), standard errors

(SE), Wald *z*-scores, and corresponding *p*-values are provided for each predictor variable.

**S8 Table. Generalized Linear Model (GLM) regression results evaluating the conditional effects and species-specific interactions of colony traits on soil pH.**

| | Coef. | Std.Err. | *z* | *p* > \|*z*\| | [0.025 | 0.975] |
|---|---|---|---|---|---|---|
| **Intercept** | 4.6301 | 0.1159 | 39.954 | 0 | 4.403 | 4.8572 |
| **Species[T.L]** | -0.1075 | 0.0924 | -1.163 | 0.245 | -0.2886 | 0.0737 |
| **Location[T.Cantera]** | -0.2146 | 0.1551 | -1.384 | 0.1664 | -0.5185 | 0.0893 |
| **Location[T.Choro Bajo]** | 0.4232 | 0.1022 | 4.143 | 0 | 0.223 | 0.6235 |
| **Location[T.El Raizal]** | -0.588 | 0.1419 | -4.143 | 0 | -0.8661 | -0.3098 |
| **Location[T.Grima Baja]** | 0.3565 | 0.1064 | 3.35 | 0.0008 | 0.1479 | 0.565 |
| **Location[T.La Siberia]** | 0.2436 | 0.1057 | 2.305 | 0.0212 | 0.0364 | 0.4508 |
| **Location[T.La Victoria]** | -0.1997 | 0.1288 | -1.55 | 0.1211 | -0.4522 | 0.0528 |
| **Location[T.Montefrio]** | -0.0081 | 0.1059 | -0.077 | 0.9389 | -0.2158 | 0.1995 |
| **Location[T.Morario]** | 0.3569 | 0.1437 | 2.483 | 0.013 | 0.0752 | 0.6387 |
| **Location[T.Nemizaque]** | 0.2803 | 0.1057 | 2.652 | 0.008 | 0.0731 | 0.4874 |
| **Location[T.Pozo Gallo]** | 0.3332 | 0.1266 | 2.633 | 0.0085 | 0.0852 | 0.5813 |
| **Location[T.Villas]** | 0.324 | 0.1632 | 1.986 | 0.0471 | 0.0042 | 0.6438 |
| **Mound_Area** | 0 | 0 | -1.502 | 0.1332 | 0 | 0 |
| **Mound_Area:Species[T.L]** | 0 | 0 | 1.489 | 0.1364 | 0 | 0 |
| **Shape_Ratio** | 0.0162 | 0.0237 | 0.682 | 0.4951 | -0.0303 | 0.0627 |
| Model Fit: No. Obs = 111 \| Df Residuals = 95 \| Pseudo R-squ. (CS) = 0.6620 \| Deviance = 5.3617 | | | | | | |

The model utilizes a Gaussian error distribution and an identity link function (*N* = 111 colonies, reflecting the subset of the 114 total downstream colonies with complete soil pH records) to test the biological hypothesis that soil engineering capacity varies as a function of species identity (represented by the multiplicative interaction term Mound_Area:Species[T.L]). Geographic localities are controlled via spatial blocking parameters. Coefficients (β), standard errors (SE), Wald *z*-scores, and *p*-values are reported for all fixed effects and spatial block contrasts.

### Supporting figures

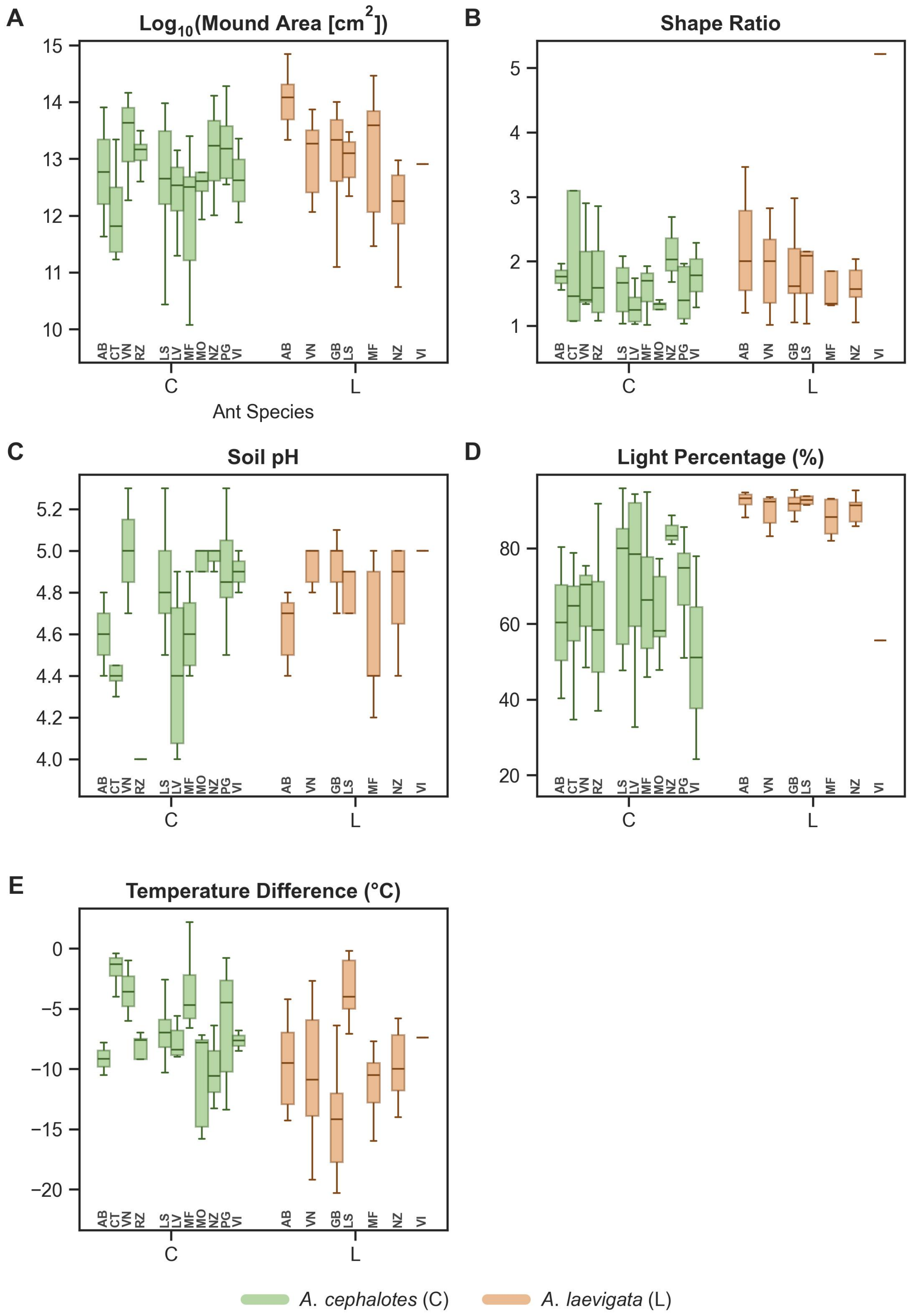


**S1 Fig. Multivariate assessment of mound architecture, edaphic properties, and thermal regulation across sampling locations.** Box-and-whisker plots **(A–E)**

illustrate the intraspecific and interspecific variation for A. *cephalotes* (green) and *A. laevigata* (orange) across twelve distinct geographic sites. **(A)** *Mound Area*, presented as $Log_{10}$-transformed values ($cm^2$) to normalize the highly skewed distribution of colony sizes. **(B)** *Shape Ratio*, defined as the degree of surface symmetry. **(C) Soil pH**, representing the chemical conditions of the mound substrate. **(D)** *Light Percentage*, indicating the level of canopy openness at the nest site. **(E)** *Temperature Difference*, representing the thermal gradient between the internal colony core at ~10 cm and the external environment (°C). Shaded boxes denote the interquartile range (IQR), horizontal lines represent the median, and whiskers extend to 1.5 times the IQR. Two-letter codes at the base of each distribution correspond to specific sampling locations (e.g., MF: Montefrio, NZ: Nemizaque, etc.; see S1 Table for full location metadata).

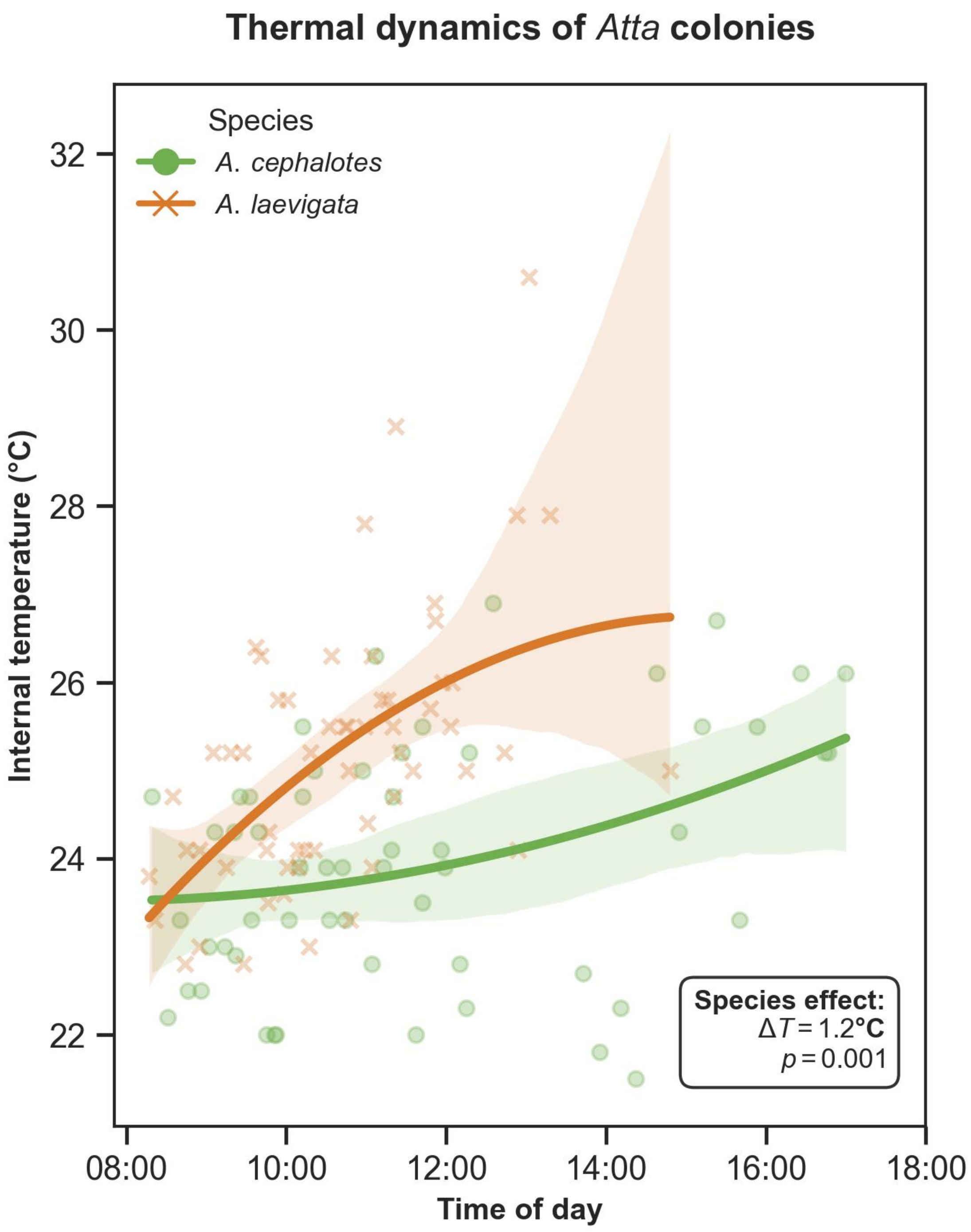


**S2 Fig. Diel thermal profiles of *Atta* colony interiors.** Points represent individual temperature measurements (n = 114) recorded at approximately 10 cm depth for *A. cephalotes* (green circles) and *A. laevigata* (orange crosses). Trend lines represent a second-order polynomial regression ($R^2$ fit), with shaded ribbons indicating the 95% confidence intervals. *A. laevigata* colonies exhibited significantly higher internal temperatures compared to *A. cephalotes* (PERMANOVA test: ΔT = 1.2°C, *p* = 0.001; S3 and S4 Tables). The widening of the confidence interval for *A. laevigata* after 14:00

reflects the increased statistical uncertainty at the upper bound of the sampling window for this species.

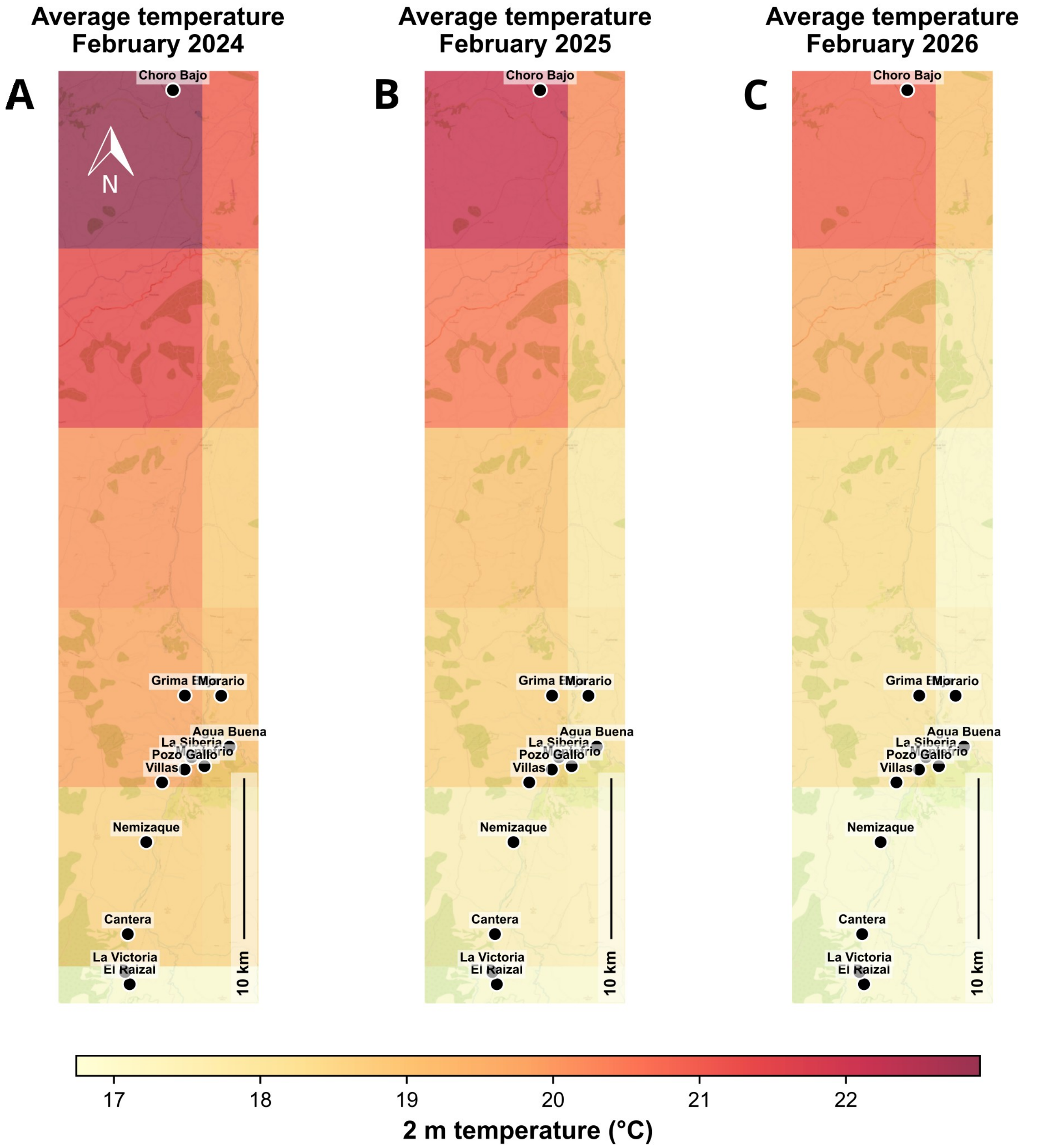


**S3 Fig. Regional average February temperature across the study area in Santander, Colombia, derived from ERA5-Land reanalysis data.** Panels **(A–C)** show the average 2 m air temperature (°C) for February 2024, 2025, and 2026, respectively, using a shared colour scale ranging from 16.74°C (yellow; lower temperatures) to 22.92°C (deep red; higher temperatures). All climatic heatmaps are georeferenced and overlaid on an OpenStreetMap basemap of the sampling region. Black points indicate the fieldwork locations where *Atta* colonies were surveyed, with location names provided for geographic reference across the latitudinal and elevational gradient of the study area. Base map © OpenStreetMap contributors. Map generated in Python 3.12 using Matplotlib, GeoPandas, Contextily, xarray, and rioxarray.

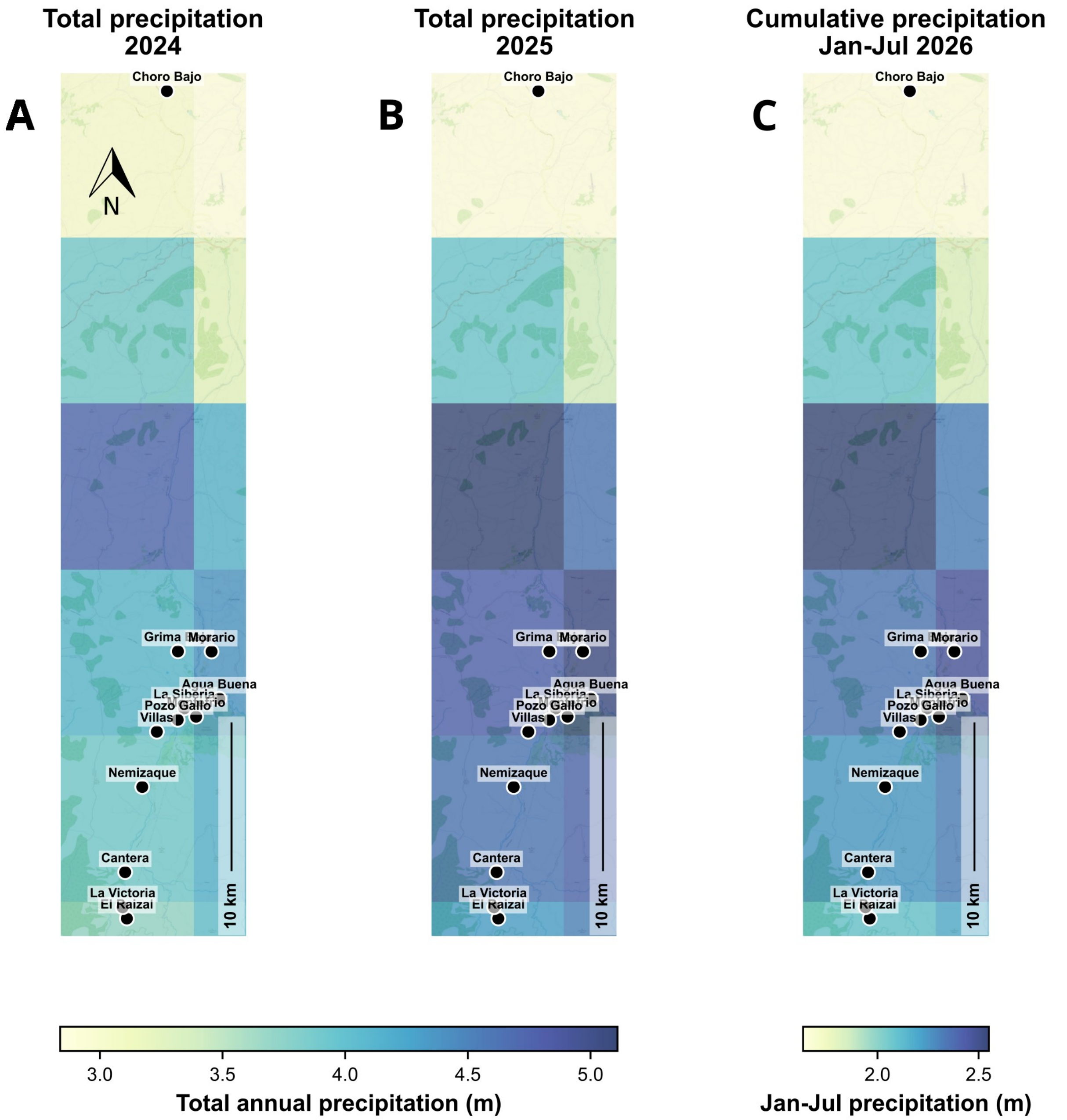


**S4 Fig. Regional total precipitation across the study area in Santander, Colombia, derived from ERA5-Land reanalysis data.** Panels **(A–B)** show the total annual accumulated precipitation (m) in 2024 and 2025, respectively, using a shared colour scale ranging from approximately 1.6 m (yellow; lower precipitation) to 2.5 m (deep blue; higher precipitation). Panel **(C)** shows cumulative precipitation from January to July 2026 and uses a separate colour scale because it represents a partial year. All climatic heatmaps are georeferenced and overlaid on an OpenStreetMap basemap of the sampling region. Black points indicate the fieldwork locations where *Atta* colonies were surveyed, with location names provided for geographic reference across the latitudinal and elevational gradient of the study area. Base map ©

OpenStreetMap contributors. Map generated in Python 3.12 using Matplotlib, GeoPandas, Contextily, xarray, and rioxarray.

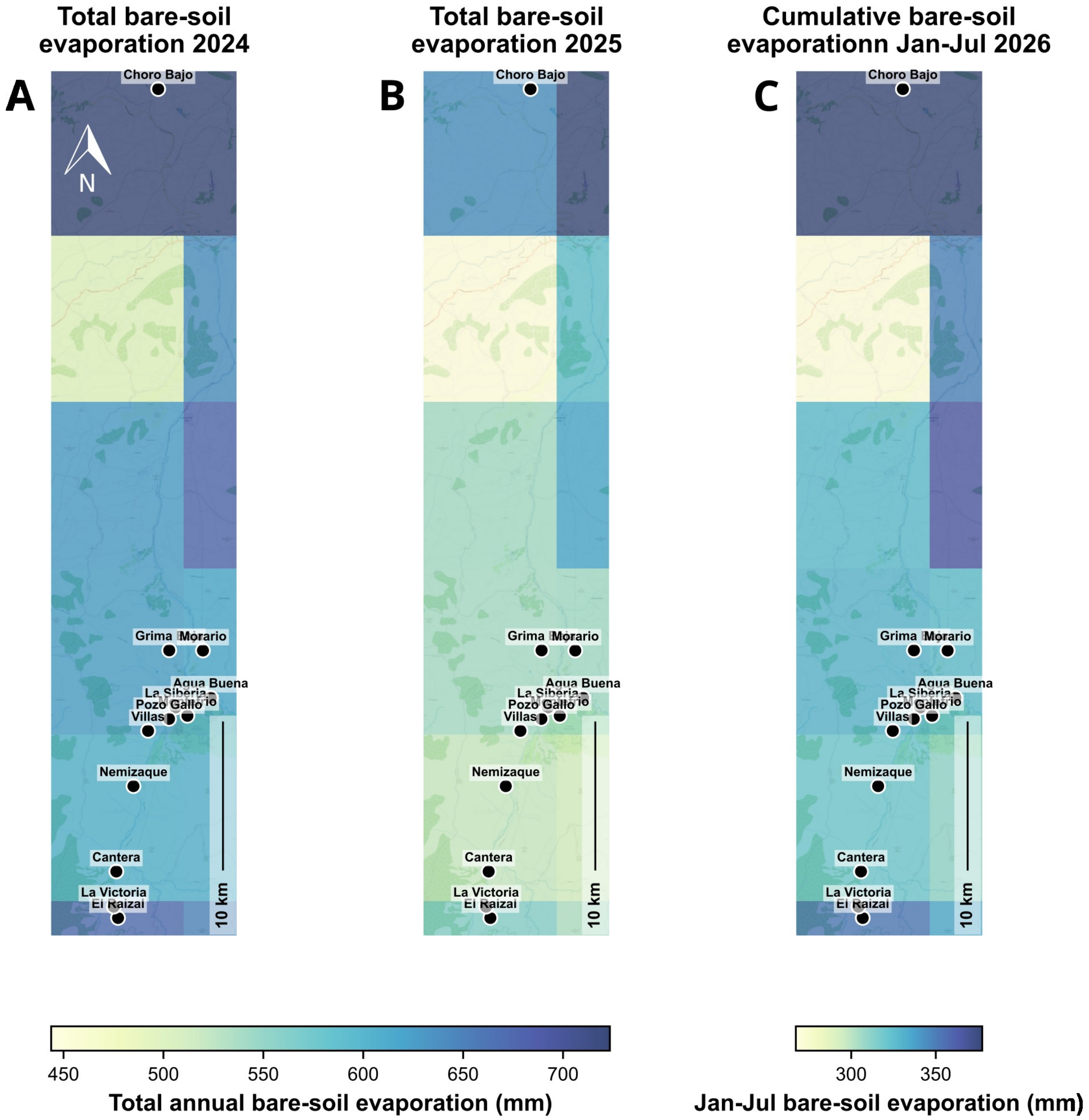


**S5 Fig. Regional total bare-soil evaporation across the study area in Santander, Colombia, derived from ERA5-Land reanalysis data.** Panels **(A–B)** show the total annual accumulated bare-soil evaporation (mm) in 2024 and 2025, respectively, using a shared colour scale ranging from approximately 267 mm (yellow; lower evaporation) to 377 mm (deep blue; higher evaporation). Panel **(C)** shows cumulative bare-soil evaporation from January to July 2026 and uses a separate colour scale because it represents a partial year. All climatic heatmaps are georeferenced and overlaid on an OpenStreetMap basemap of the sampling region. Black points indicate the fieldwork locations where *Atta* colonies were surveyed, with location names provided for geographic reference across the latitudinal and elevational gradient of the study area.

Base map © OpenStreetMap contributors. Map generated in Python 3.12 using Matplotlib, GeoPandas, Contextily, xarray, and rioxarray.

### Supporting datasets

**S1 Dataset. Characterization of canopy openness across sampled colonies via hemispherical photography.** This PDF document compile the complete photographic registry for all 117 surveyed *Atta* colonies along the regional transect. Each row presents a pairwise comparison containing: (left) the raw hemispherical photograph converted to greyscale, and (right) the binarized silhouette version resulting from the interactive threshold masking process. Each pair is labelled with its unique image identifier (Photo_File), the selected binarization algorithm context (Sky: S [sunny], C [cloudy] or P [partially cloudy]), the specific mathematically applied Gamma correction factor, and the final estimated light percentage calculated over the mound. This light percentage serves as our proxy for canopy openness used in all downstream niche breadth and partitioning multivariate analyses.

**S2 Dataset. Ground-level photographic registry of colony mound architectures.** This compressed file (Supplementary_Dataset_S2.zip) contains the complete repository of fieldwork photographs of mounds for the 117 surveyed *Atta* colonies. The dataset includes wide-angle ground-level photographs of each distinct nest mound, captured utilizing the same hemispherical fish-eye lens system described for the canopy sub-canopy characterization in the *Materials and Methods* section. Additionally, some standard-format (non-distorted) landscape photographs are provided. While these visual records were not deployed as quantitative variables in downstream statistical modelling, they are archived here to provide transparent empirical evidence and an environmental record of the regional fieldwork. The .zip file was deposited at Zenodo and it is publicly available at https://doi.org/10.5281/zenodo.20342937
