## Supplementary material for "Generalism vs. specialization: Does niche breadth influence species responses to anthropogenic land-use change in Neotropical leaf-cutter ants?": S1 Dataset. Characterization of canopy openness across sampled colonies via hemispherical photography

### Supplementary Dataset S1. Characterization of canopy openness across sampled colonies via hemispherical photography.

*Manuscript title: Plasticity vs. specialization: Does niche breadth predict resilience to Anthropogenic land-use change in ecosystem engineers?*

Author: Garcia Castillo, D. 2026.

**ID: IMG_20260207_081903.jpg | Sky: P (Gamma: 1.5) | Light: 81.15%**

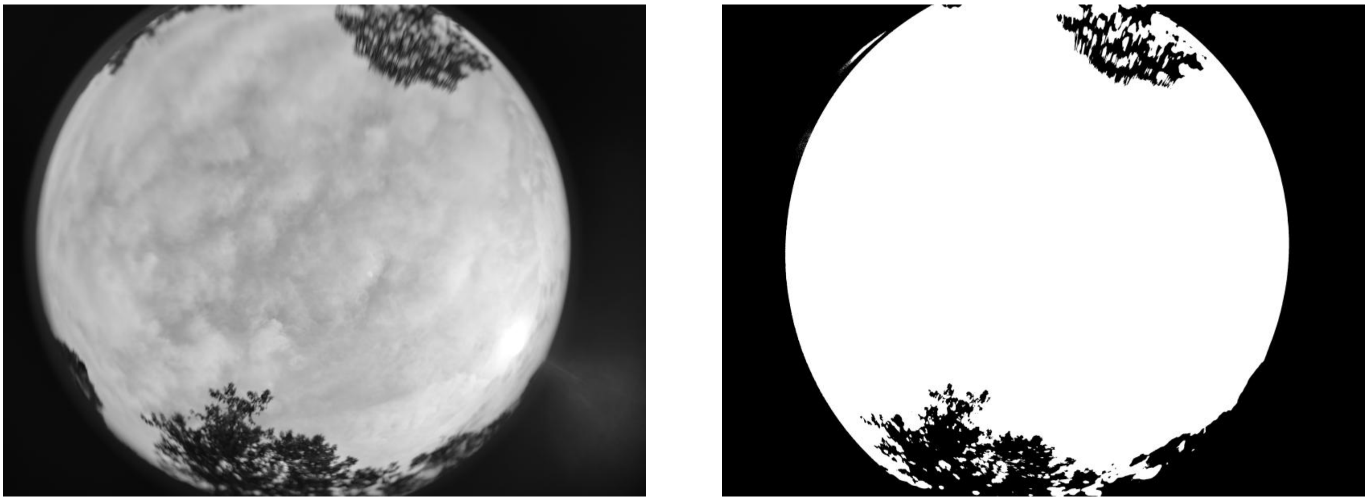

**ID: IMG_20260207_083925.jpg | Sky: P (Gamma: 1.5) | Light: 74.15%**

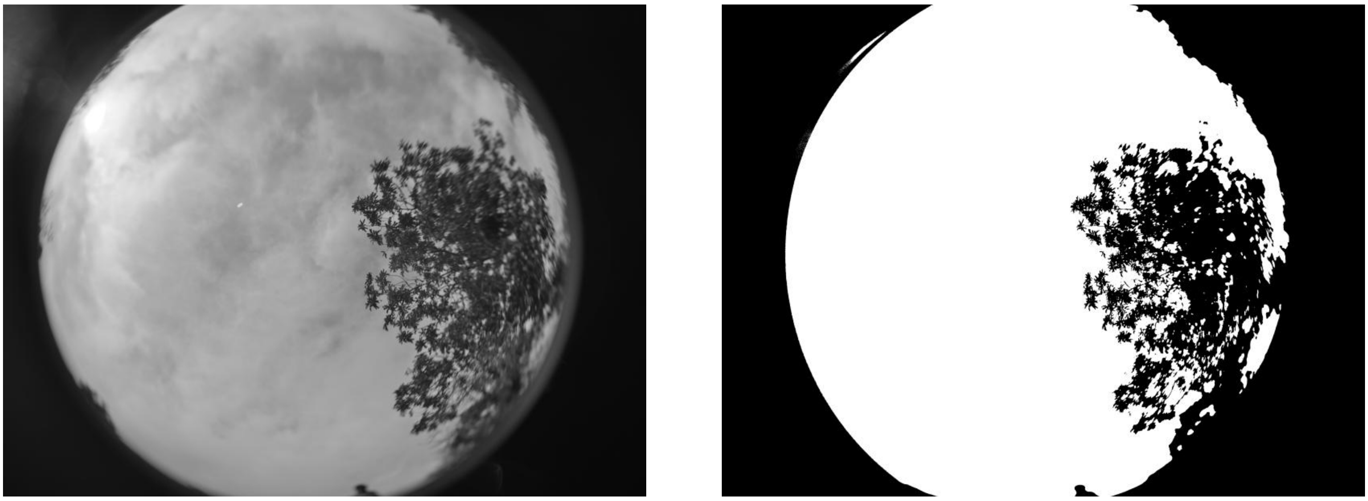

**ID: IMG_20260207_085621.jpg | Sky: P (Gamma: 1.5) | Light: 66.35%**

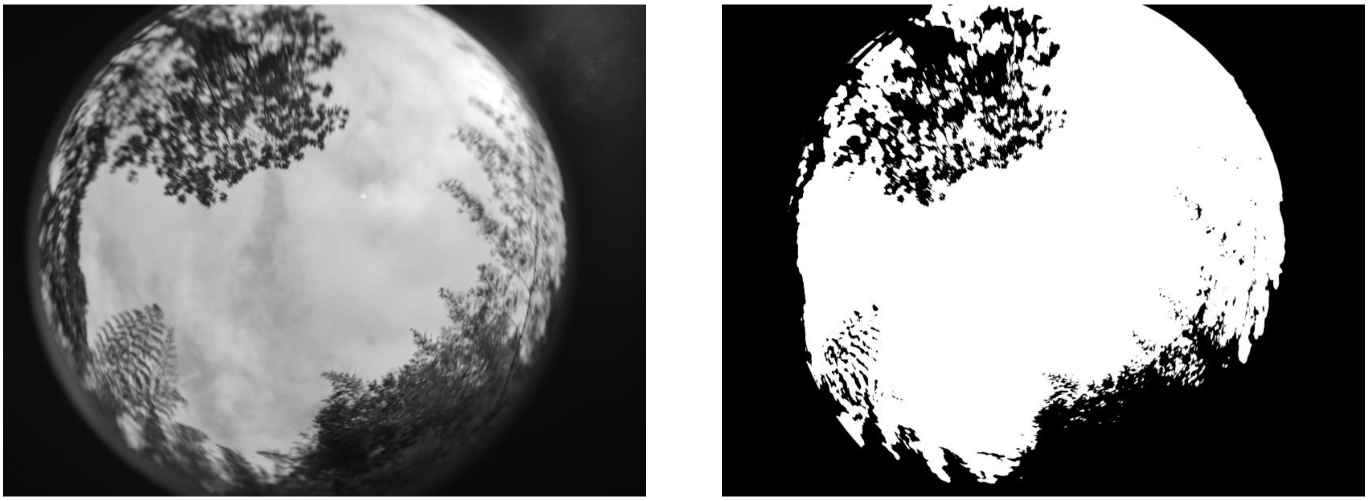

**ID: IMG_20260207_091608.jpg | Sky: S (Gamma: 1.0) | Light: 58.7%**

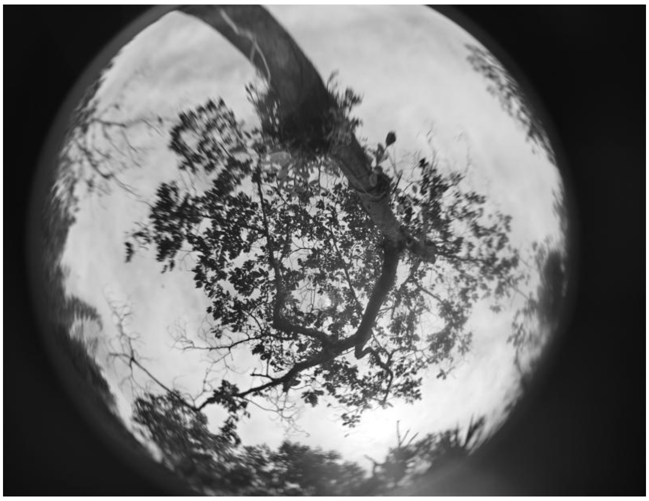
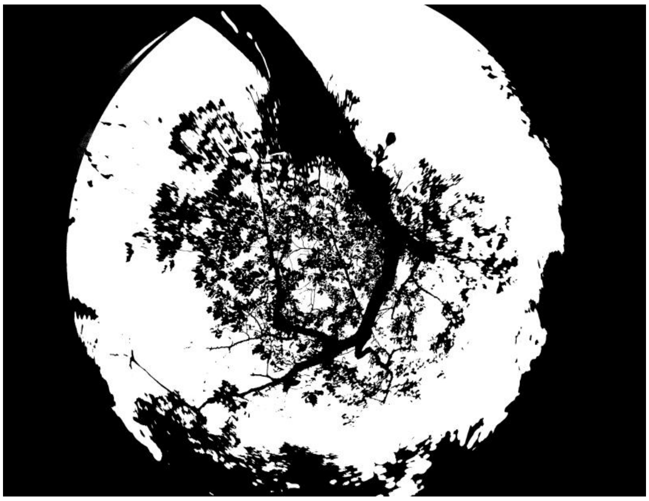

**ID: IMG_20260207_093449.jpg | Sky: S (Gamma: 1.0) | Light: 48.3%**

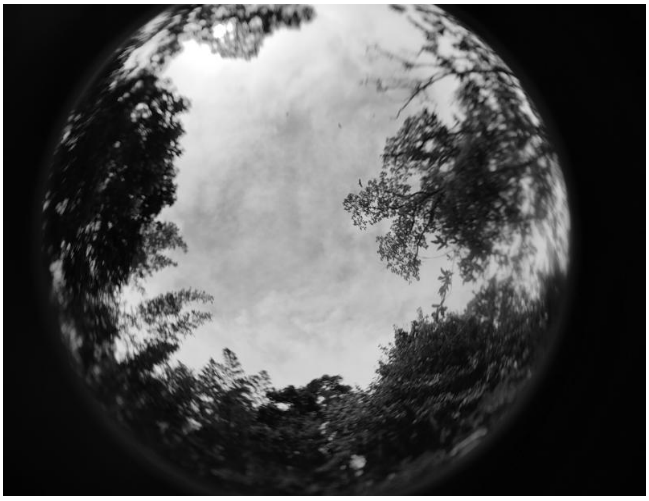
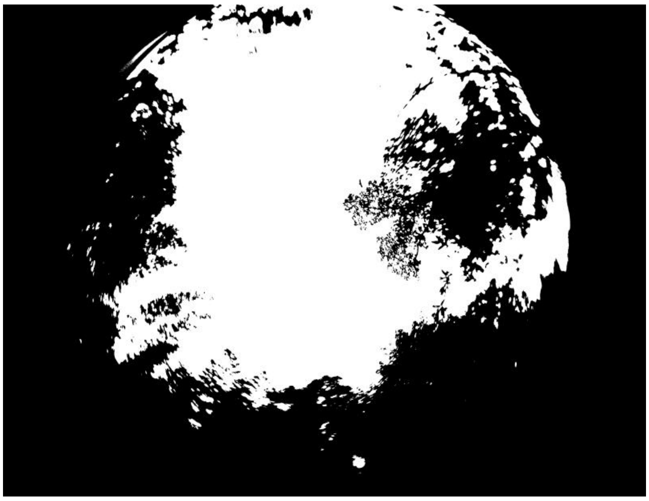

**ID: IMG_20260207_094951.jpg | Sky: S (Gamma: 1.0) | Light: 45.89%**

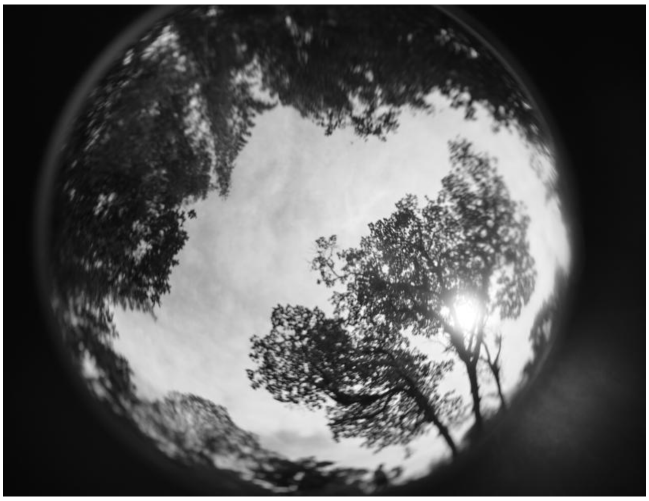
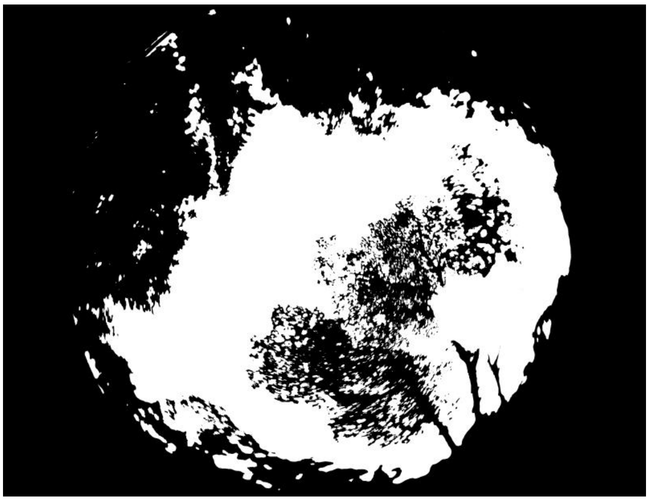

**ID: IMG_20260207_101811.jpg | Sky: P (Gamma: 1.5) | Light: 93.06%**

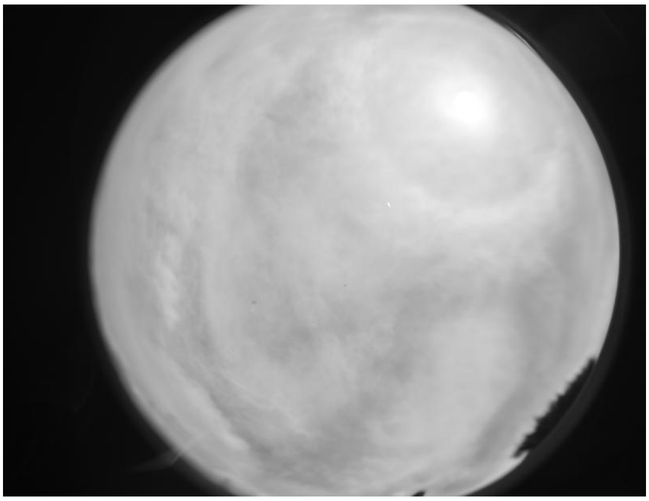

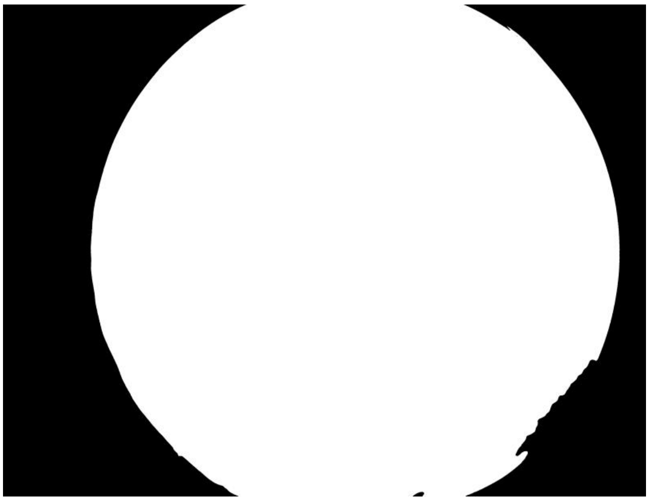

**ID: IMG_20260207_104431.jpg | Sky: P (Gamma: 1.5) | Light: 88.27%**

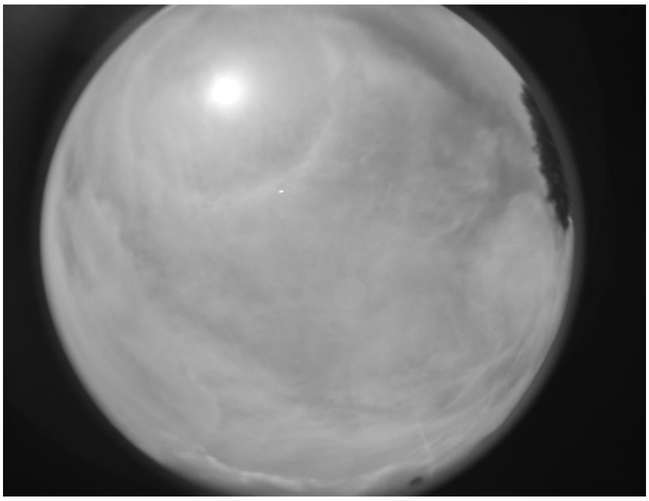

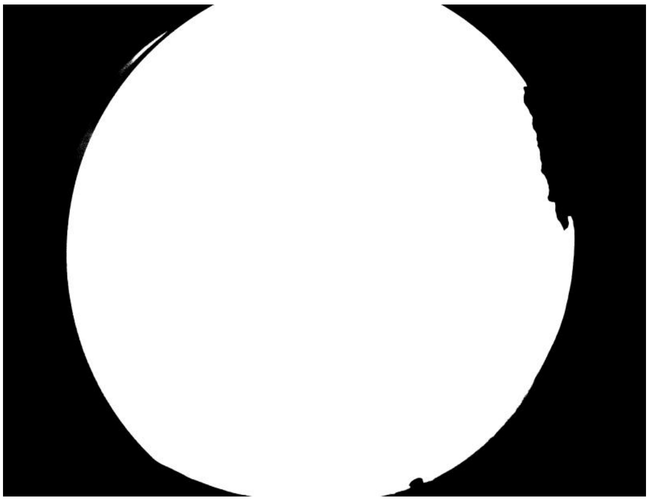

**ID: IMG_20260207_105957.jpg | Sky: P (Gamma: 1.5) | Light: 82.0%**

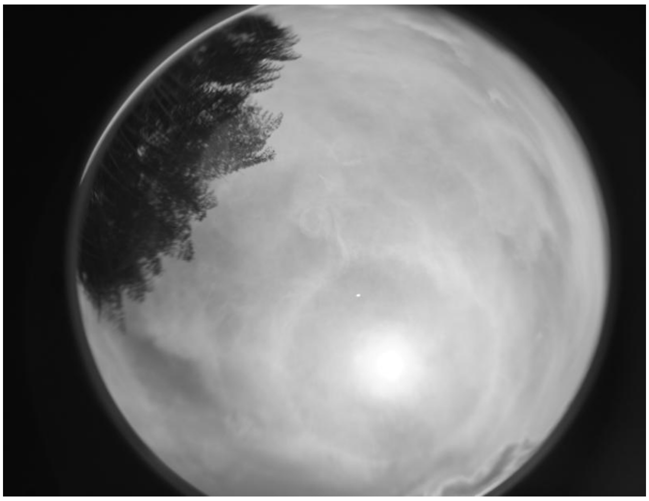

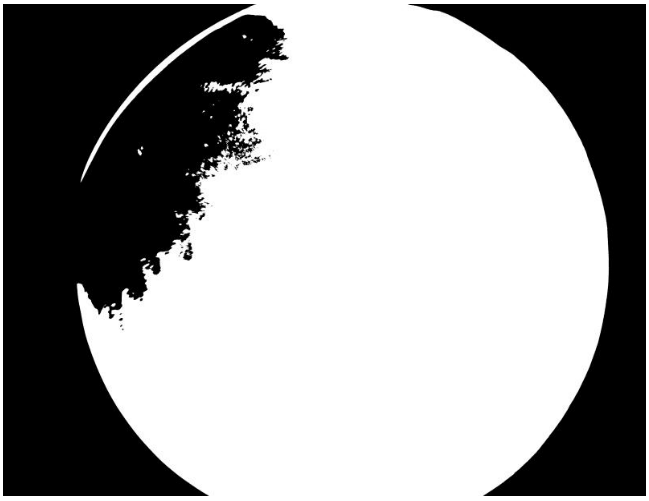

**ID: IMG_20260207_111741.jpg | Sky: P (Gamma: 1.5) | Light: 83.88%**

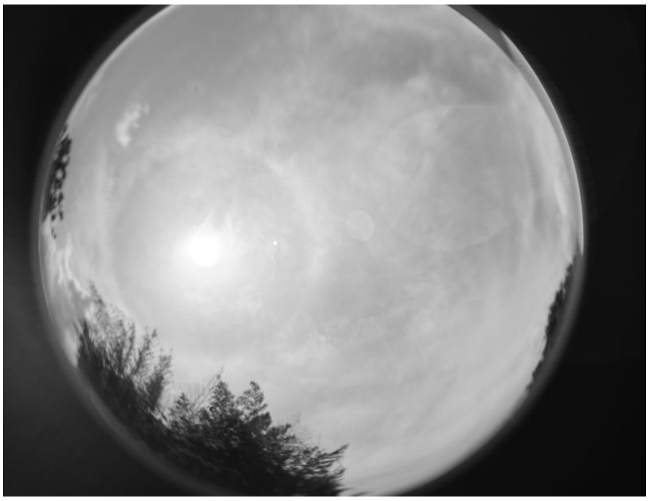
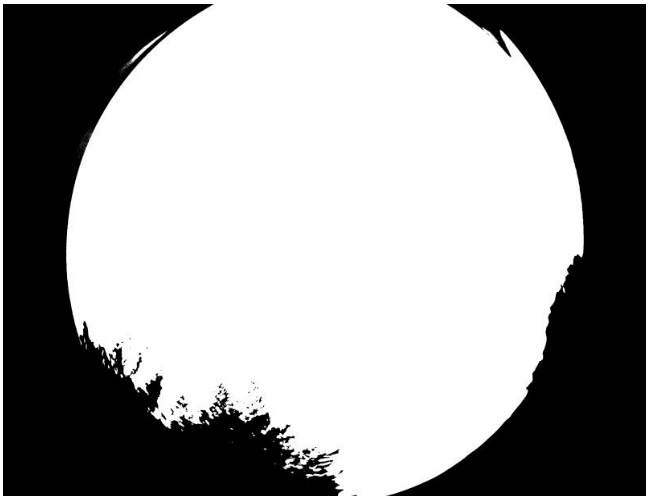

**ID: IMG_20260207_115955.jpg | Sky: P (Gamma: 1.5) | Light: 94.9%**

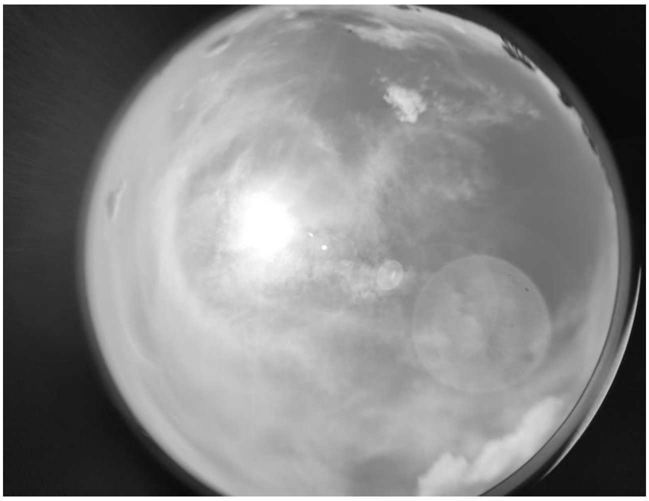
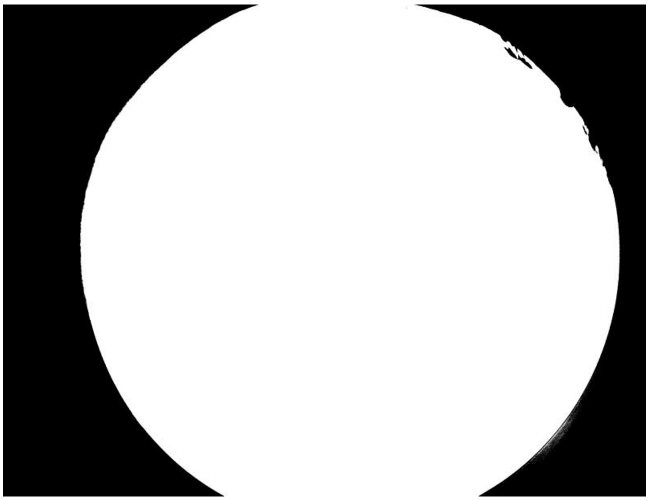

**ID: IMG_20260207_120311.jpg | Sky: P (Gamma: 1.5) | Light: 92.92%**

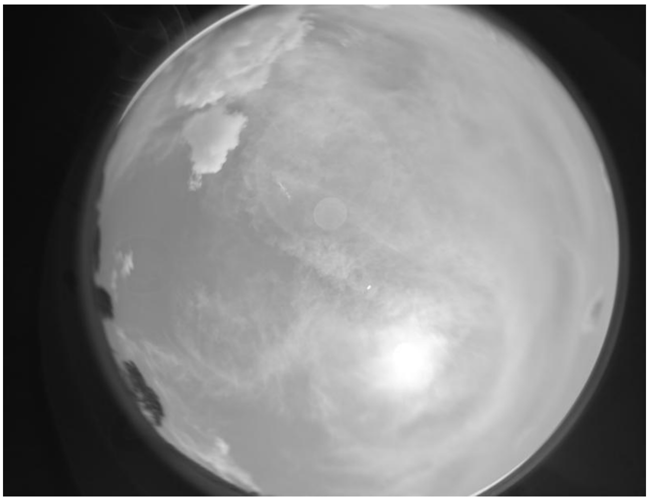
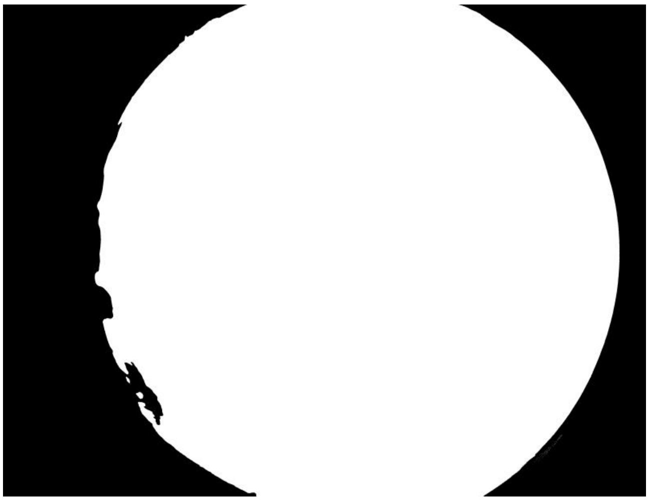

ID: IMG_20260208_092650.jpg | Sky: S (Gamma: 1.0) | Light: 83.3%

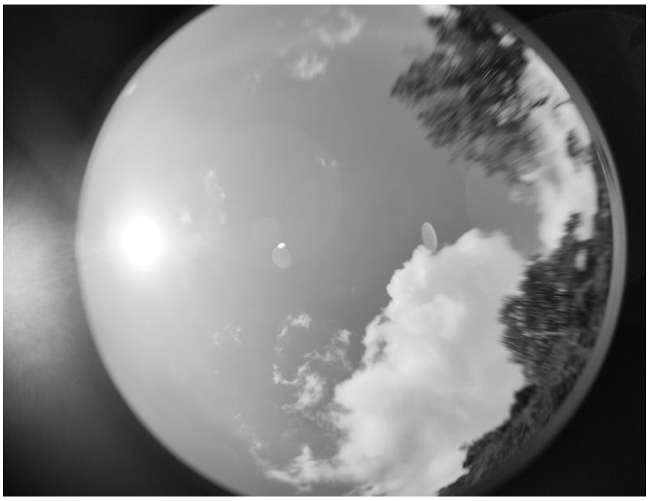
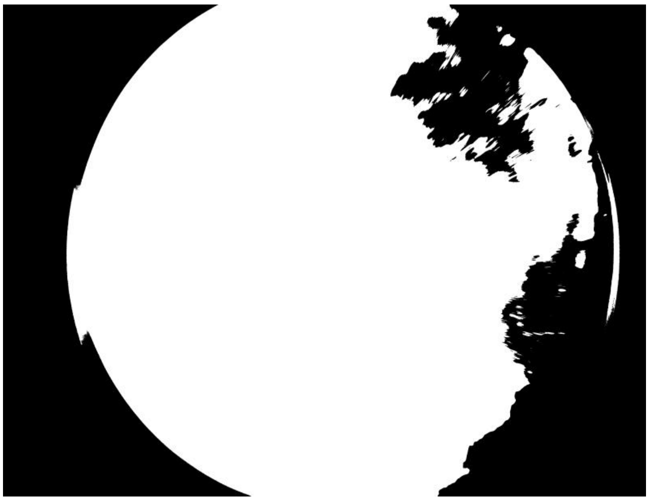

ID: IMG_20260208_094624.jpg | Sky: S (Gamma: 1.0) | Light: 91.65%

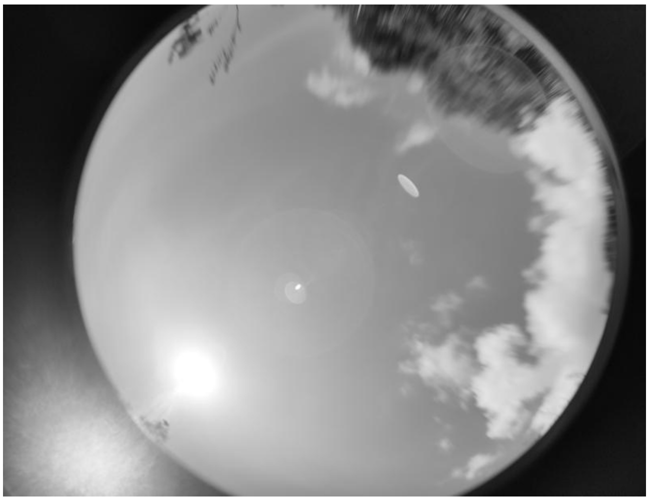
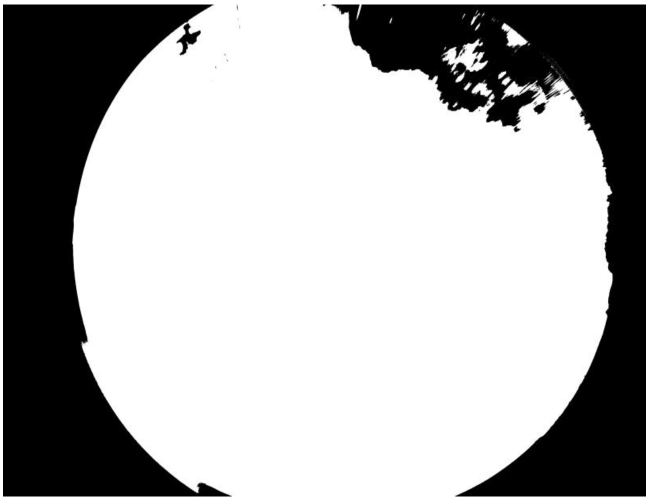

ID: IMG_20260208_100201.jpg | Sky: S (Gamma: 1.0) | Light: 91.46%

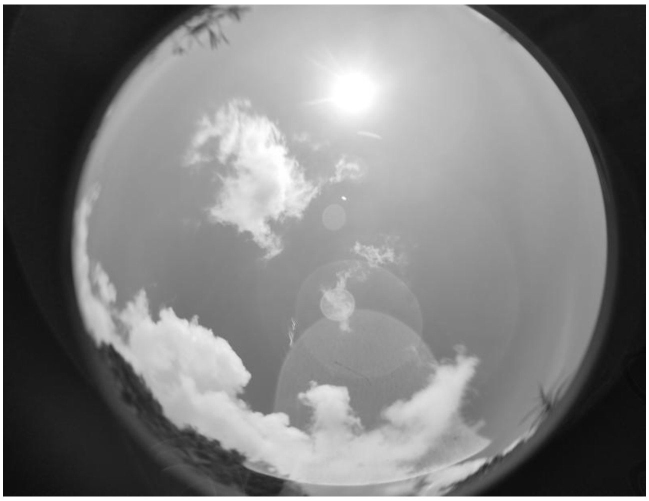
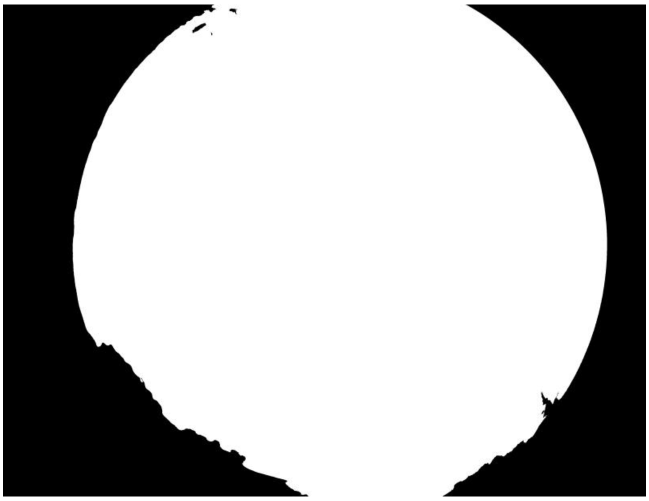

**ID: IMG_20260208_101254.jpg | Sky: S (Gamma: 1.0) | Light: 88.68%**

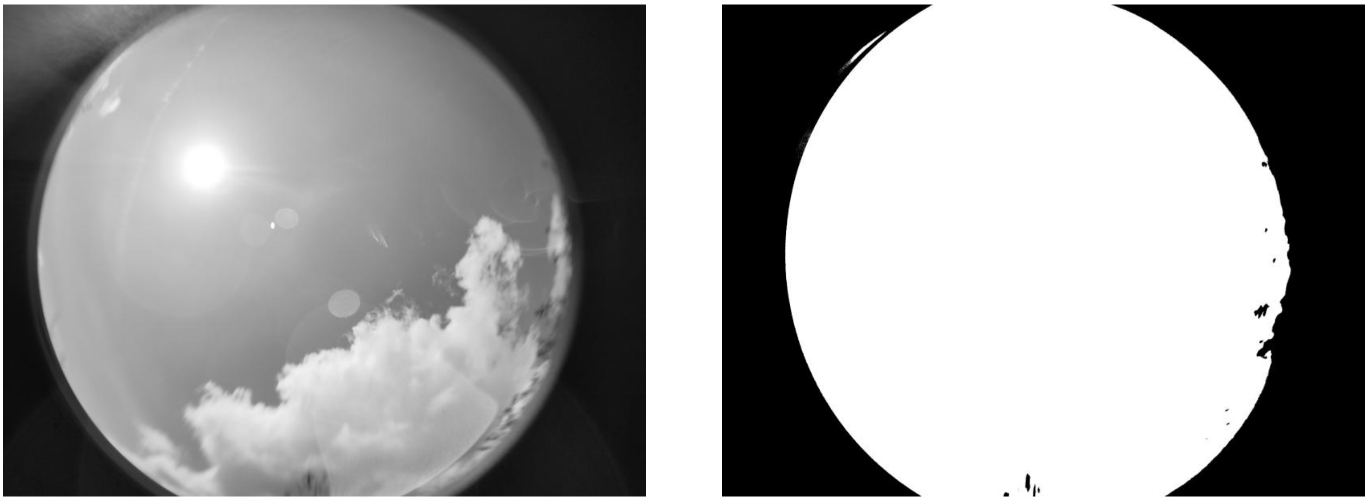

**ID: IMG_20260208_104854.jpg | Sky: S (Gamma: 1.0) | Light: 77.68%**

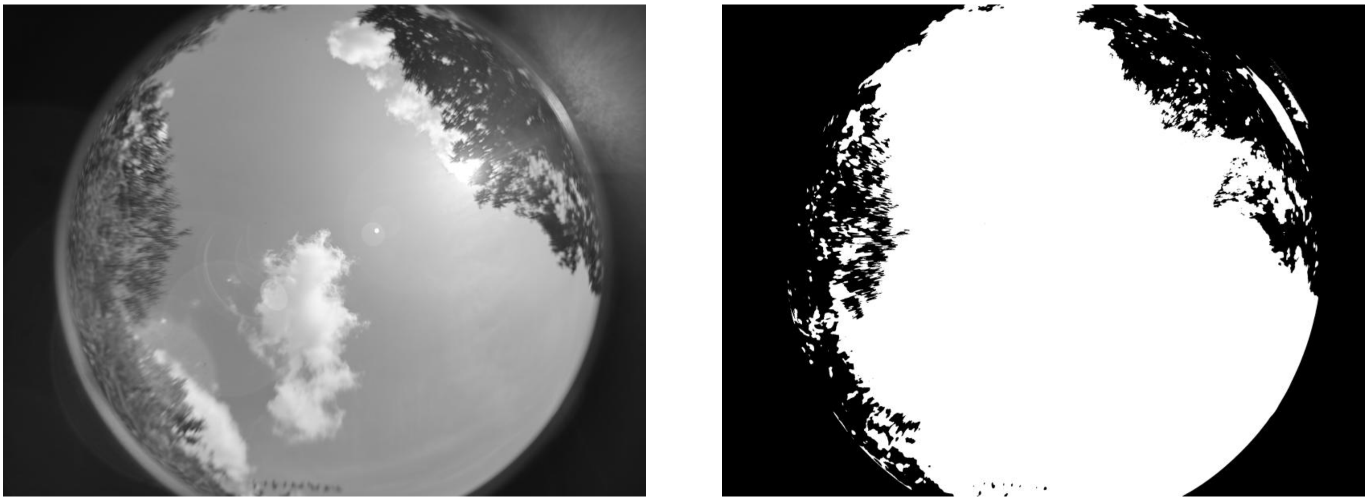

**ID: IMG_20260208_110517.jpg | Sky: S (Gamma: 1.0) | Light: 72.7%**

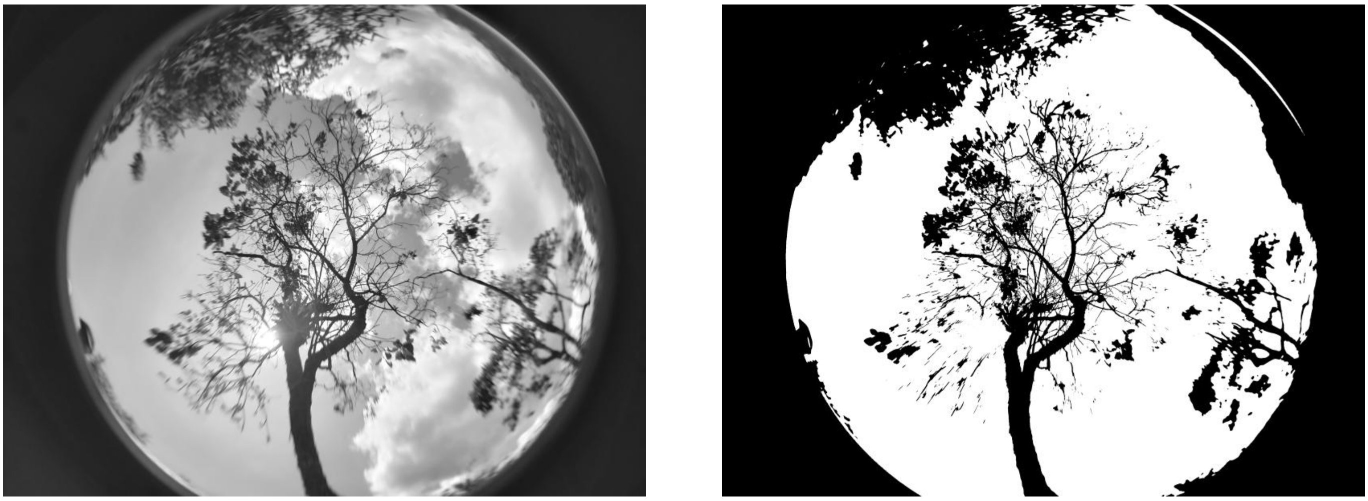

ID: IMG_20260208_112545.jpg | Sky: S (Gamma: 1.0) | Light: 91.36%

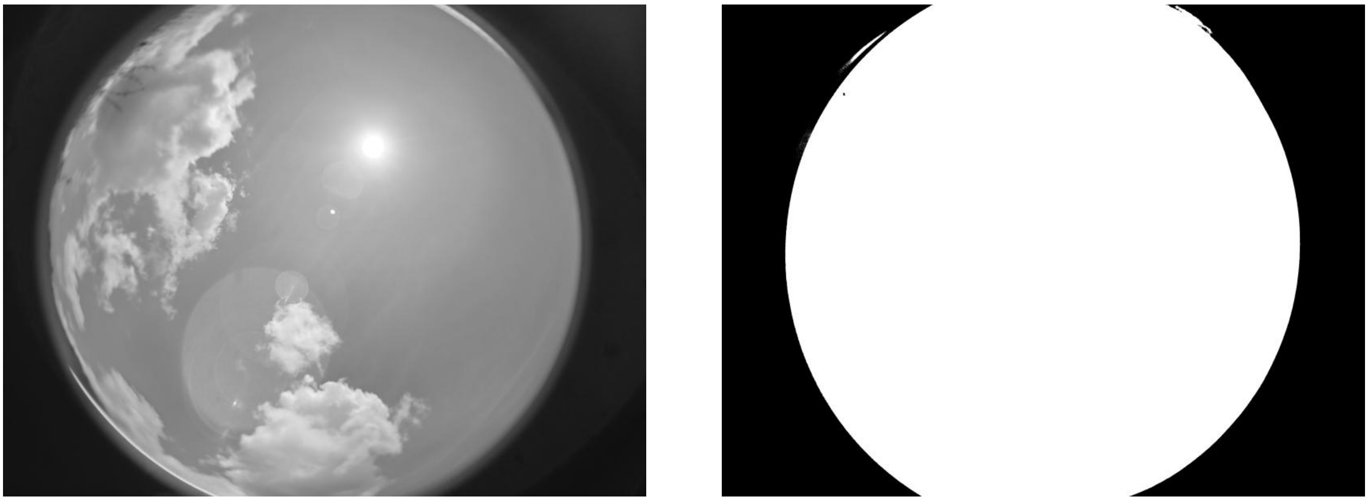

ID: IMG_20260208_115342.jpg | Sky: S (Gamma: 1.0) | Light: 88.15%

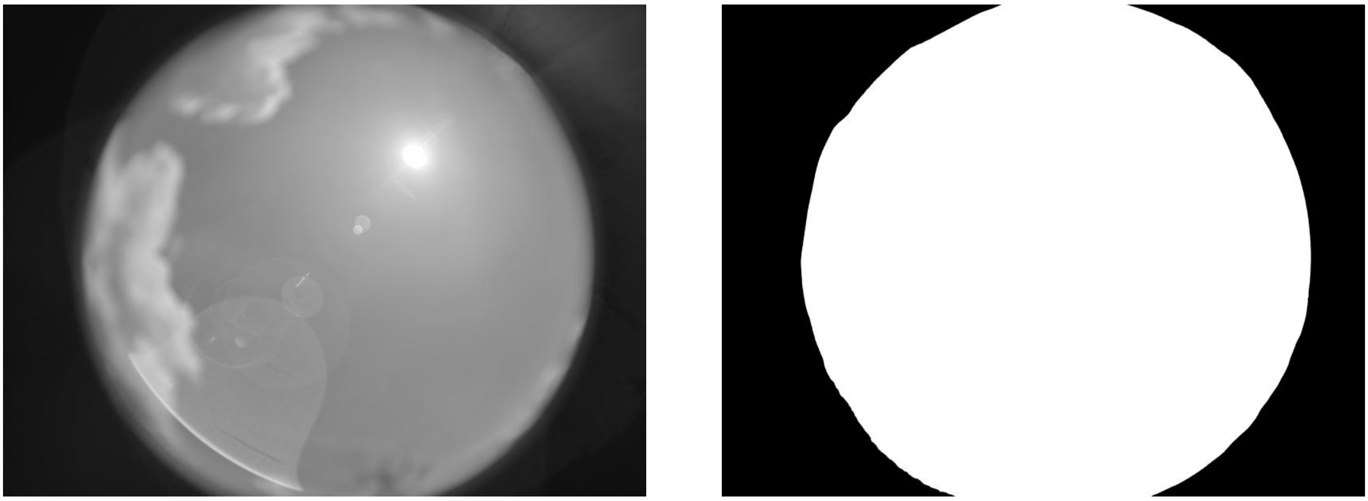

ID: IMG_20260208_120439.jpg | Sky: S (Gamma: 1.0) | Light: 85.82%

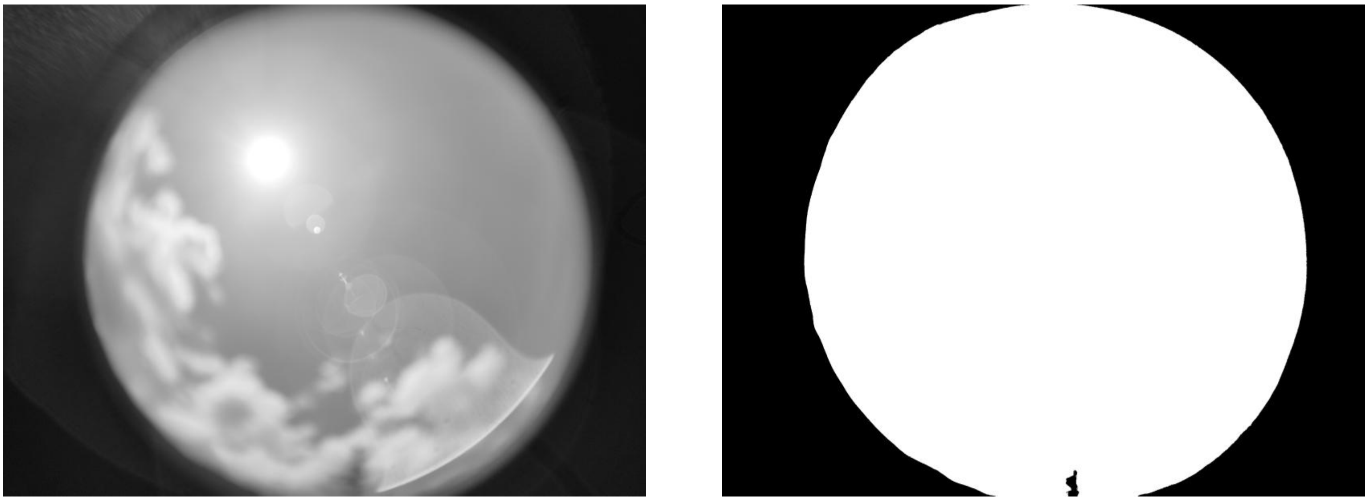

ID: IMG_20260208_121619.jpg | Sky: S (Gamma: 1.0) | Light: 81.1%

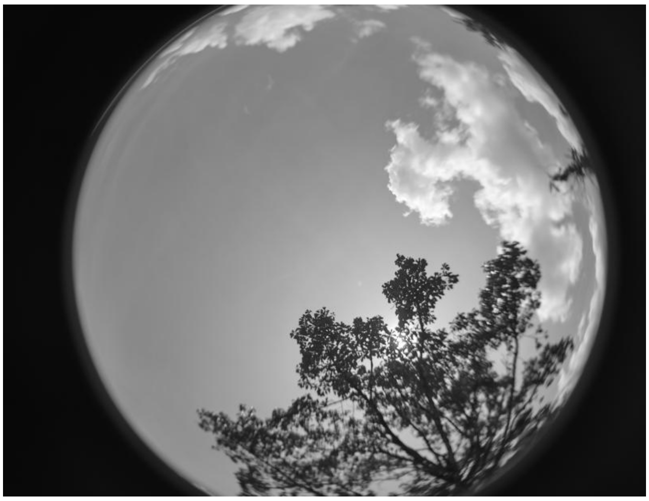

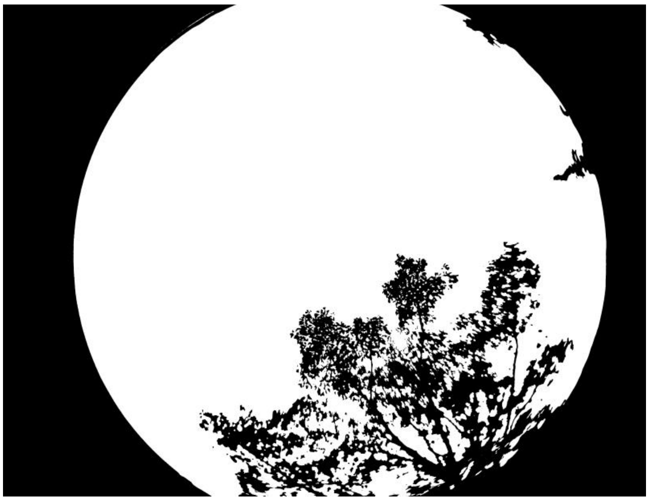

ID: IMG_20260208_124517.jpg | Sky: S (Gamma: 1.0) | Light: 89.54%

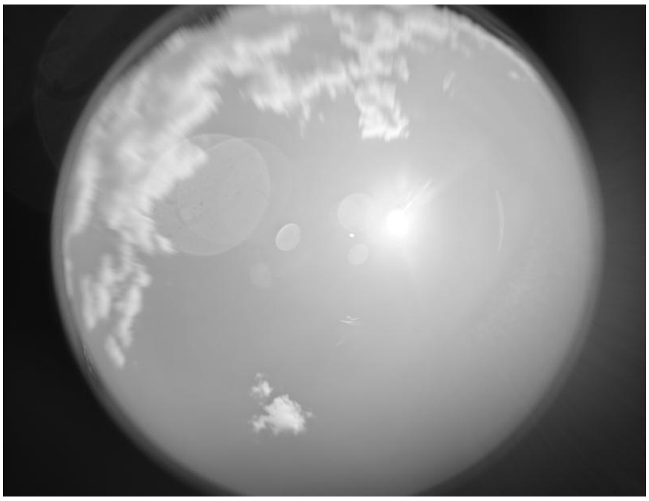

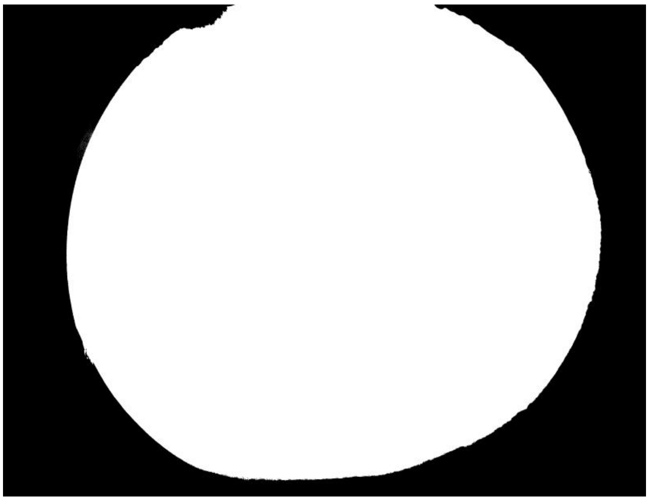

ID: IMG_20260208_125511.jpg | Sky: S (Gamma: 1.0) | Light: 92.65%

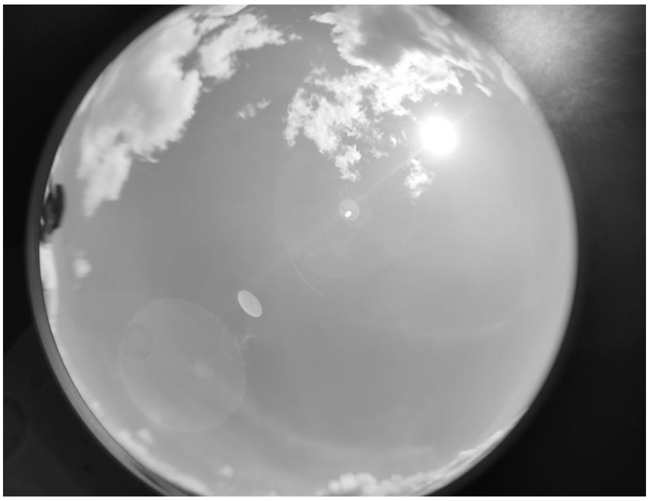

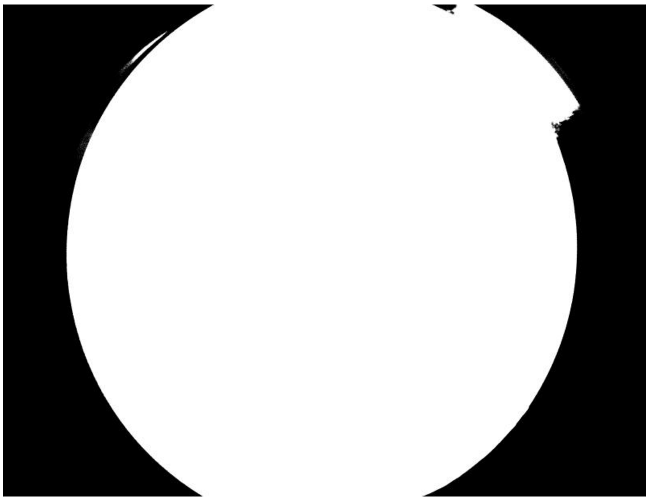

ID: IMG_20260208_130739.jpg | Sky: S (Gamma: 1.0) | Light: 95.32%

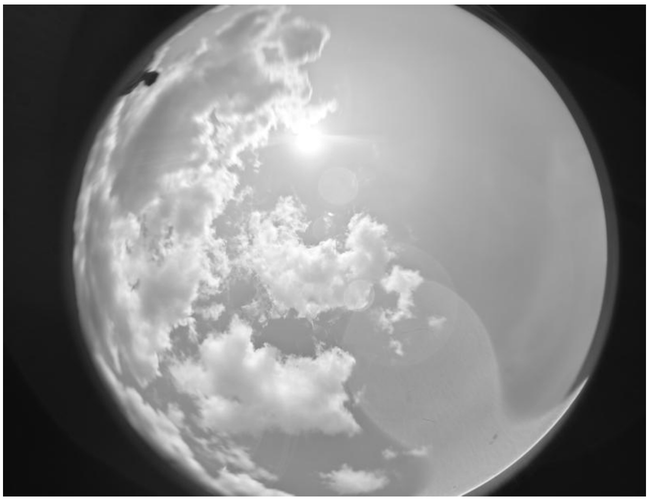

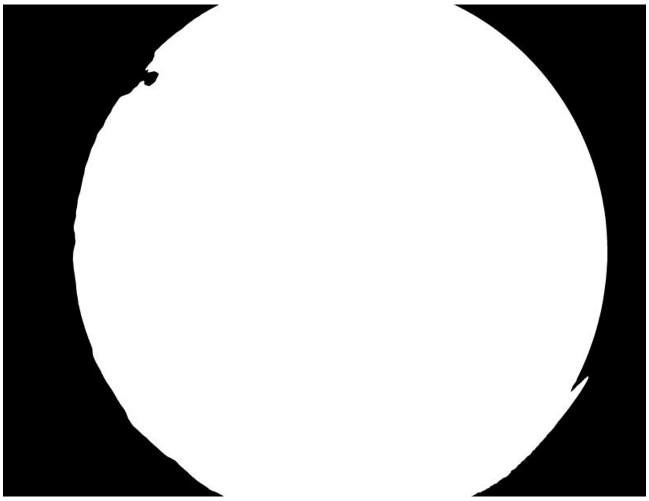

ID: IMG_20260208_131850.jpg | Sky: S (Gamma: 1.0) | Light: 95.04%

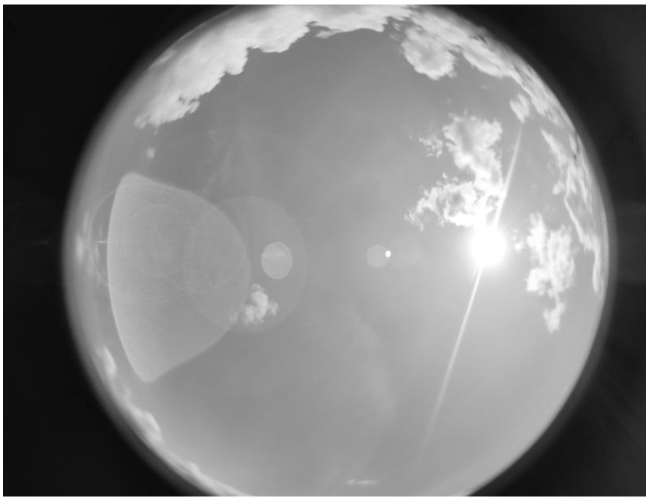

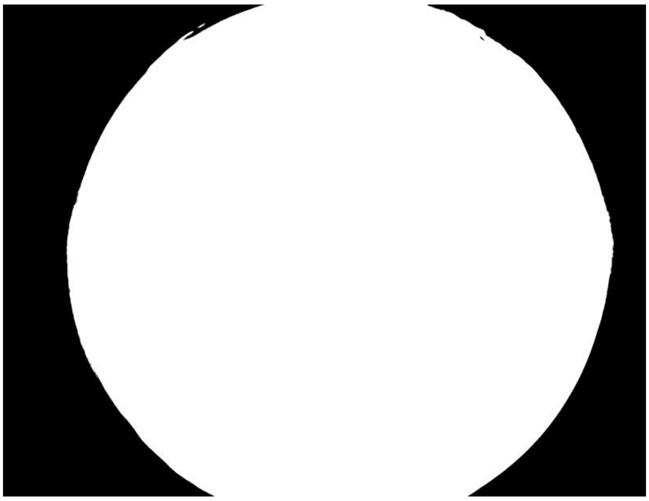

ID: IMG_20260209_112115.jpg | Sky: S (Gamma: 1.0) | Light: 58.15%

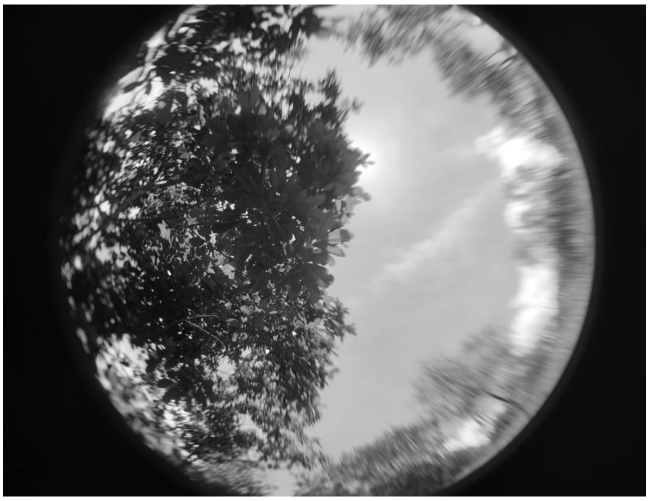

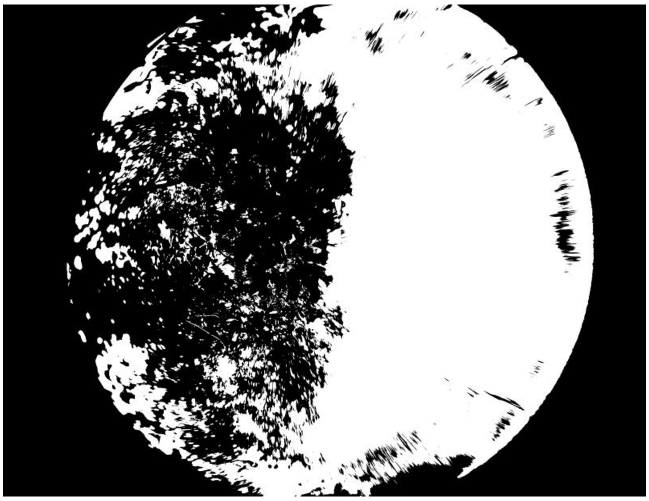

ID: IMG_20260209_114126.jpg | Sky: S (Gamma: 1.0) | Light: 56.63%

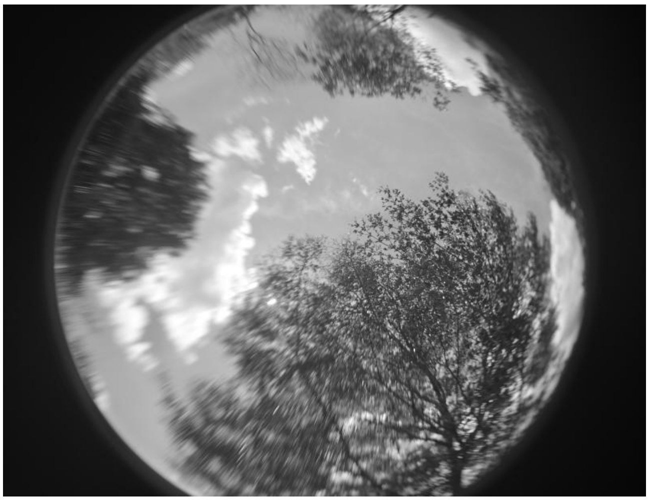

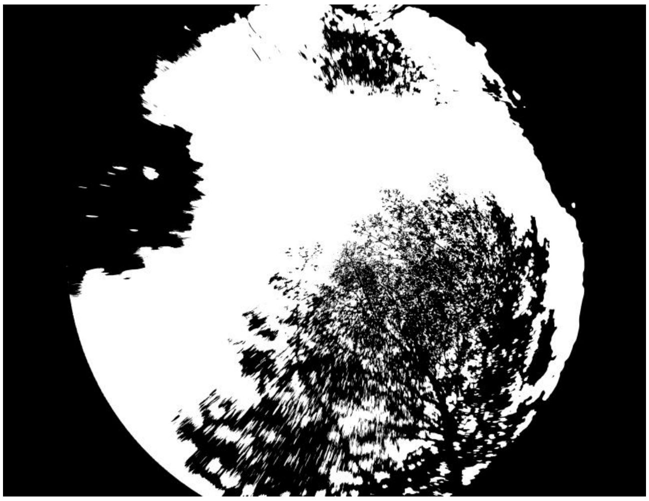

ID: IMG_20260209_115649.jpg | Sky: S (Gamma: 1.0) | Light: 47.76%

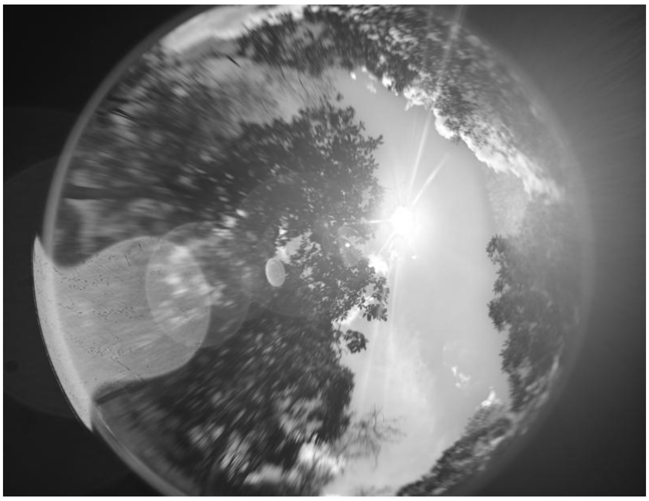

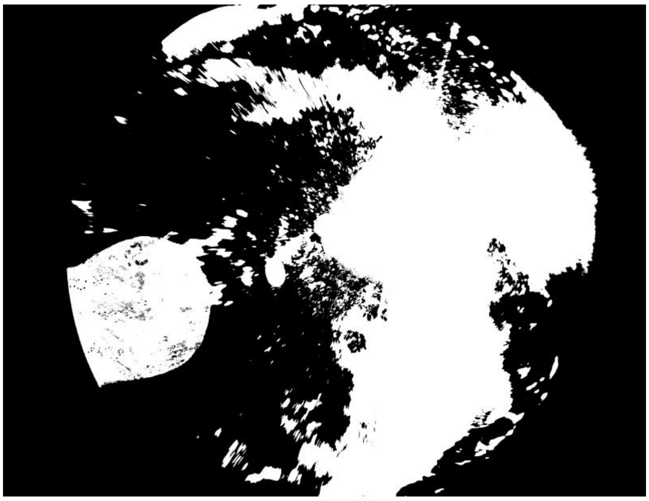

ID: IMG_20260209_121905.jpg | Sky: S (Gamma: 1.0) | Light: 72.49%

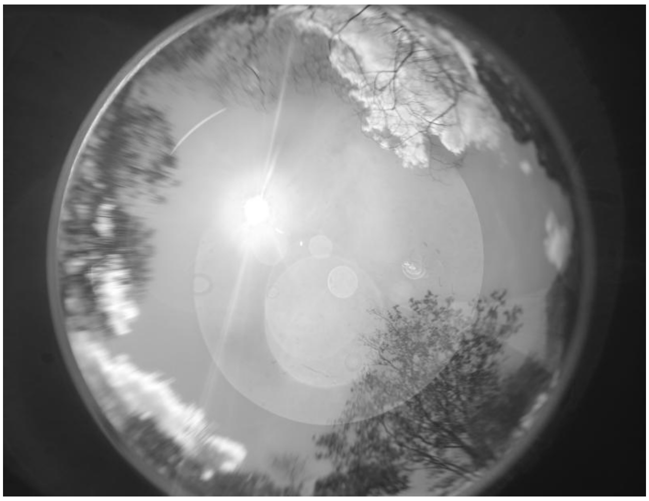

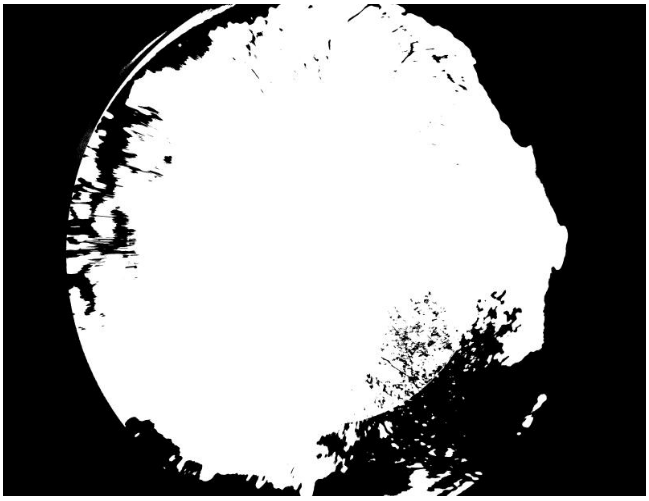

ID: IMG_20260209_123556.jpg | Sky: S (Gamma: 1.0) | Light: 77.27%

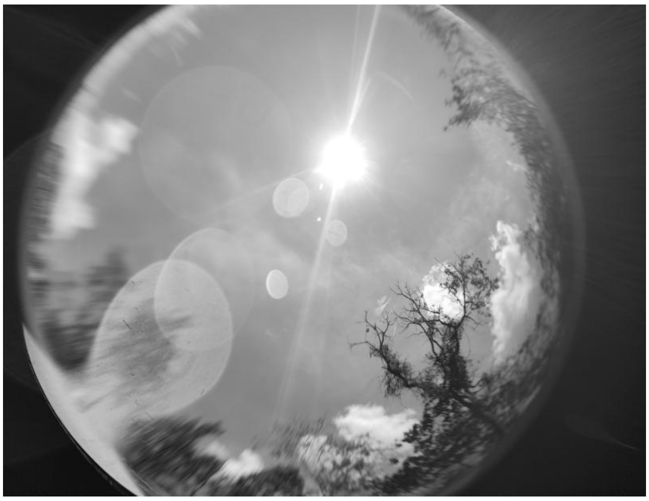

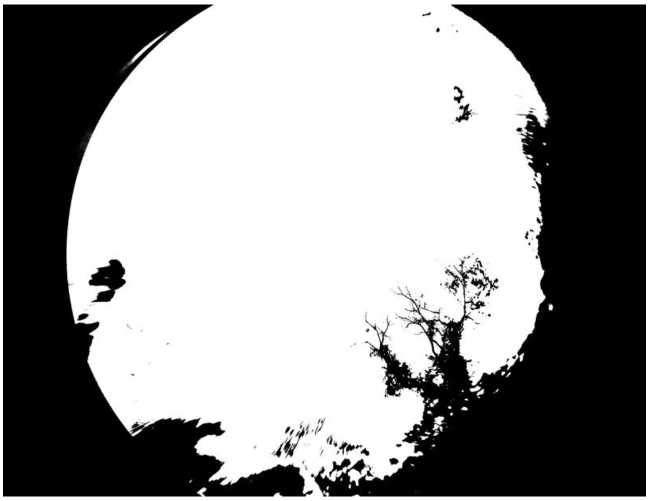

ID: IMG_20260211_104406.jpg | Sky: S (Gamma: 1.0) | Light: 33.24%

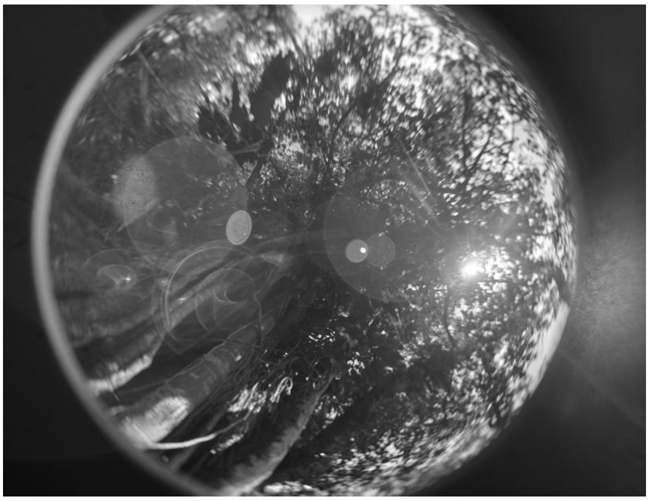

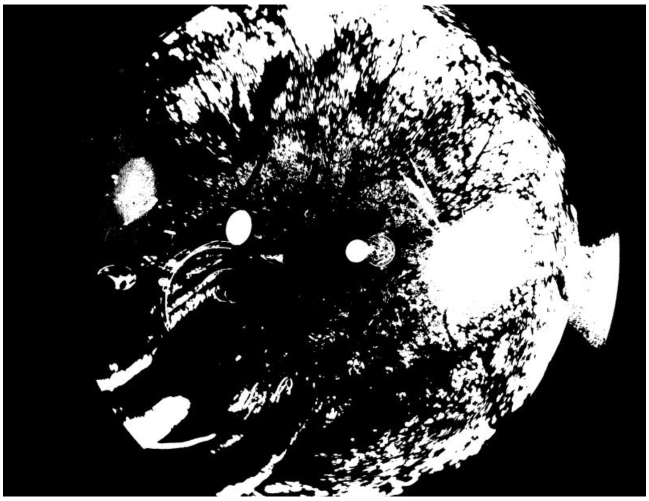

ID: IMG_20260211_110837.jpg | Sky: S (Gamma: 1.0) | Light: 57.65%

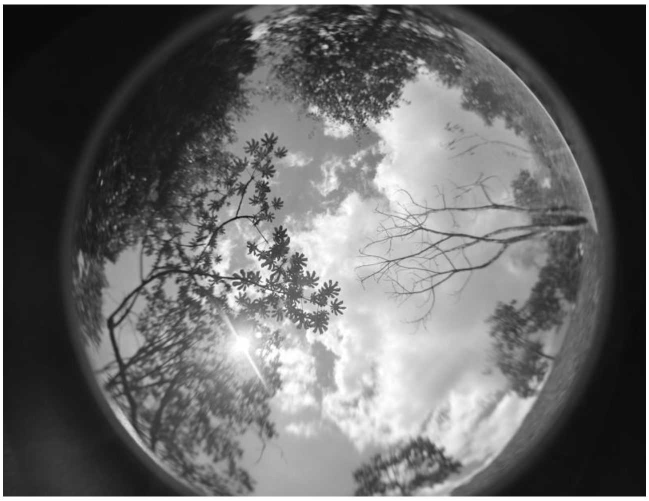

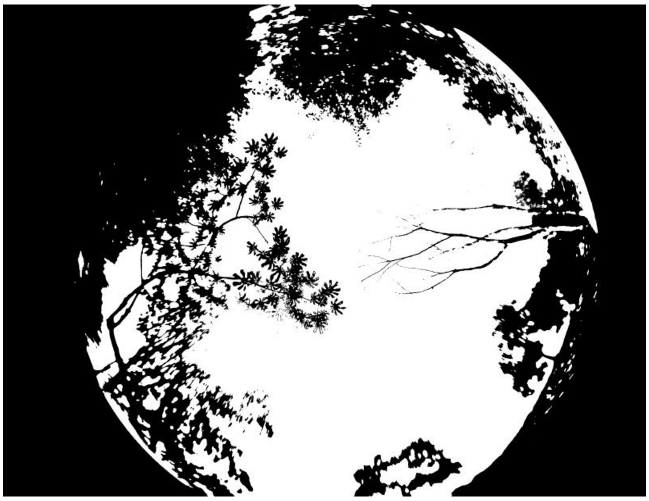

**ID: IMG_20260211_112655.jpg | Sky: S (Gamma: 1.0) | Light: 56.07%**

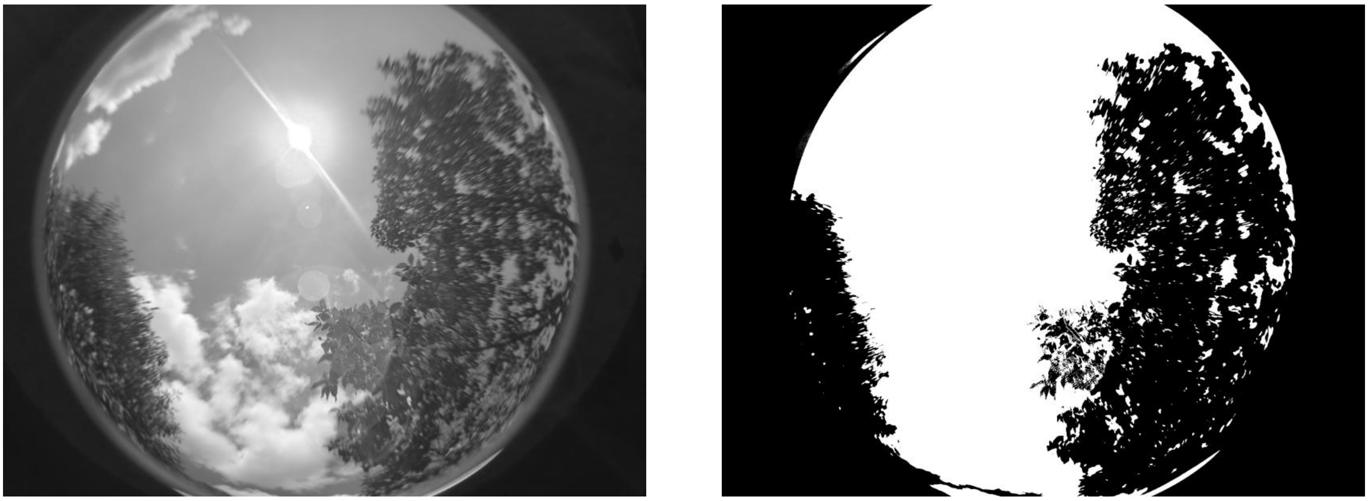

**ID: IMG_20260211_114340.jpg | Sky: S (Gamma: 1.0) | Light: 77.9%**

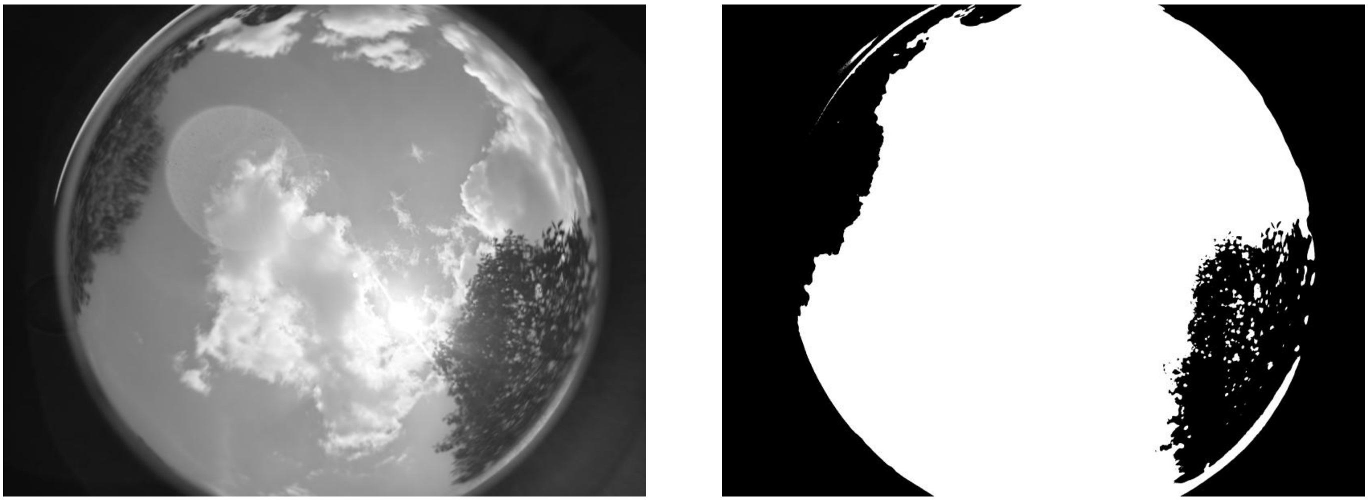

**ID: IMG_20260211_121047.jpg | Sky: S (Gamma: 1.0) | Light: 24.24%**

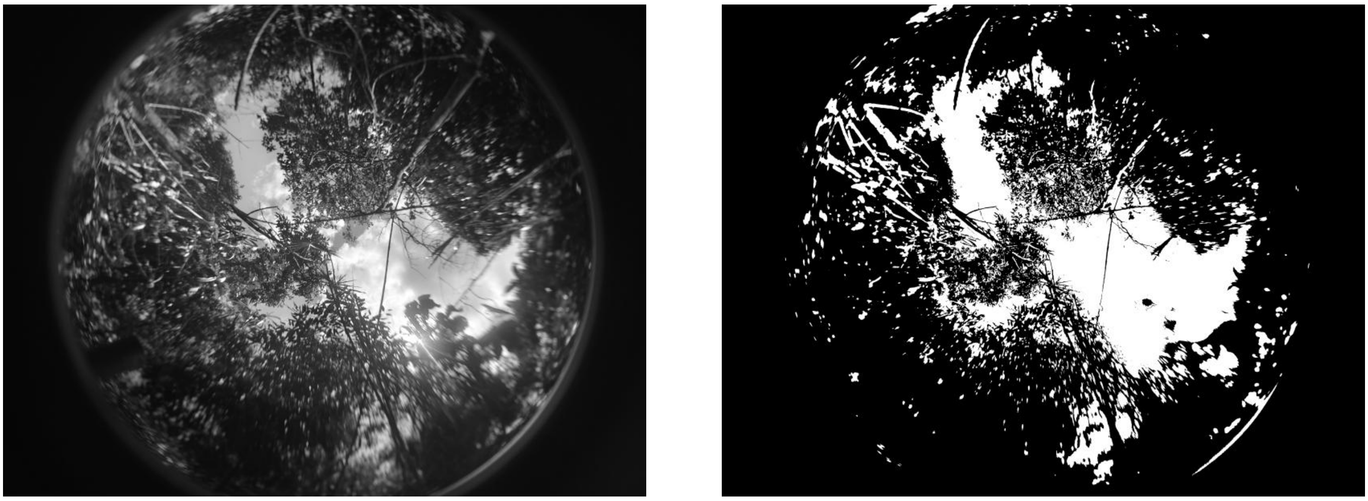

ID: IMG_20260211_125503.jpg | Sky: S (Gamma: 1.0) | Light: 55.66%

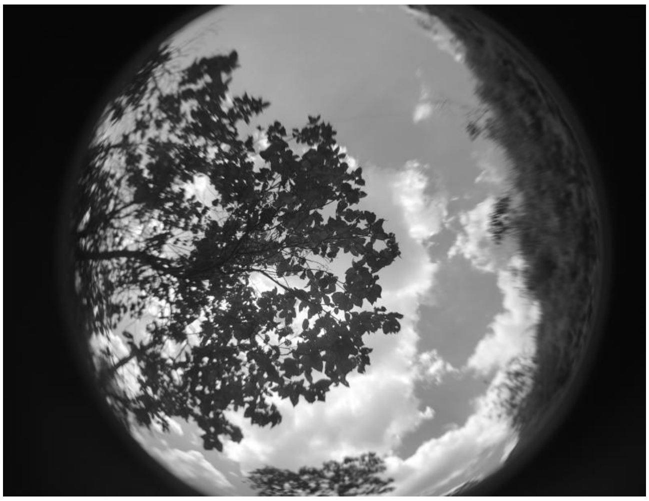

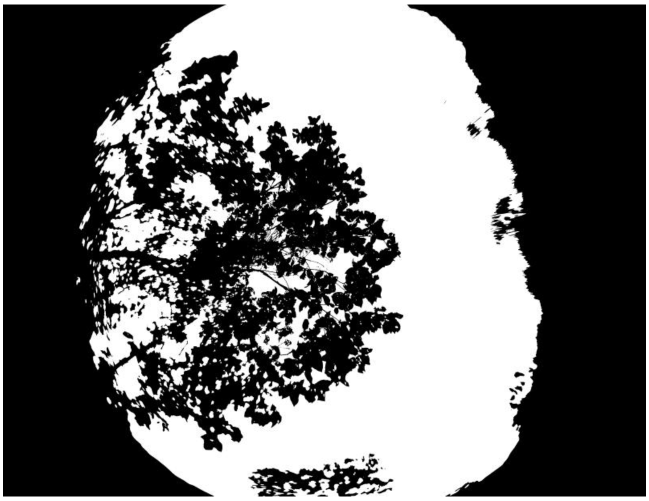

ID: IMG_20260212_091742.jpg | Sky: S (Gamma: 1.0) | Light: 89.24%

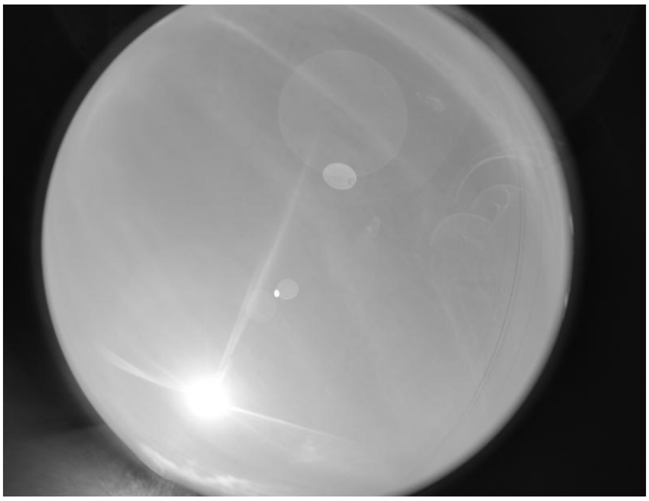

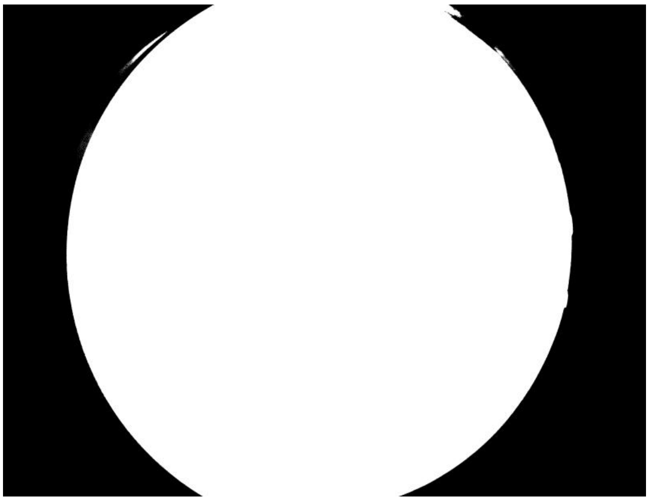

ID: IMG_20260212_092837.jpg | Sky: S (Gamma: 1.0) | Light: 95.01%

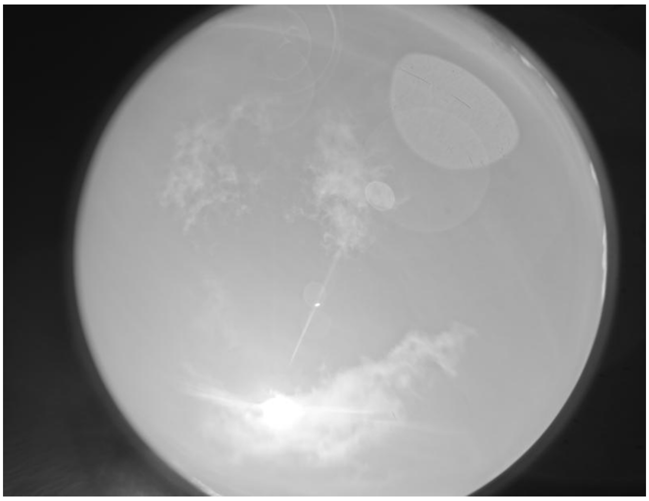

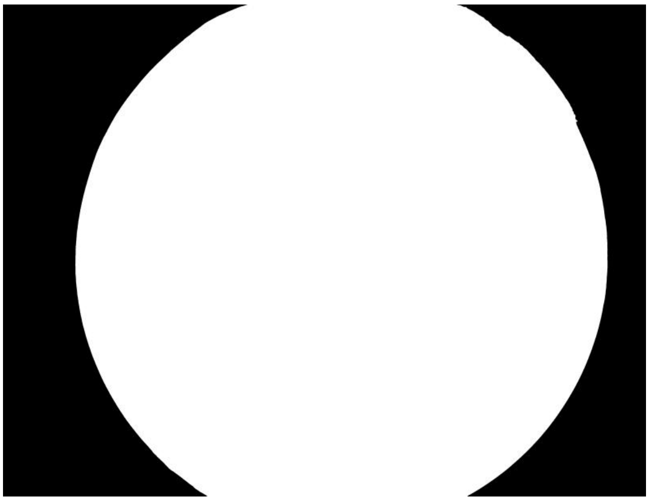

**ID: IMG_20260212_094538.jpg | Sky: S (Gamma: 1.0) | Light: 90.35%**

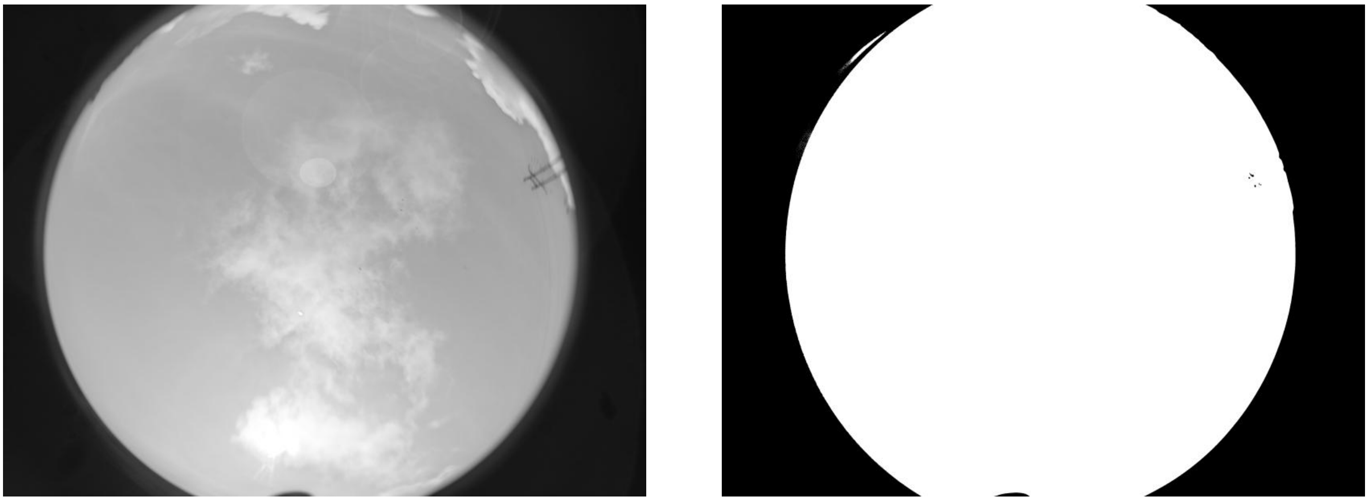

**ID: IMG_20260212_095741.jpg | Sky: S (Gamma: 1.0) | Light: 87.05%**

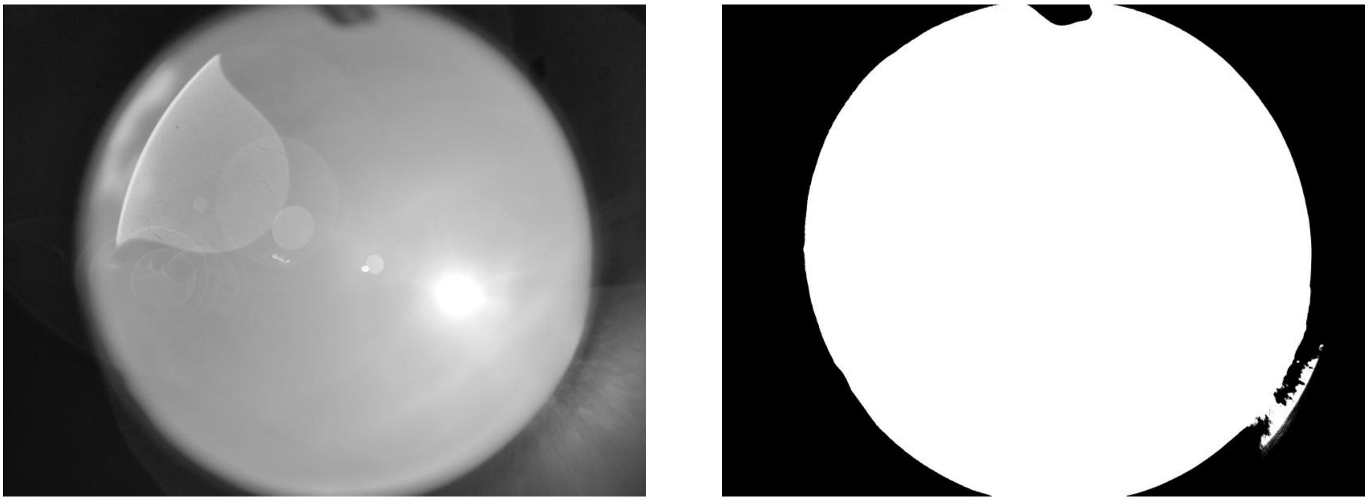

**ID: IMG_20260212_101029.jpg | Sky: S (Gamma: 1.0) | Light: 87.45%**

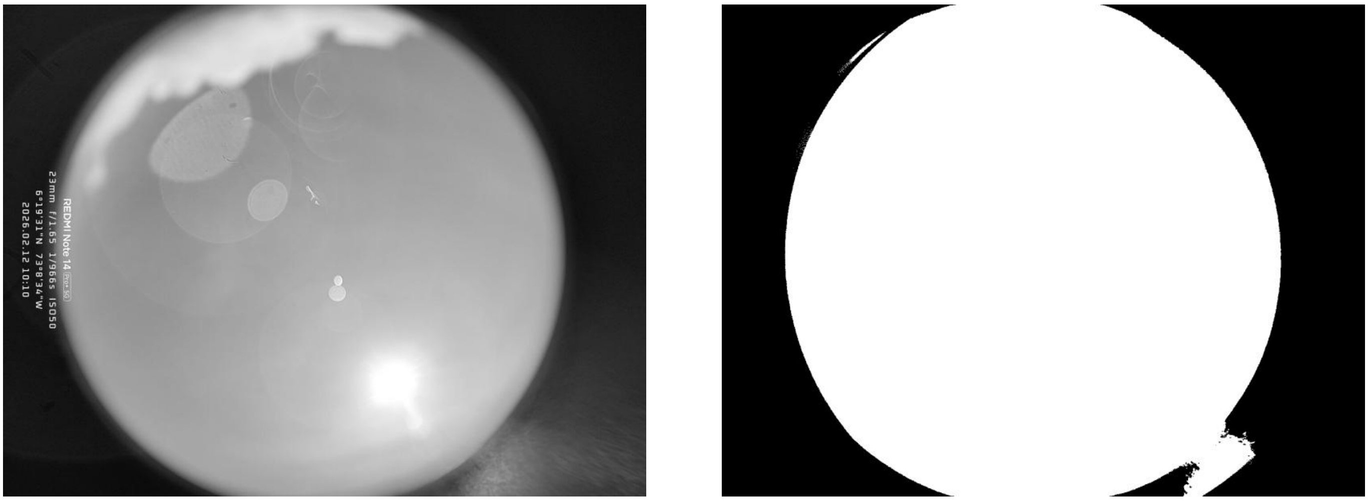

**ID: IMG_20260212_102104.jpg | Sky: S (Gamma: 1.0) | Light: 90.85%**

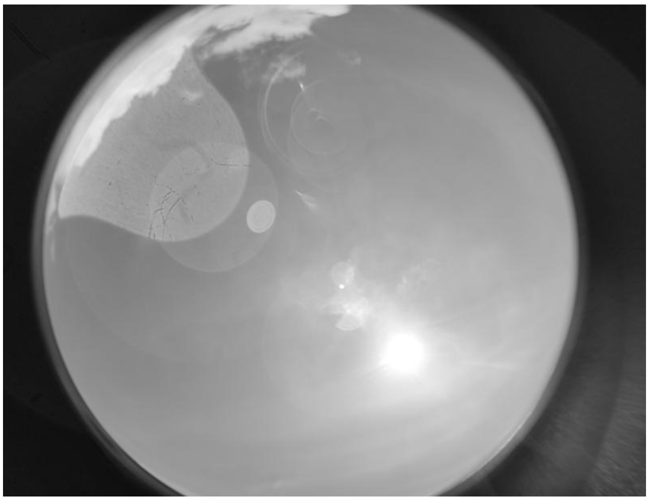

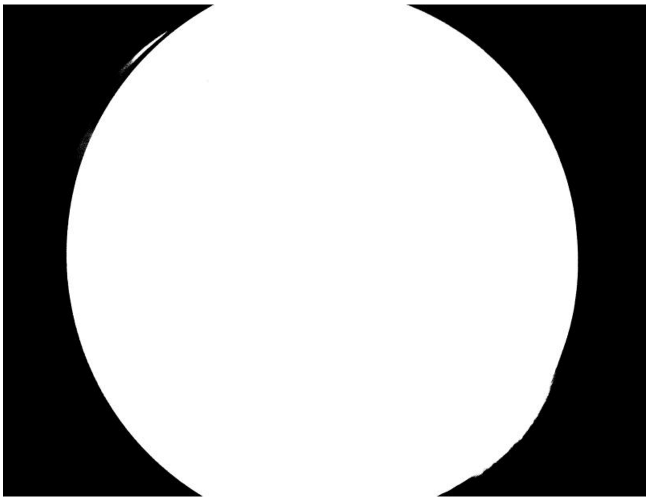

**ID: IMG_20260212_103523.jpg | Sky: S (Gamma: 1.0) | Light: 86.99%**

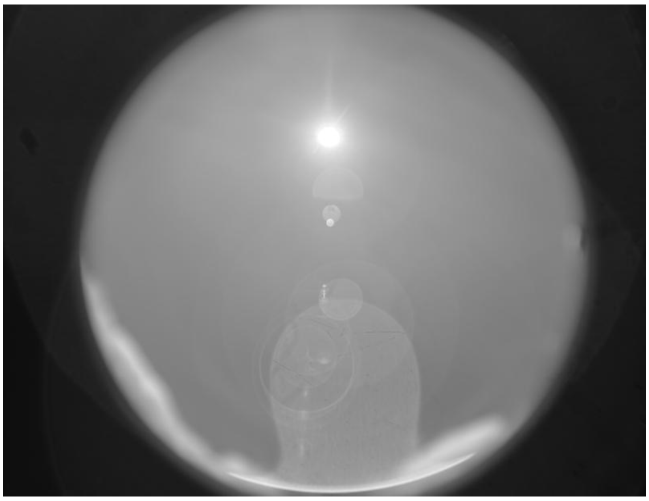

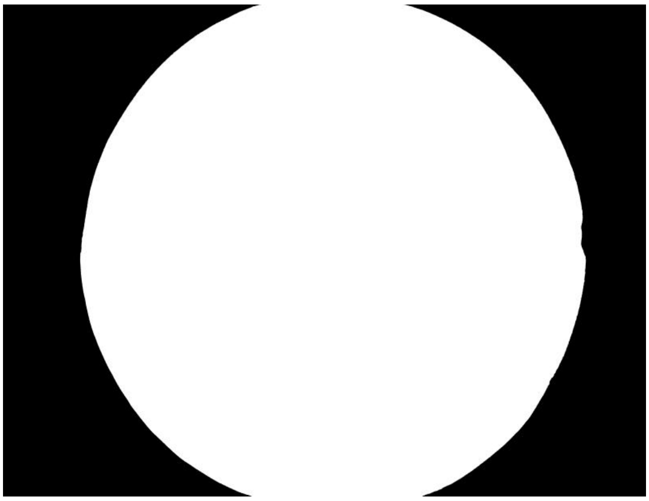

**ID: IMG_20260212_104655.jpg | Sky: S (Gamma: 1.0) | Light: 95.44%**

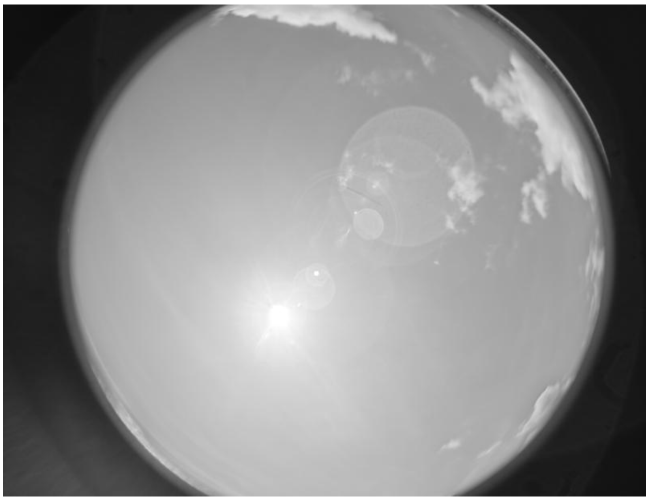

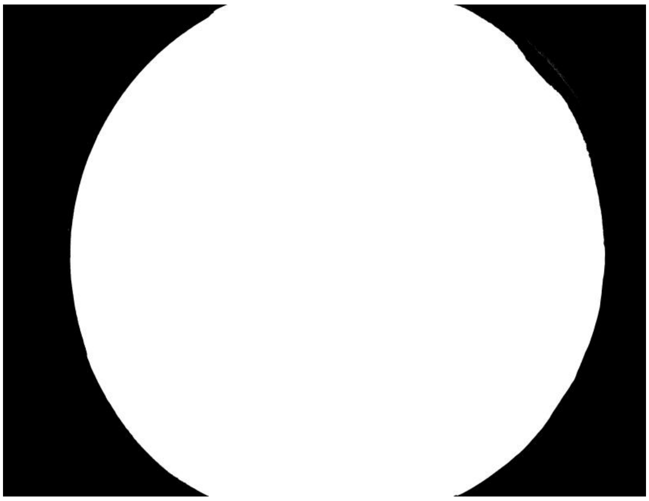

**ID: IMG_20260212_110206.jpg | Sky: S (Gamma: 1.0) | Light: 89.34%**

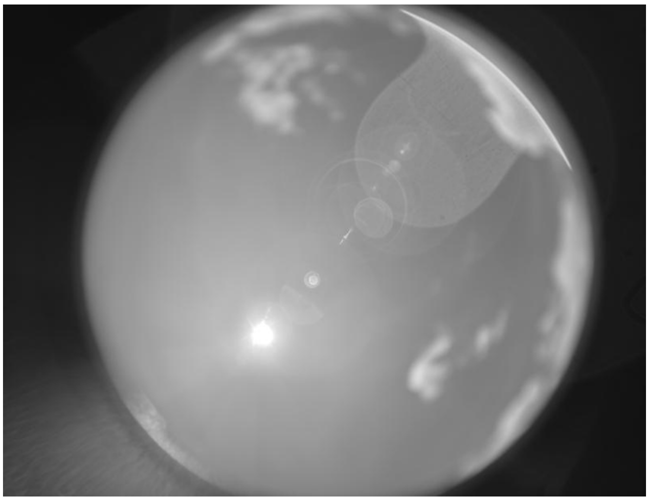

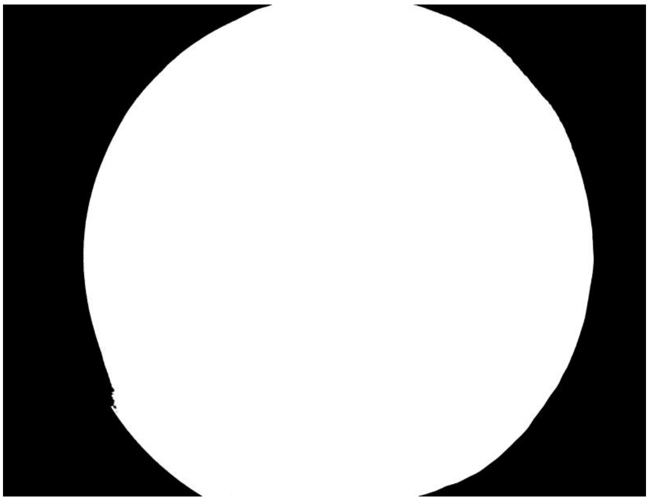

**ID: IMG_20260212_112159.jpg | Sky: S (Gamma: 1.0) | Light: 93.76%**

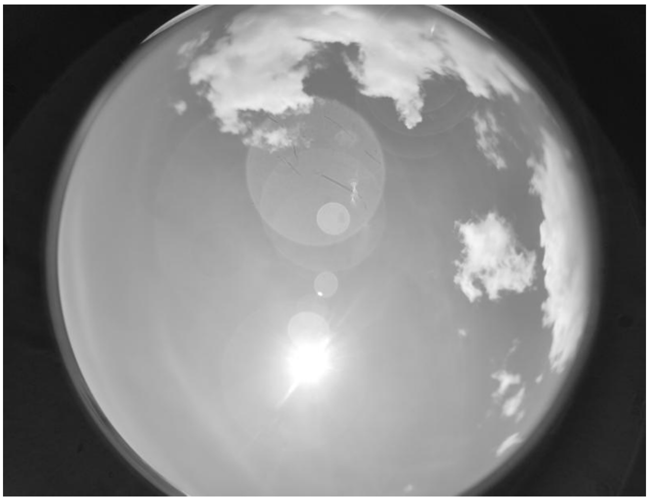

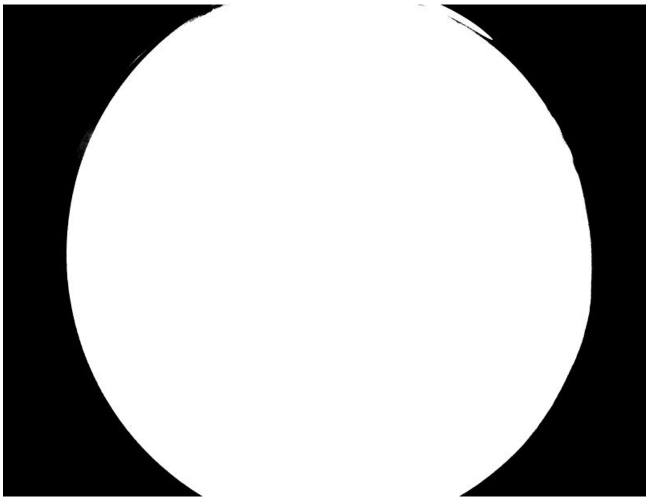

**ID: IMG_20260212_113543.jpg | Sky: S (Gamma: 1.0) | Light: 91.92%**

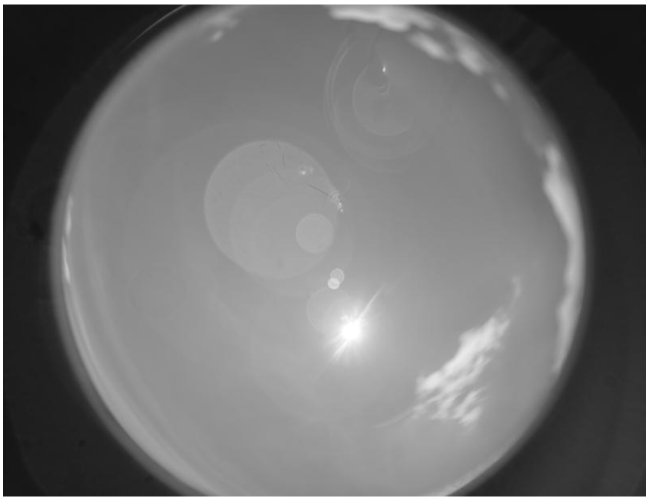

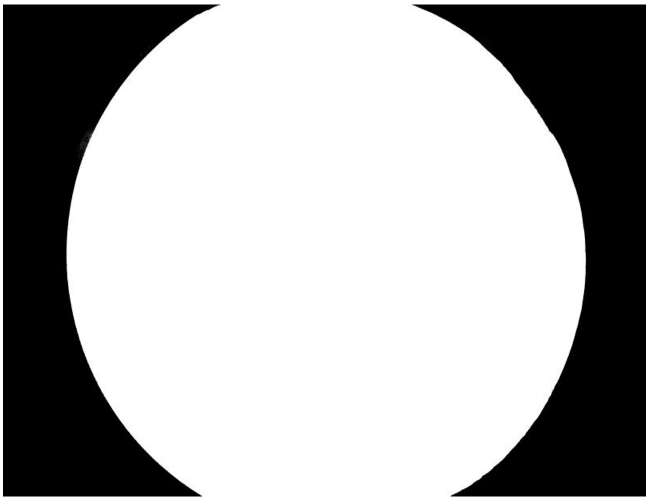

**ID: IMG_20260212_114901.jpg | Sky: S (Gamma: 1.0) | Light: 93.06%**

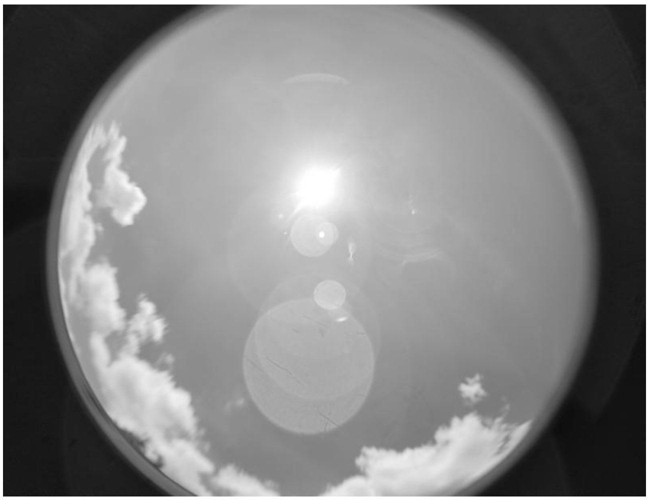

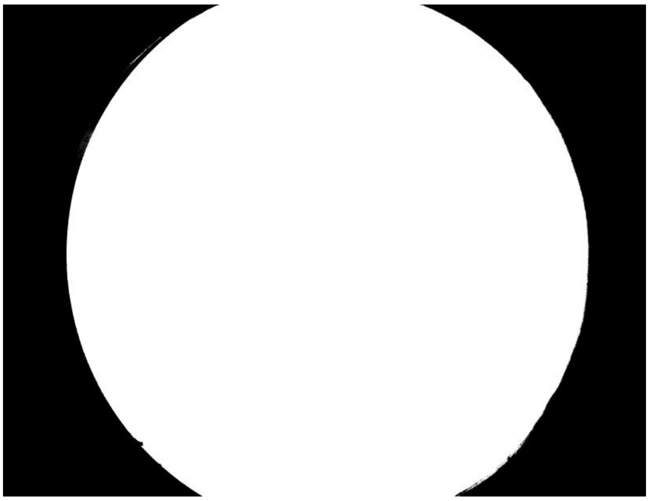

**ID: IMG_20260212_115823.jpg | Sky: S (Gamma: 1.0) | Light: 91.8%**

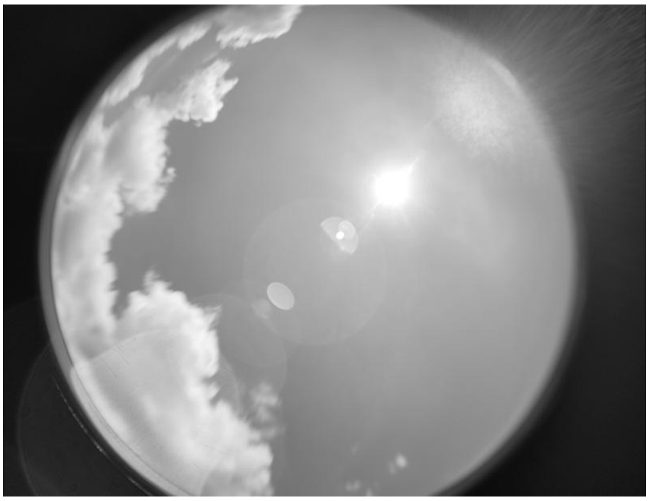

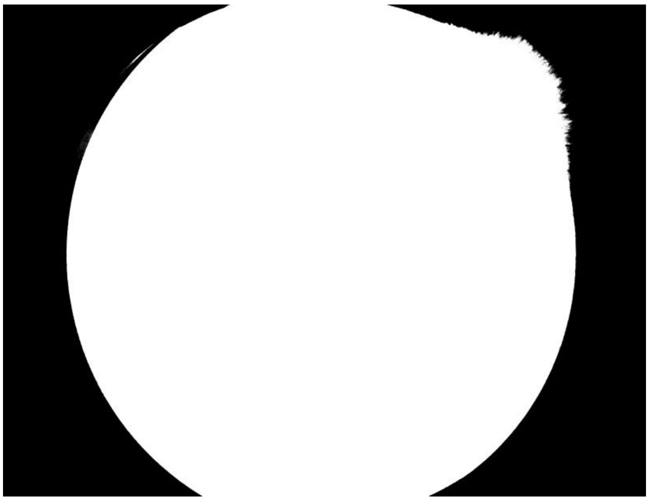

**ID: IMG_20260213_151135.jpg | Sky: P (Gamma: 1.5) | Light: 50.98%**

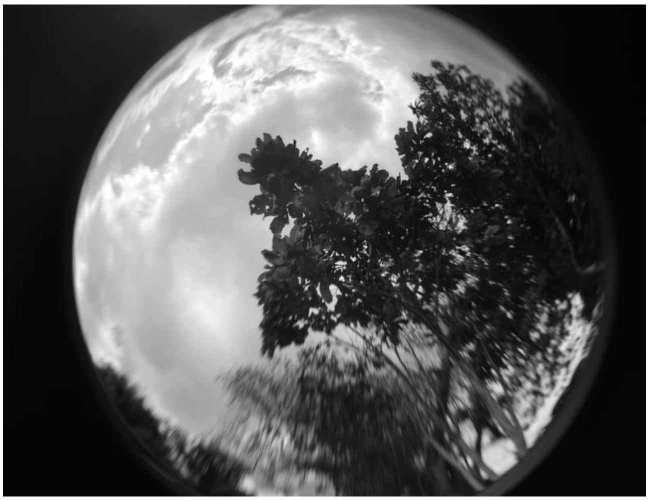

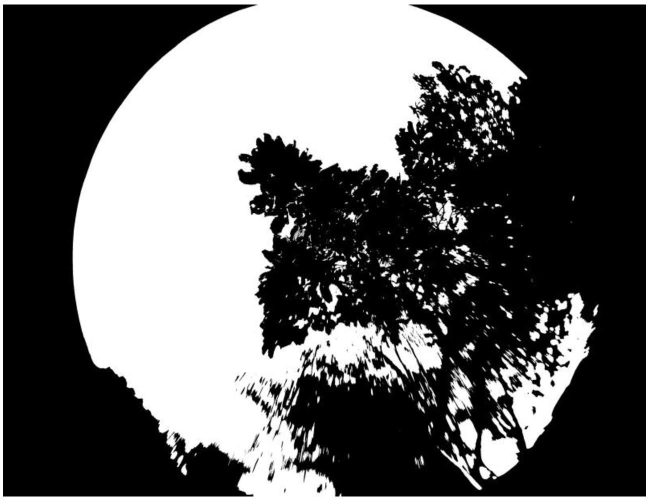

**ID: IMG_20260213_152359.jpg | Sky: P (Gamma: 1.5) | Light: 77.07%**

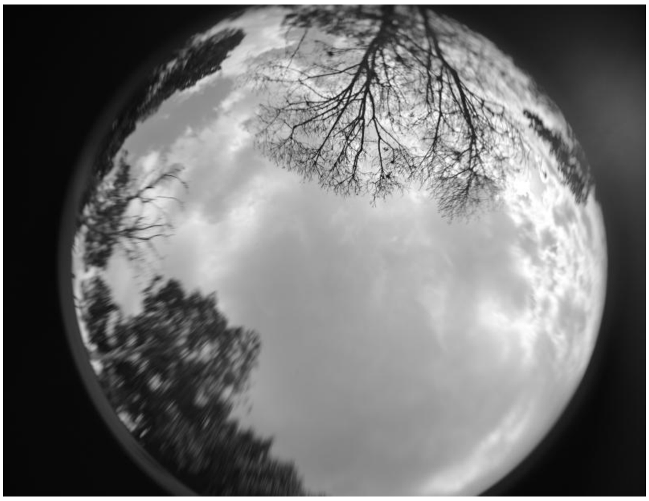

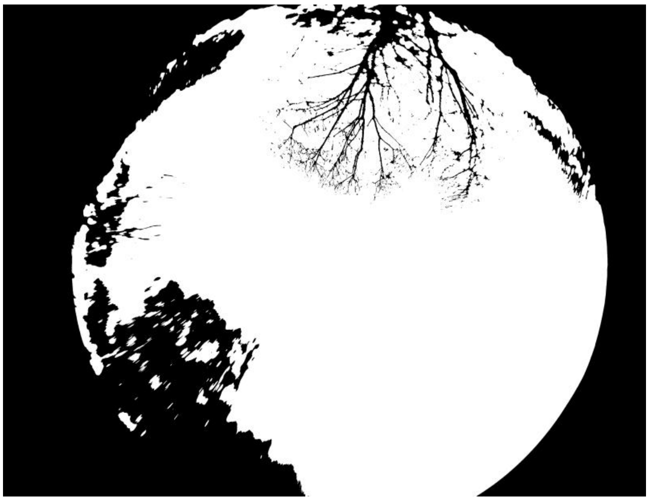

**ID: IMG_20260213_154043.jpg | Sky: P (Gamma: 1.5) | Light: 38.96%**

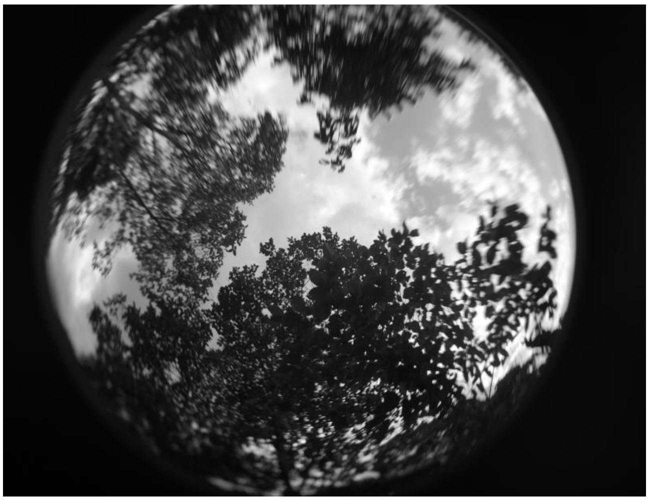

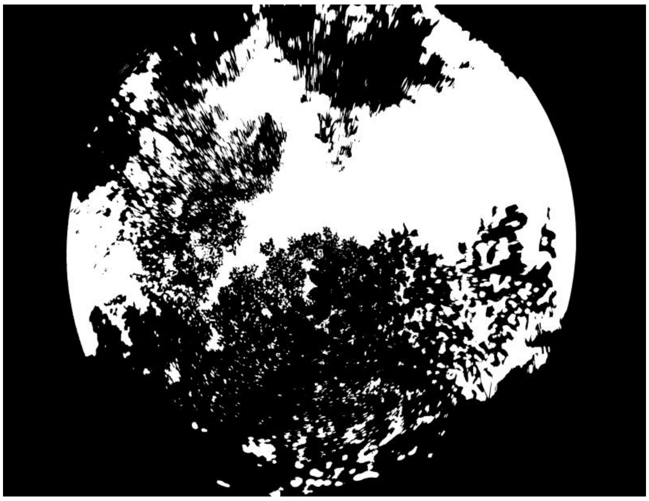

**ID: IMG_20260213_155419.jpg | Sky: P (Gamma: 1.5) | Light: 69.67%**

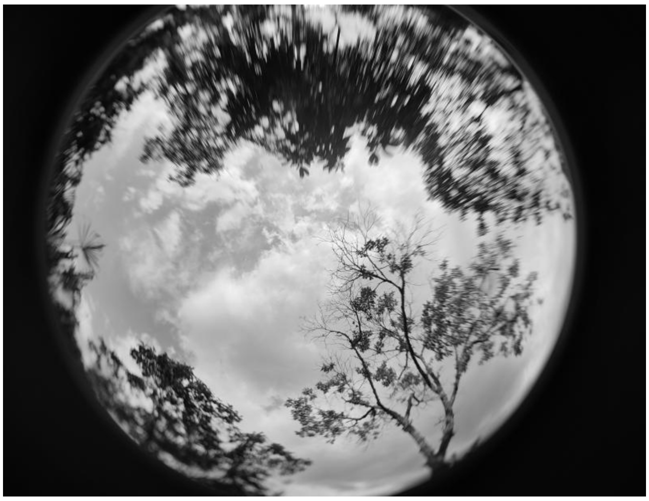

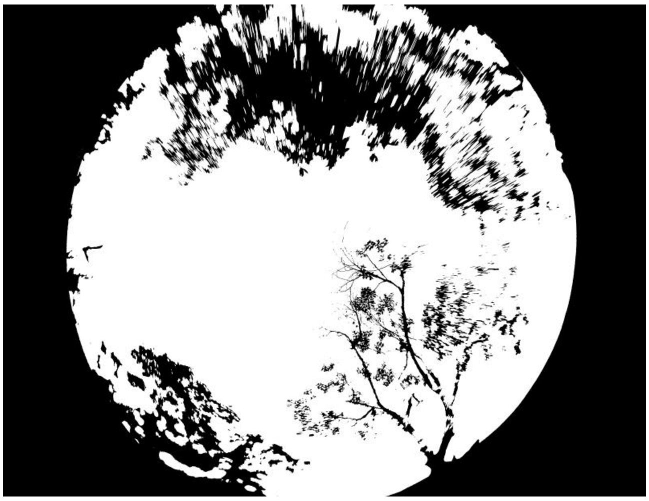

**ID: IMG_20260213_162553.jpg | Sky: P (Gamma: 1.5) | Light: 74.84%**

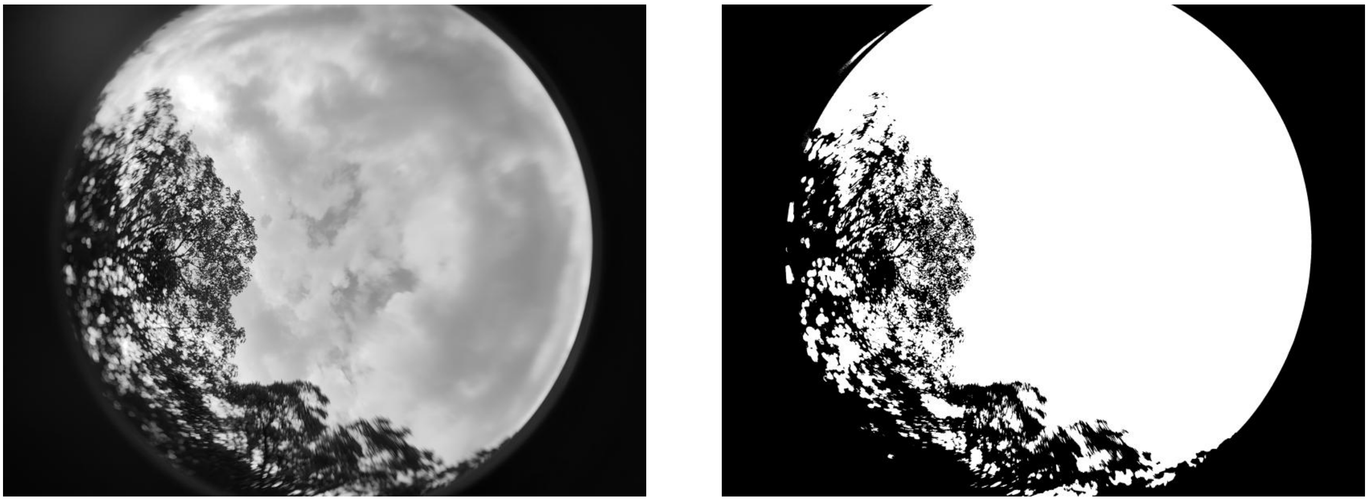

**ID: IMG_20260213_164427.jpg | Sky: C (Gamma: 2.0) | Light: 74.64%**

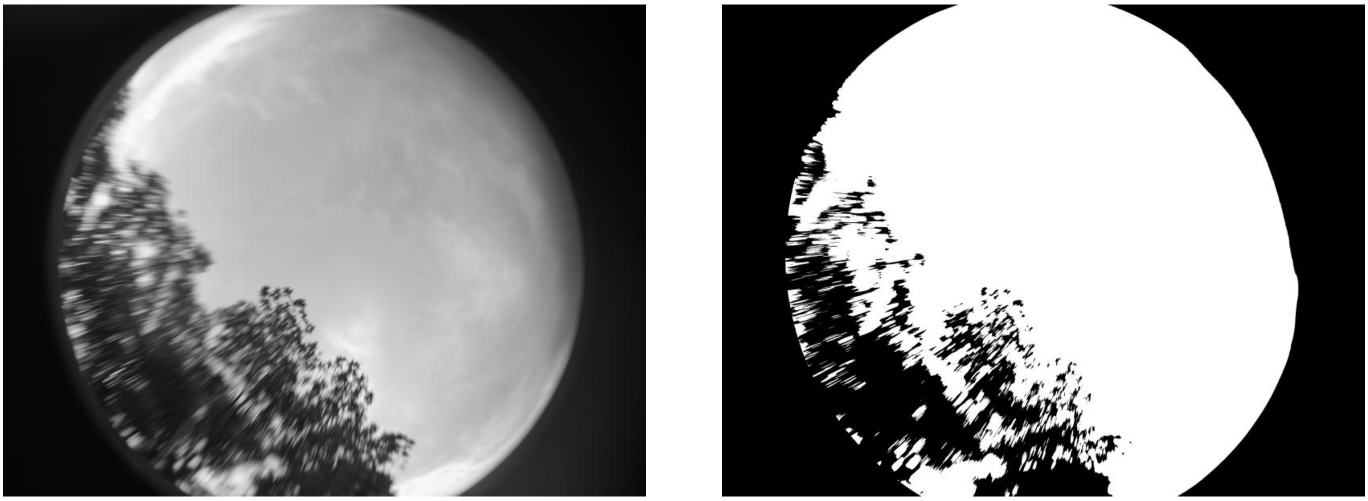

**ID: IMG_20260213_165028.jpg | Sky: C (Gamma: 2.0) | Light: 83.46%**

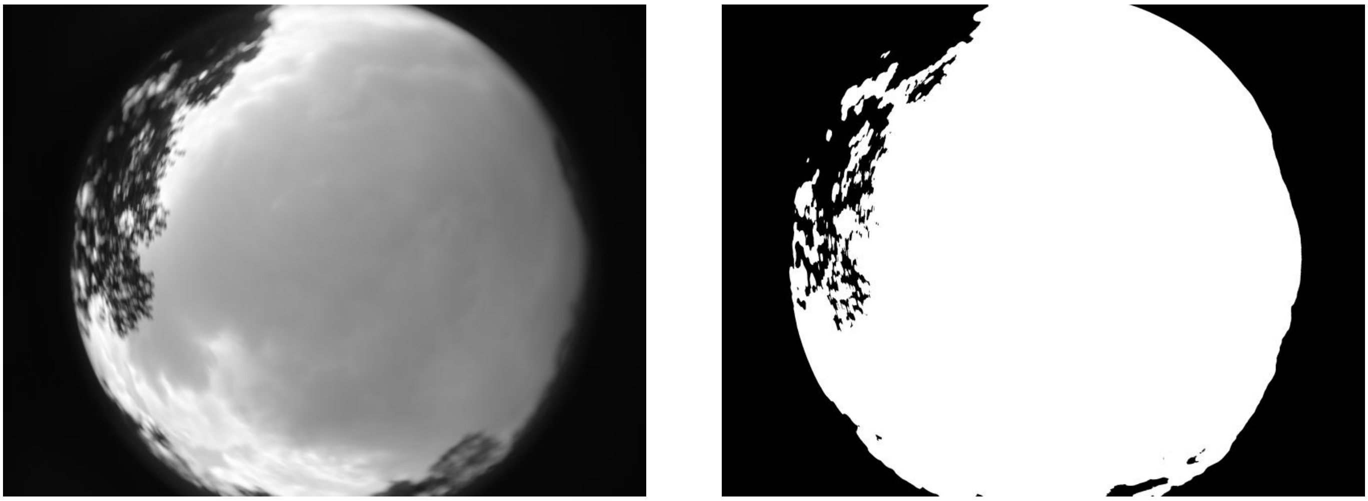

**ID: IMG_20260213_170017.jpg | Sky: C (Gamma: 2.0) | Light: 85.61%**

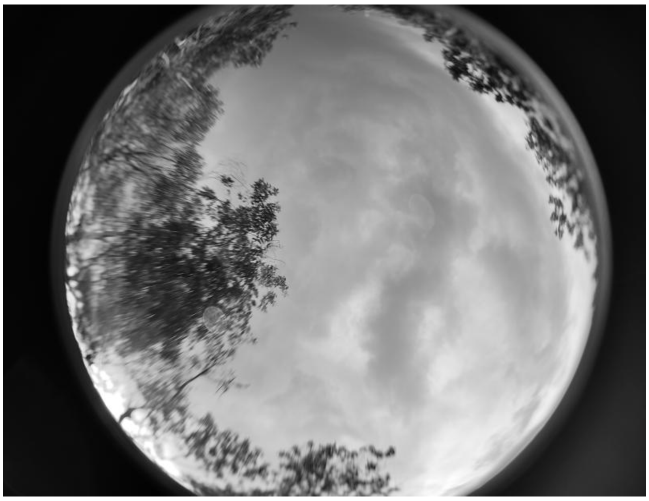

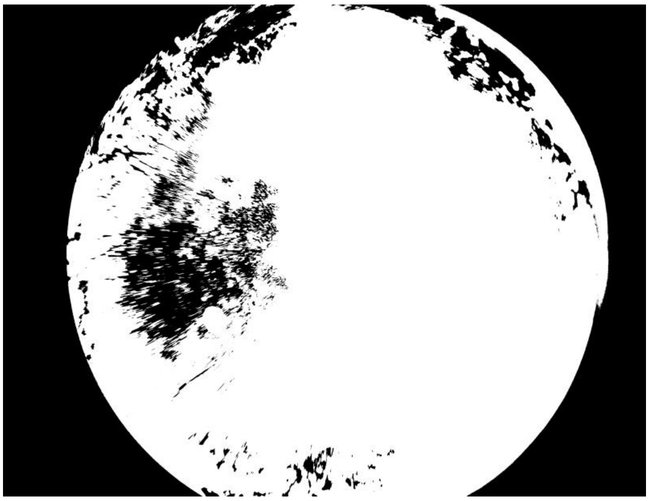

**ID: IMG_20260215_083511.jpg | Sky: P (Gamma: 1.5) | Light: 94.14%**

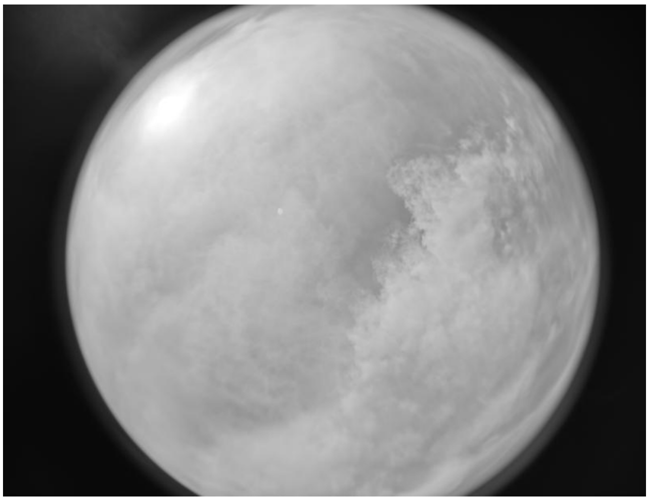

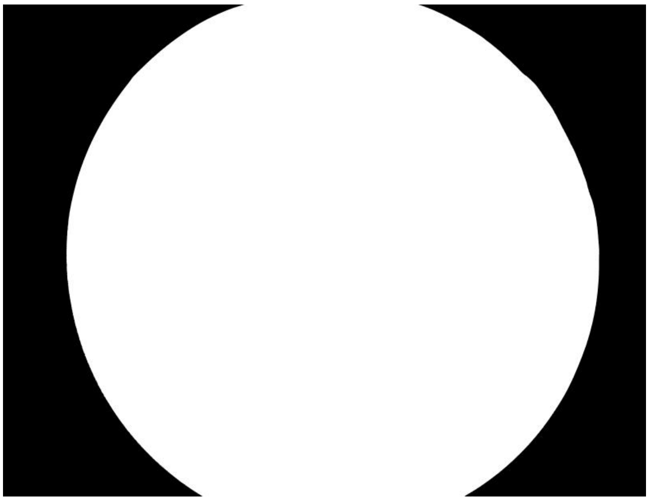

**ID: IMG_20260215_084607.jpg | Sky: P (Gamma: 1.5) | Light: 94.32%**

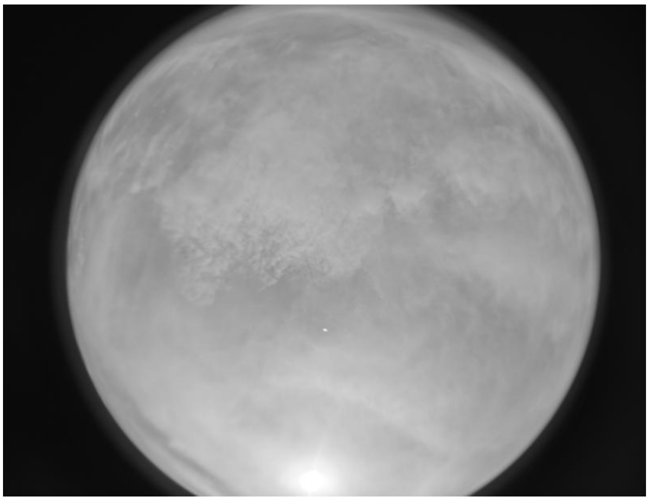

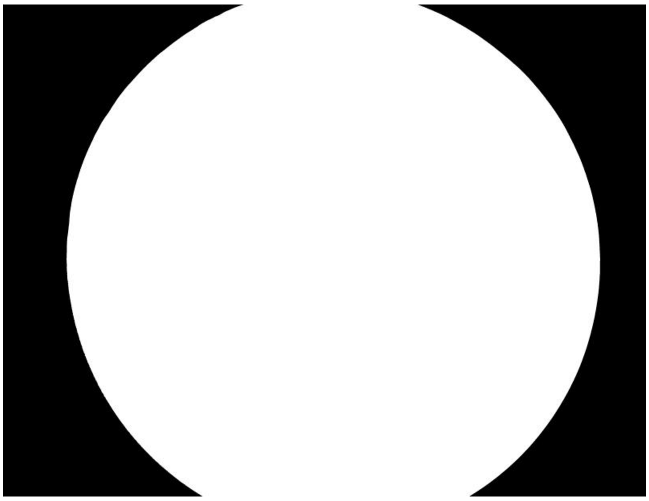

**ID: IMG_20260215_085447.jpg | Sky: P (Gamma: 1.5) | Light: 93.95%**

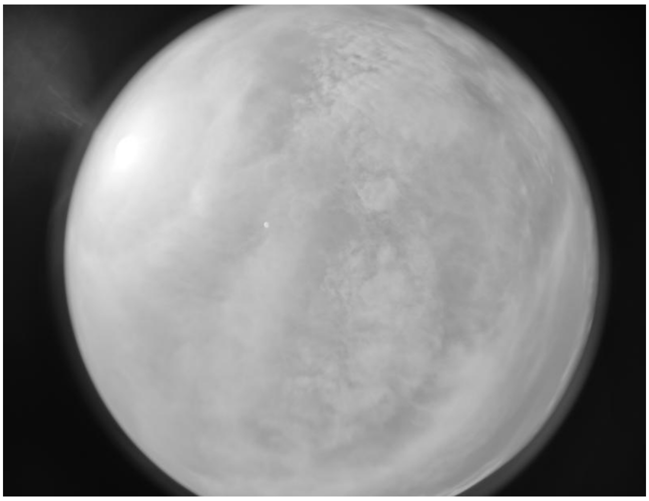
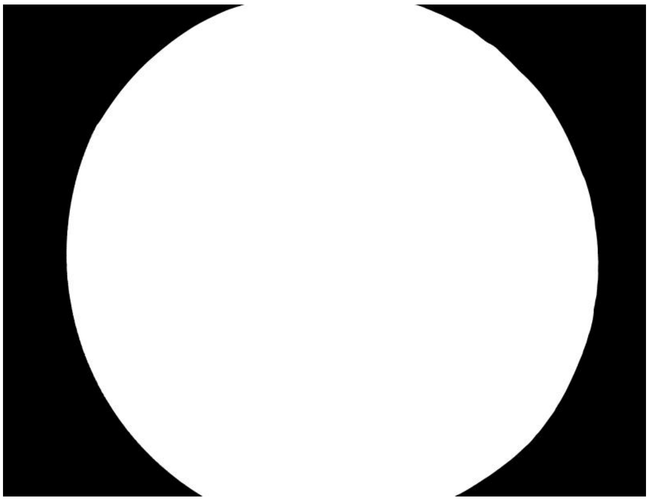

**ID: IMG_20260215_090609.jpg | Sky: P (Gamma: 1.5) | Light: 91.57%**

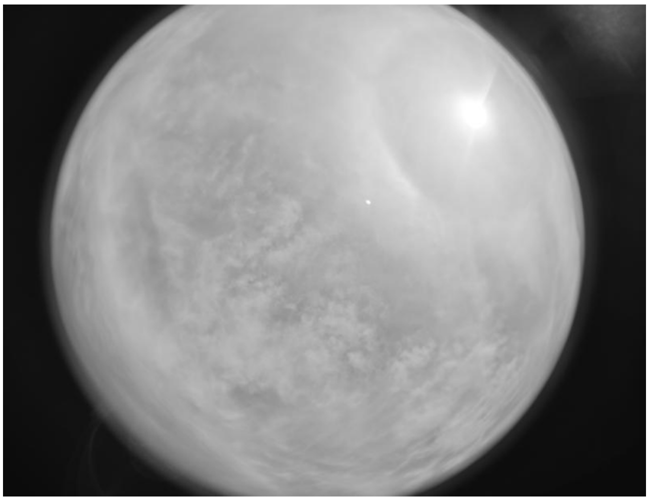
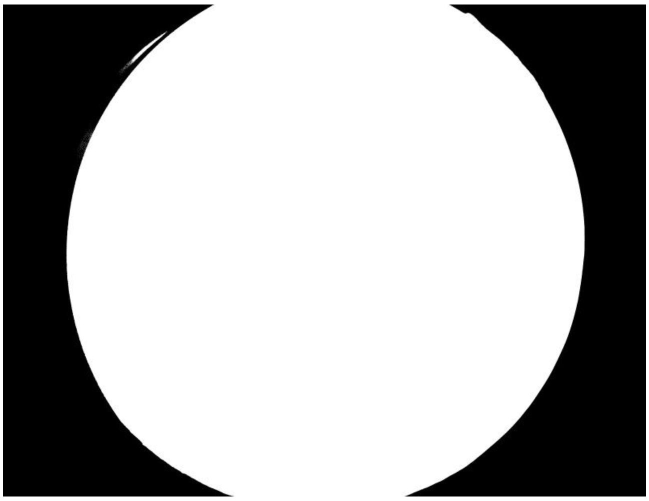

**ID: IMG_20260215_091739.jpg | Sky: P (Gamma: 1.5) | Light: 91.85%**

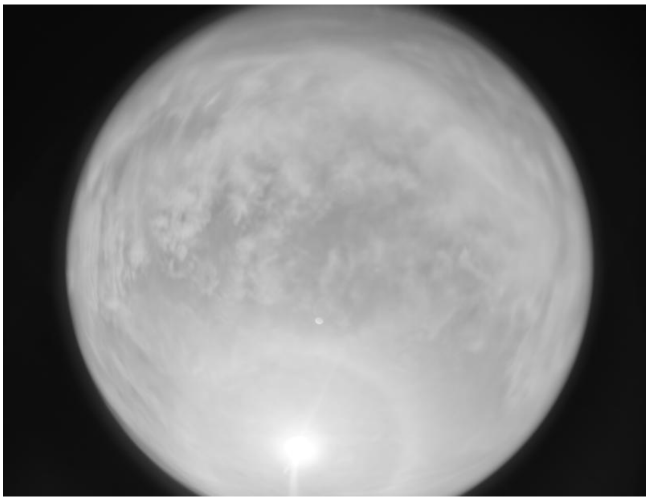
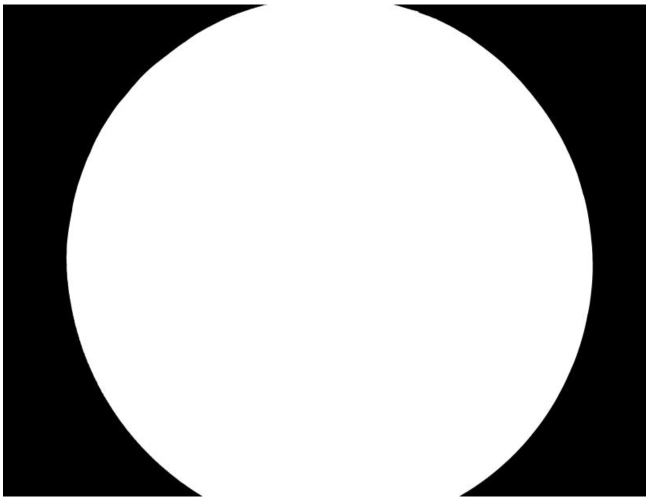

**ID: IMG_20260215_092829.jpg | Sky: C (Gamma: 2.0) | Light: 88.15%**

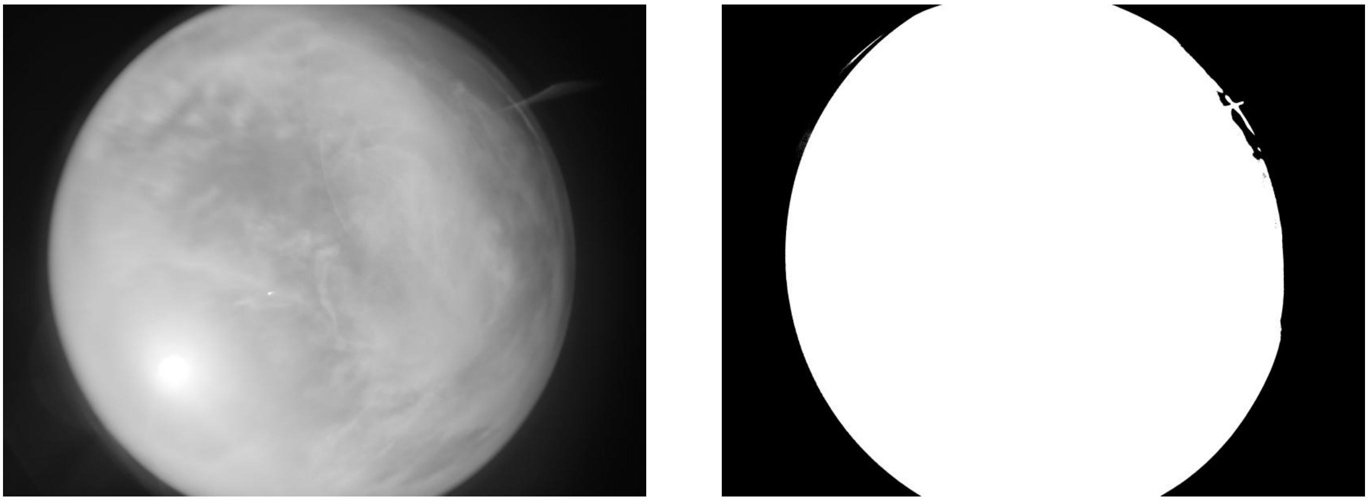

**ID: IMG_20260215_094123.jpg | Sky: C (Gamma: 2.0) | Light: 84.56%**

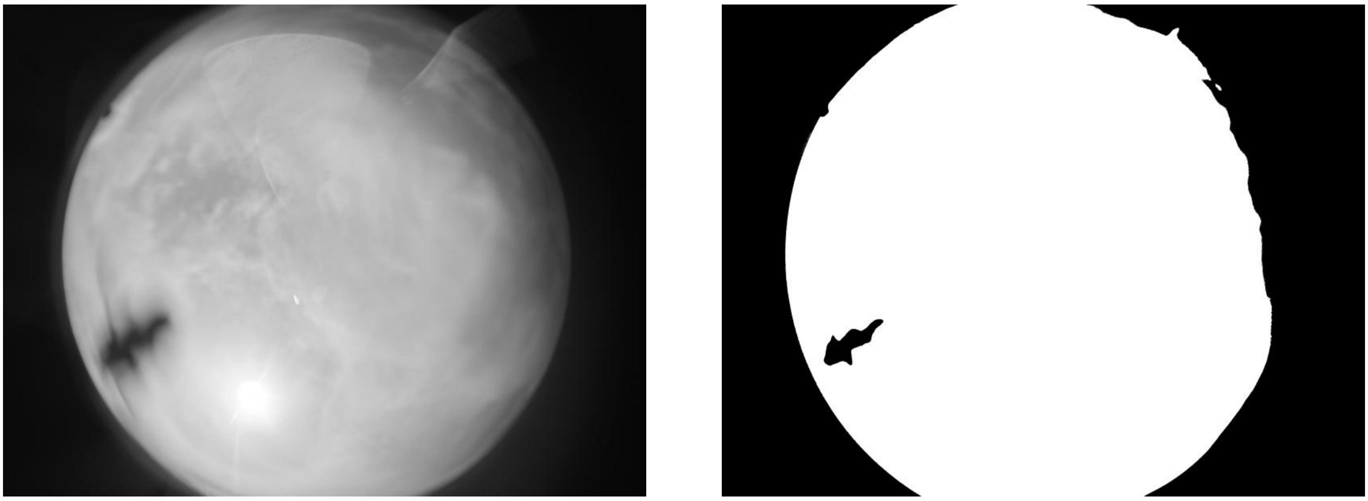

**ID: IMG_20260215_095544.jpg | Sky: P (Gamma: 1.5) | Light: 93.17%**

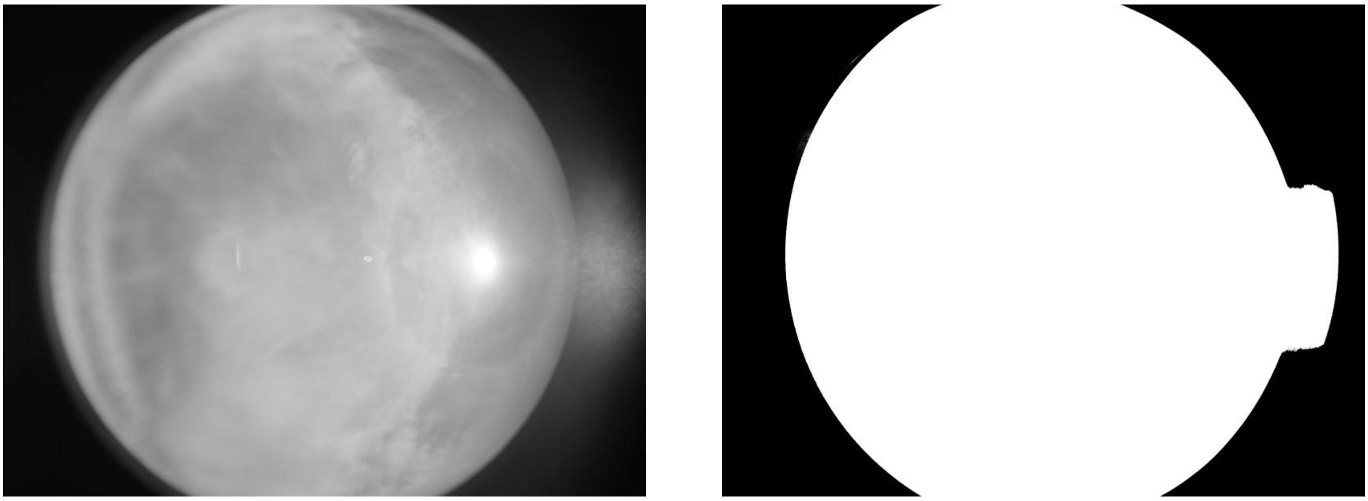

ID: IMG_20260215_101230.jpg | Sky: P (Gamma: 1.5) | Light: 80.34%

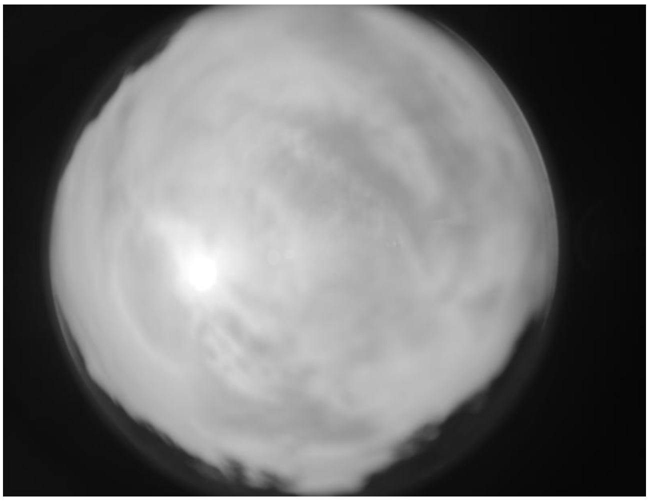

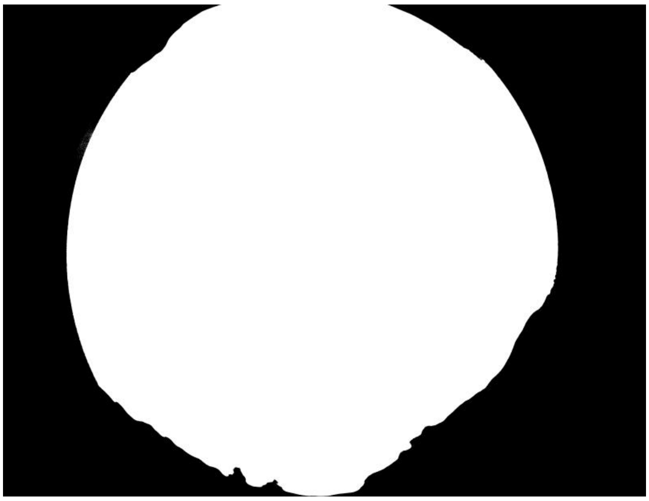

ID: IMG_20260215_103331.jpg | Sky: S (Gamma: 1.0) | Light: 40.29%

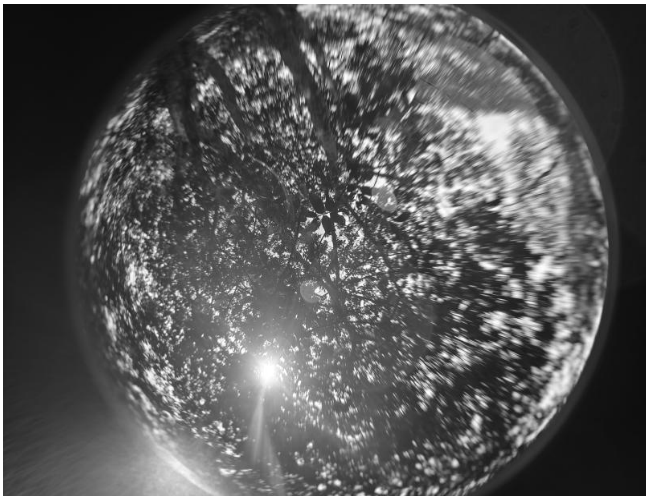

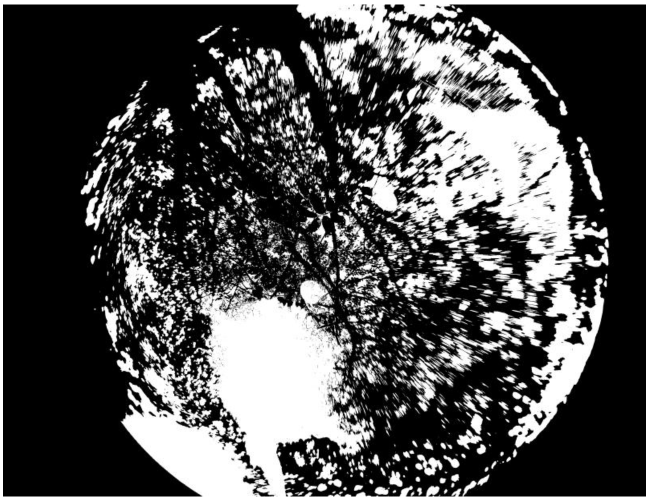

ID: IMG_20260215_110351.jpg | Sky: P (Gamma: 1.5) | Light: 92.12%

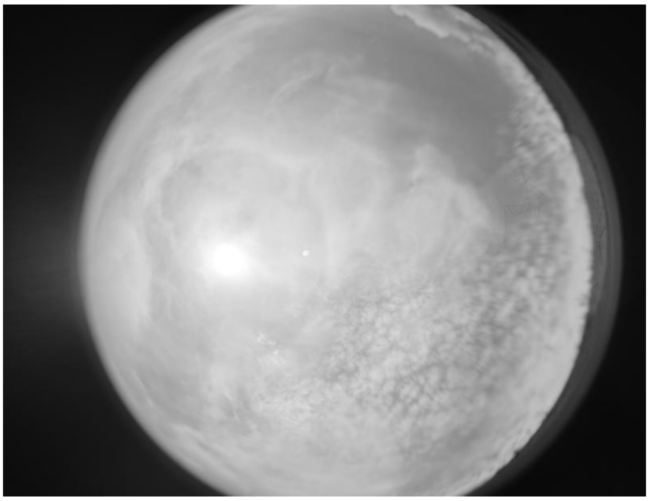

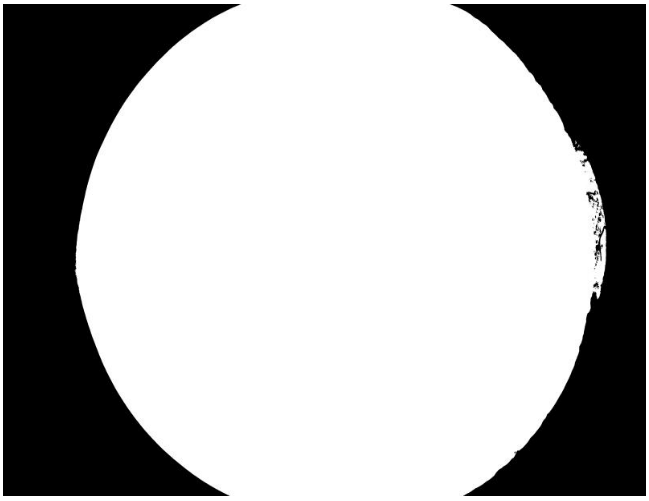

ID: IMG_20260215_112053.jpg | Sky: P (Gamma: 1.5) | Light: 91.62%

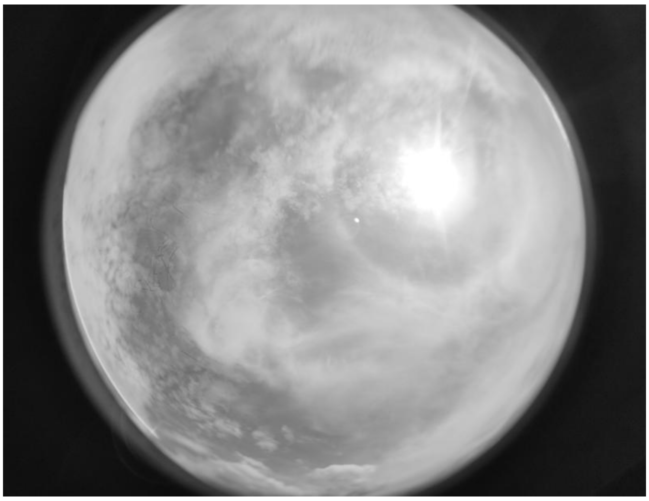

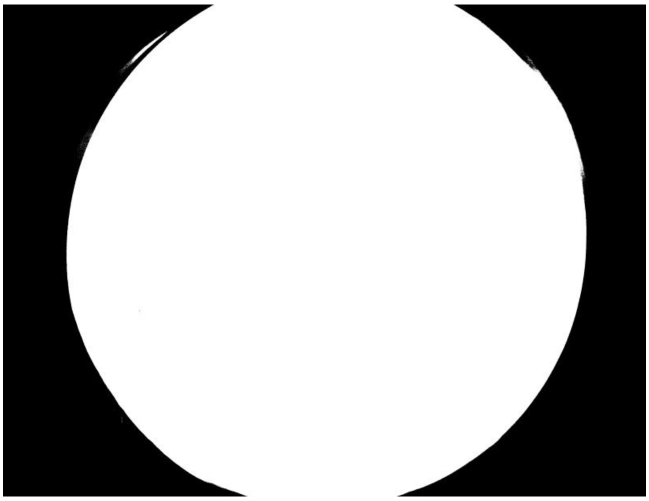

ID: IMG_20260215_115143.jpg | Sky: P (Gamma: 1.5) | Light: 94.17%

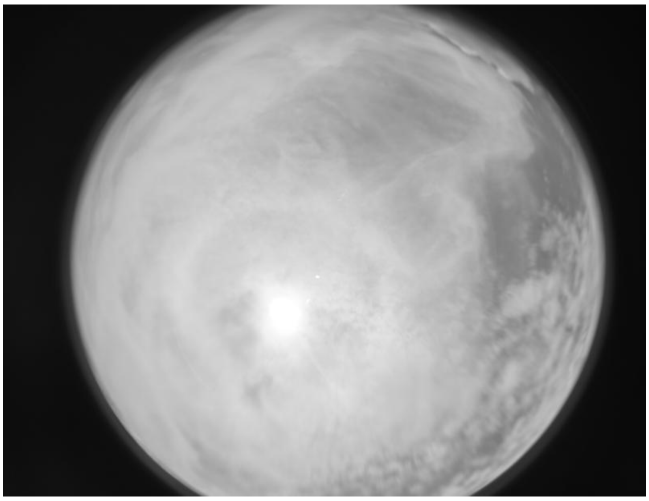

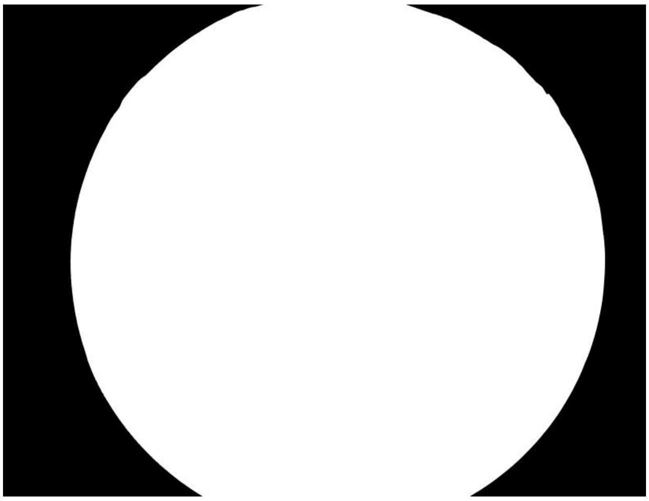

ID: IMG_20260215_121524.jpg | Sky: P (Gamma: 1.5) | Light: 94.79%

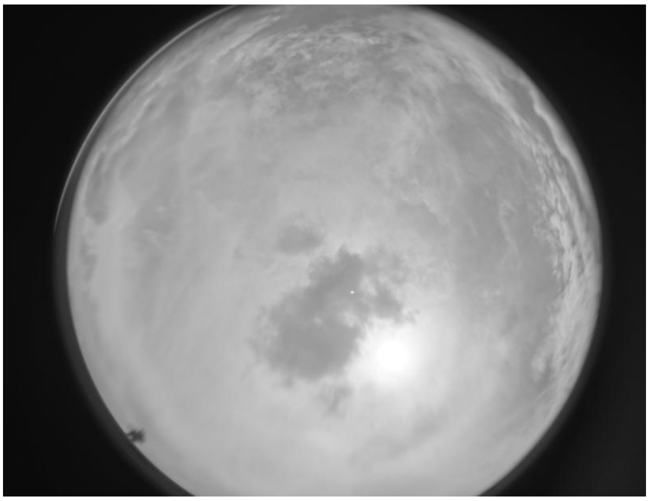

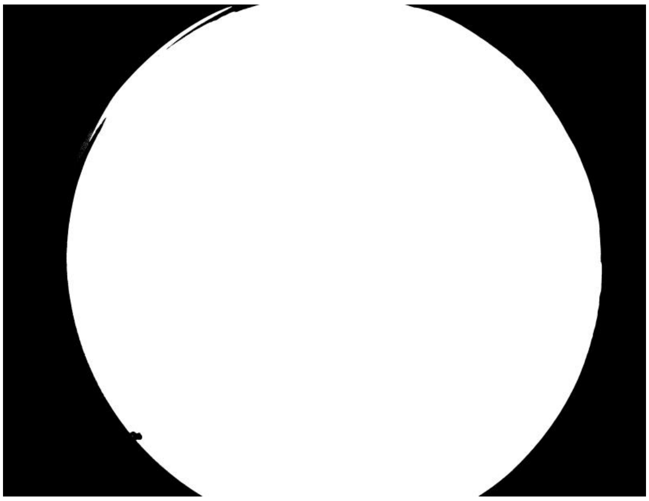

**ID: IMG_20260216_092242.jpg | Sky: S (Gamma: 1.0) | Light: 48.47%**

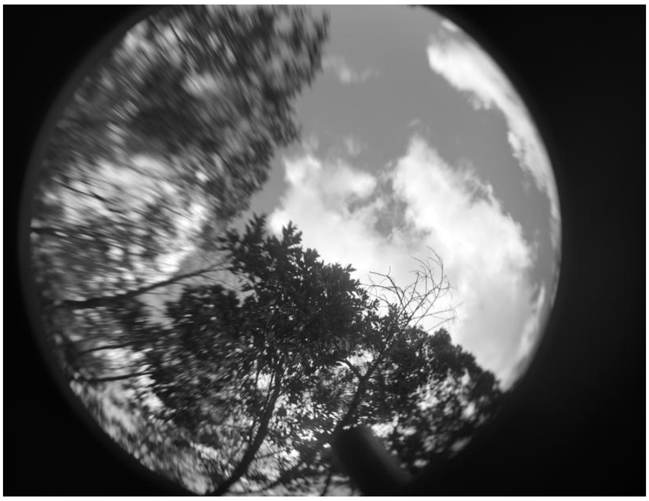

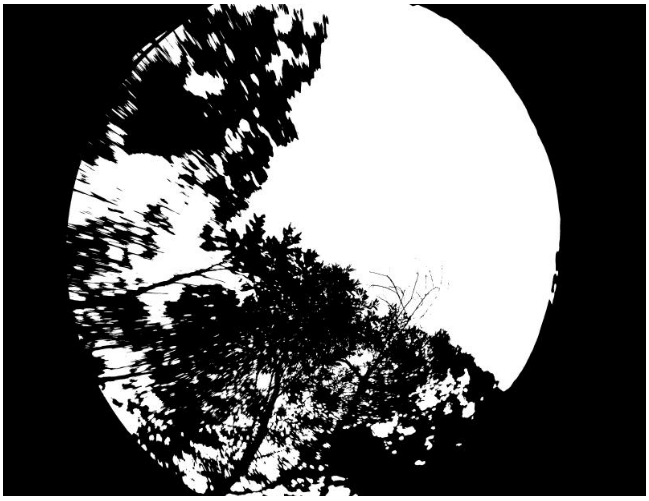

**ID: IMG_20260216_093750.jpg | Sky: S (Gamma: 1.0) | Light: 75.72%**

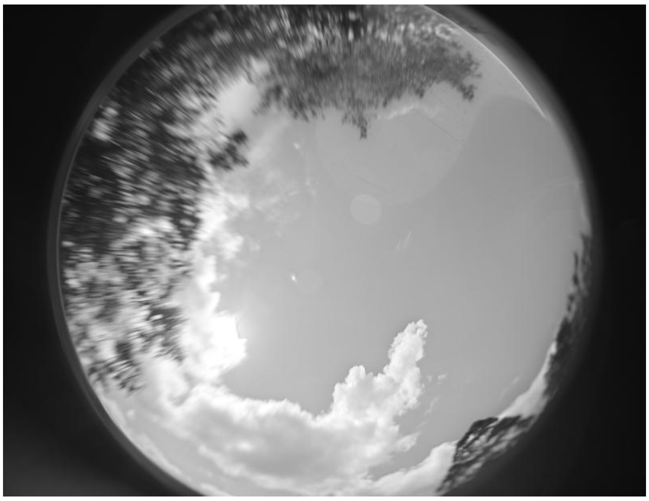

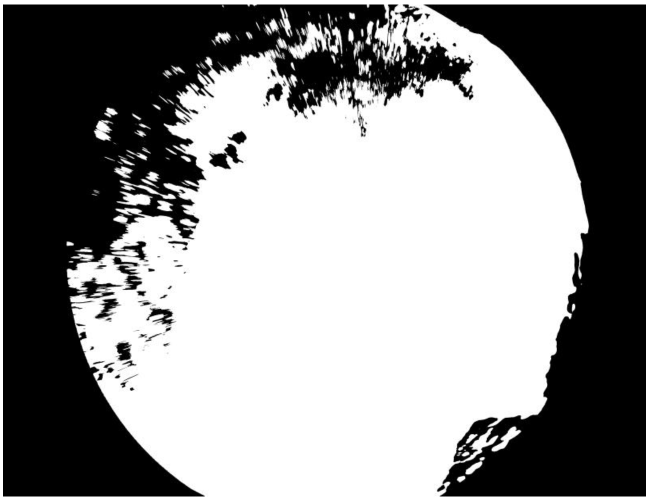

**ID: IMG_20260216_094711.jpg | Sky: P (Gamma: 1.5) | Light: 93.22%**

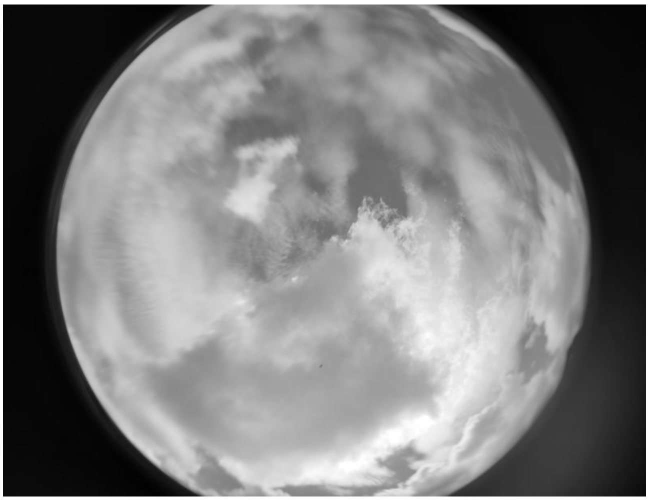

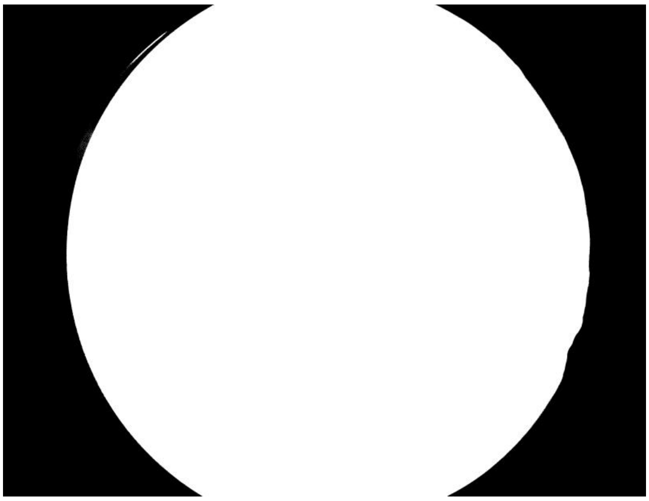

ID: IMG_20260216_100207.jpg | Sky: P (Gamma: 1.5) | Light: 93.0%

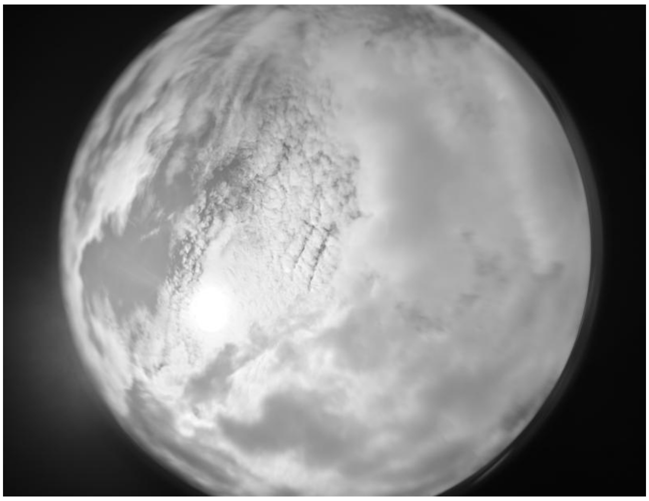

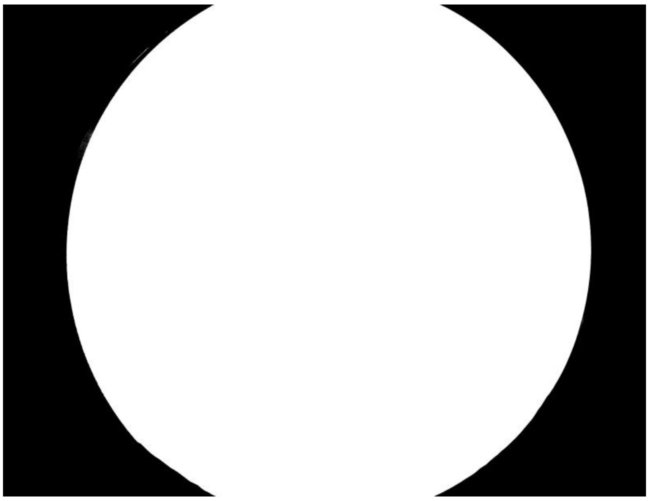

ID: IMG_20260216_100929.jpg | Sky: P (Gamma: 1.5) | Light: 92.37%

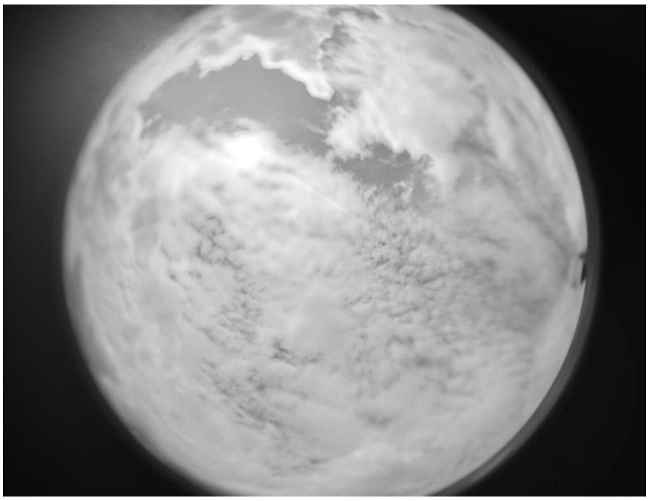

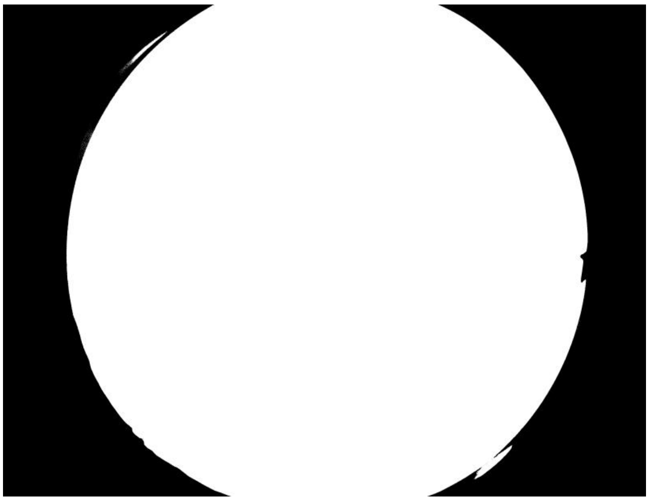

ID: IMG_20260216_101843.jpg | Sky: P (Gamma: 1.5) | Light: 93.16%

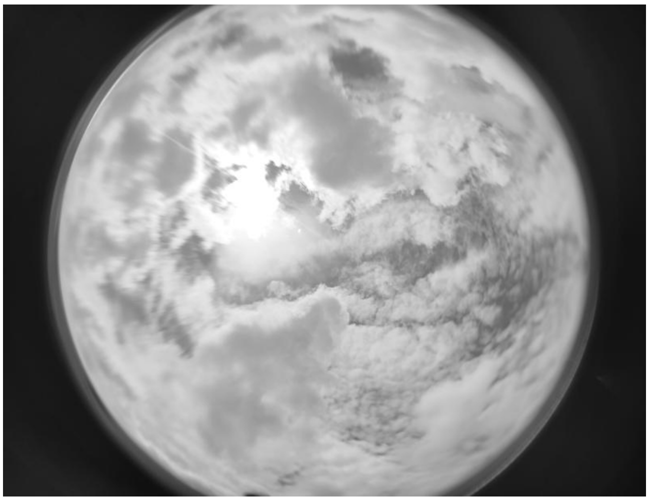

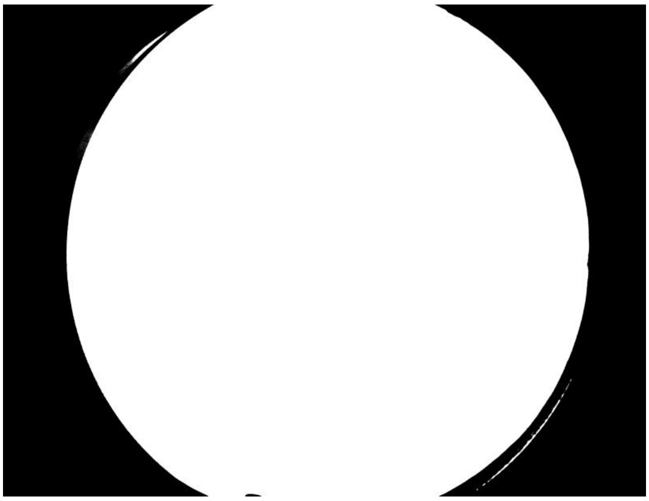

ID: IMG_20260216_103151.jpg | Sky: P (Gamma: 1.5) | Light: 92.61%

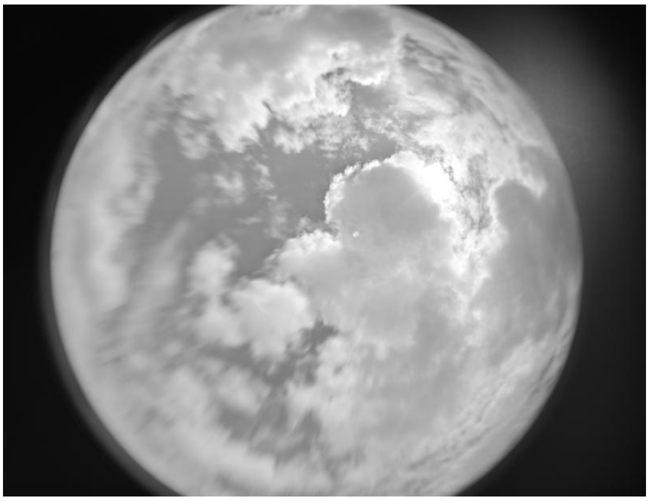

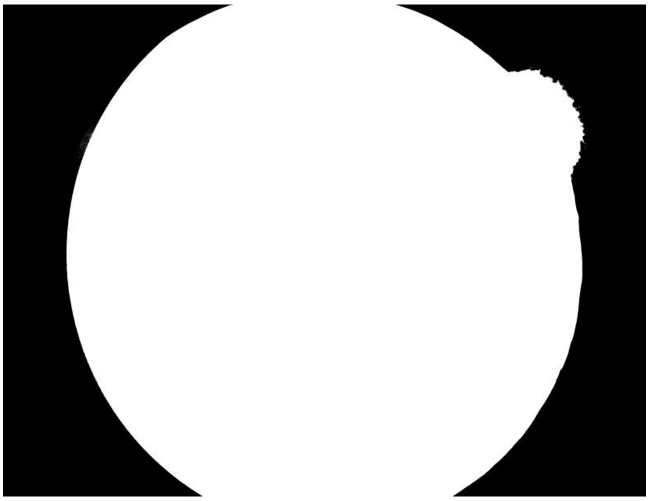

ID: IMG_20260216_104817.jpg | Sky: P (Gamma: 1.5) | Light: 93.51%

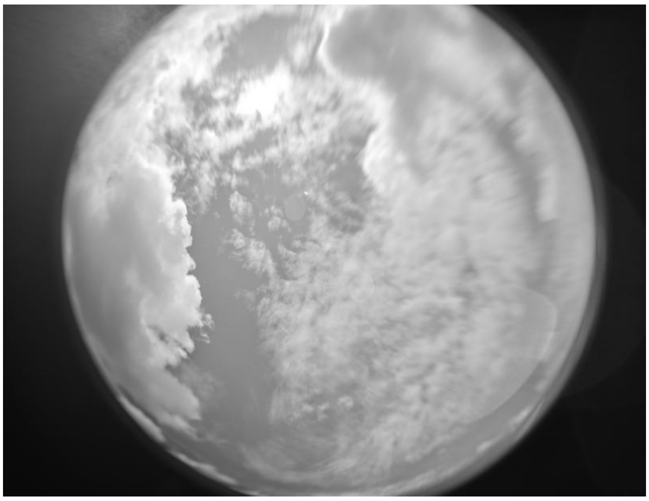

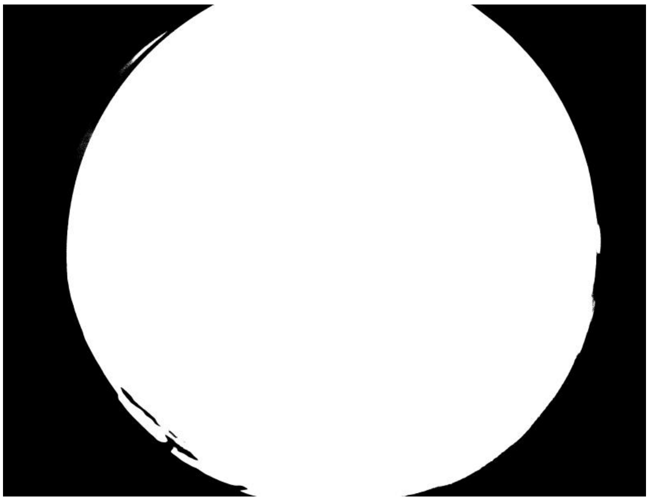

ID: IMG_20260216_110011.jpg | Sky: C (Gamma: 2.0) | Light: 91.36%

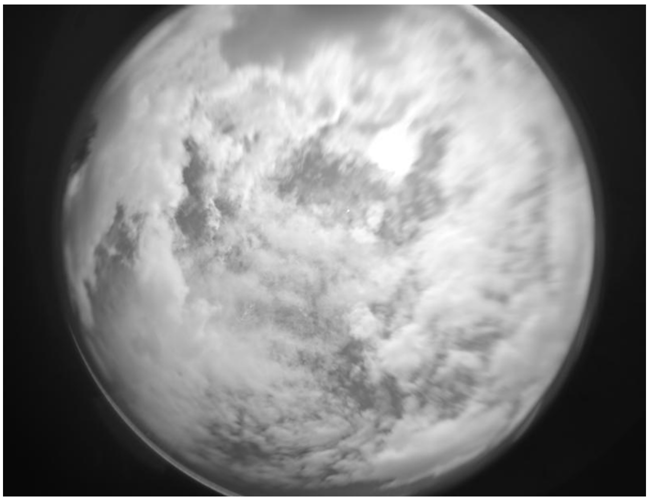

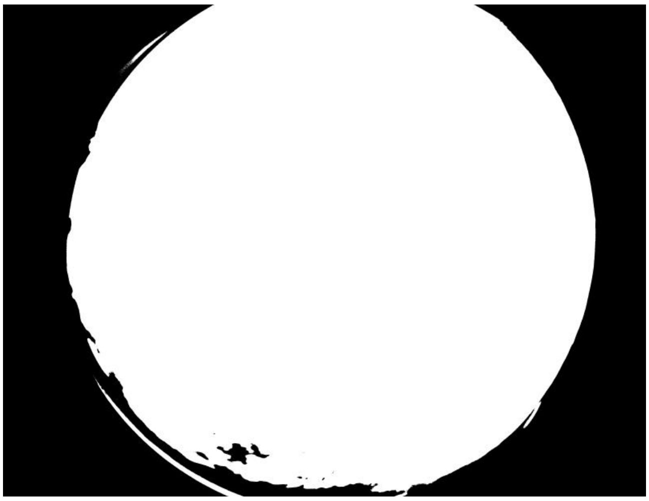

ID: IMG_20260216_111209.jpg | Sky: C (Gamma: 2.0) | Light: 90.22%

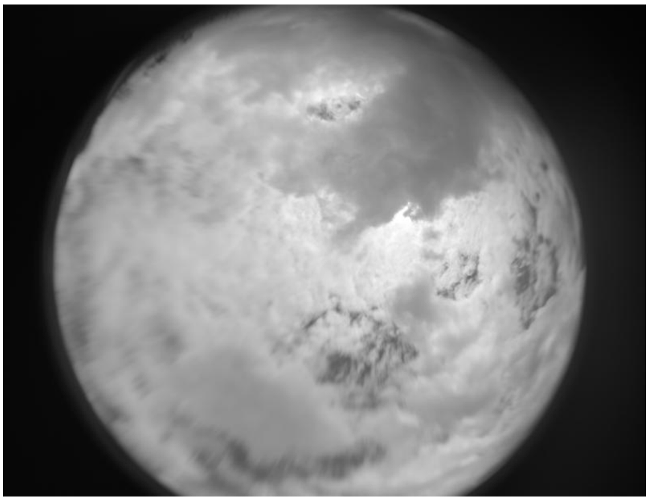

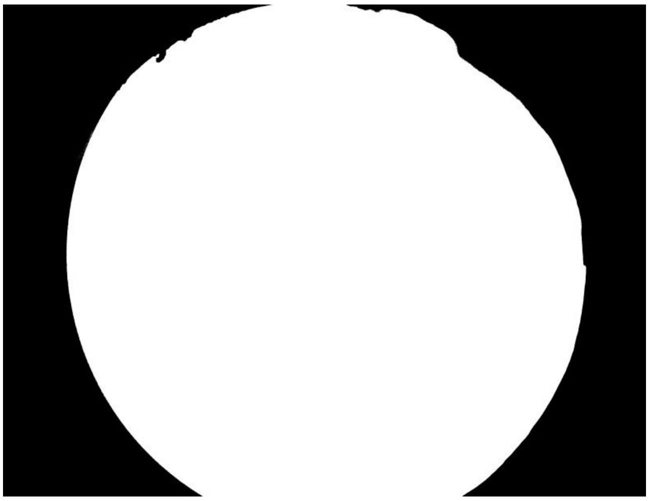

ID: IMG_20260216_112308.jpg | Sky: C (Gamma: 2.0) | Light: 83.18%

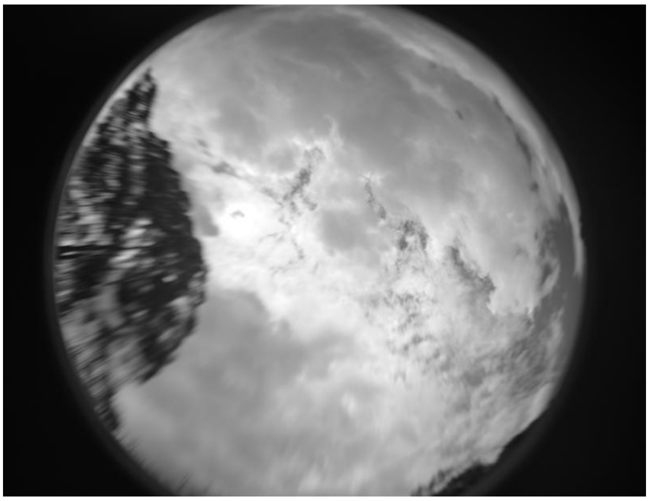

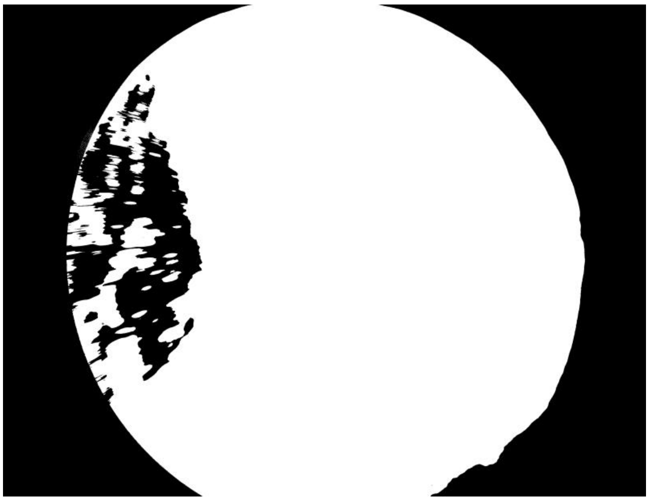

ID: IMG_20260216_112756.jpg | Sky: C (Gamma: 2.0) | Light: 75.33%

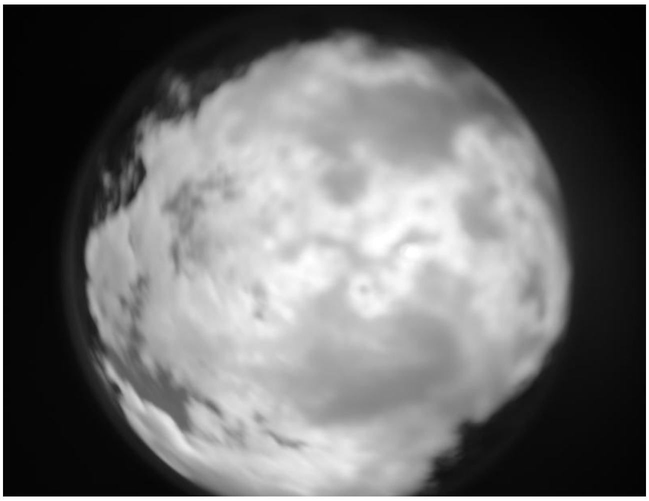

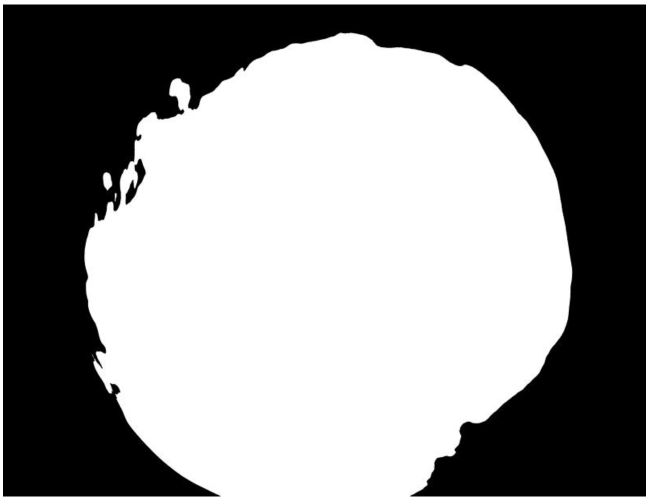

ID: IMG_20260216_143805.jpg | Sky: C (Gamma: 2.0) | Light: 70.35%

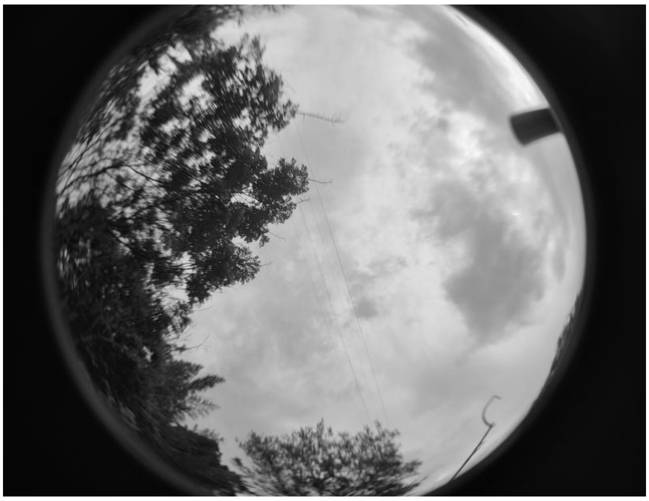
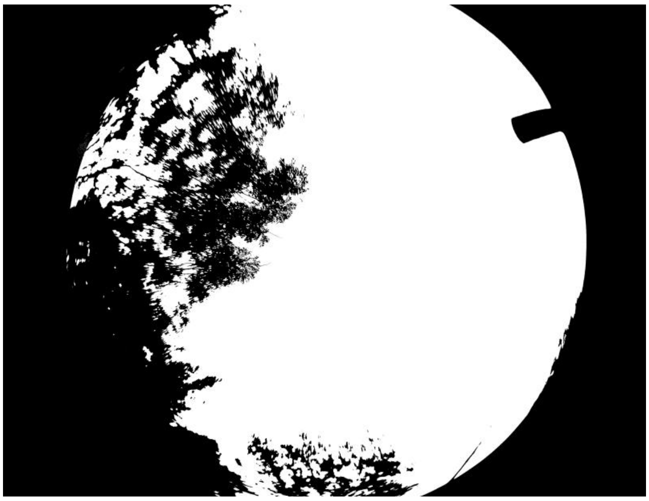

ID: IMG_20260216_144856.jpg | Sky: C (Gamma: 2.0) | Light: 77.08%

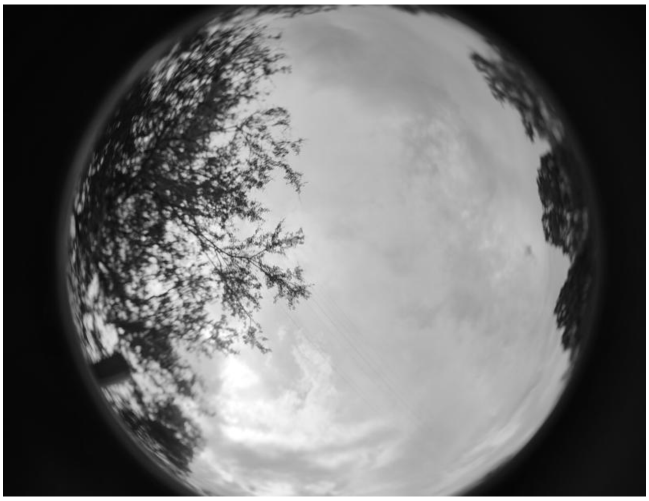
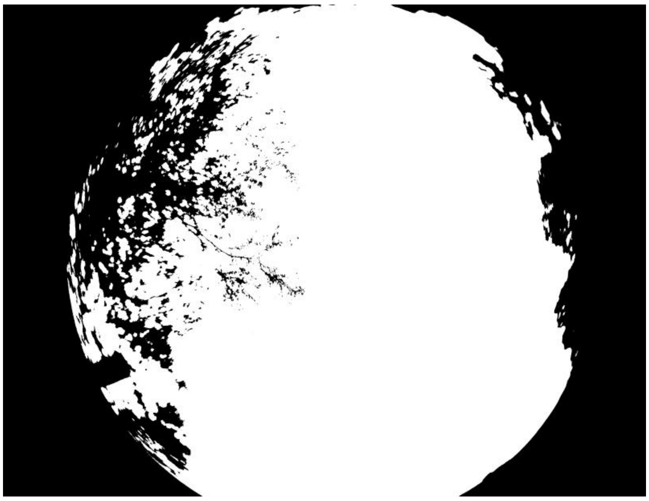

ID: IMG_20260217_081740.jpg | Sky: S (Gamma: 1.0) | Light: 93.72%

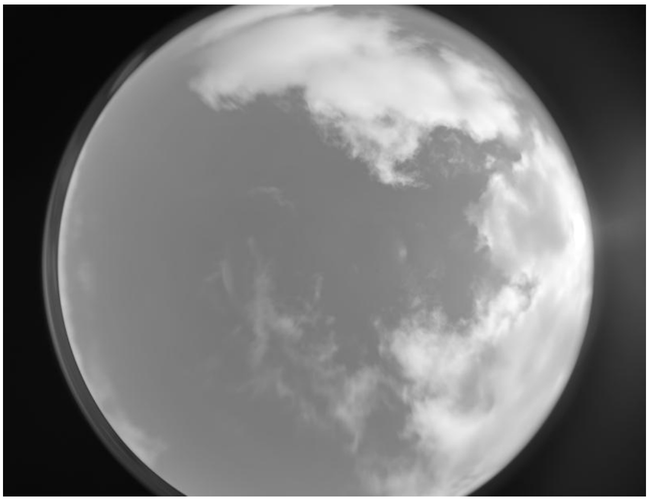
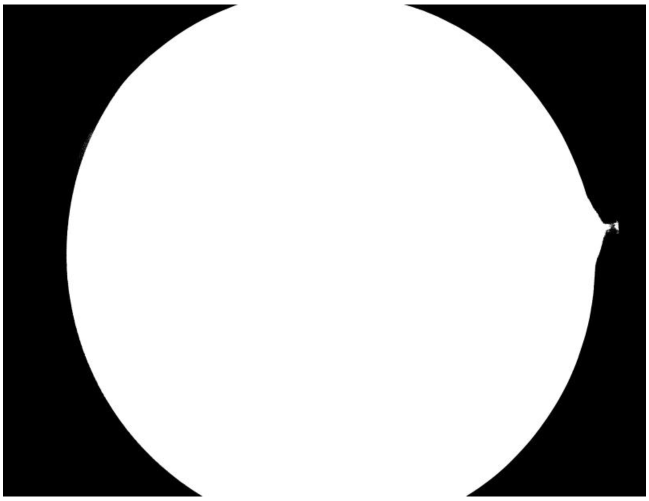

ID: IMG_20260217_082146.jpg | Sky: S (Gamma: 1.0) | Light: 91.64%

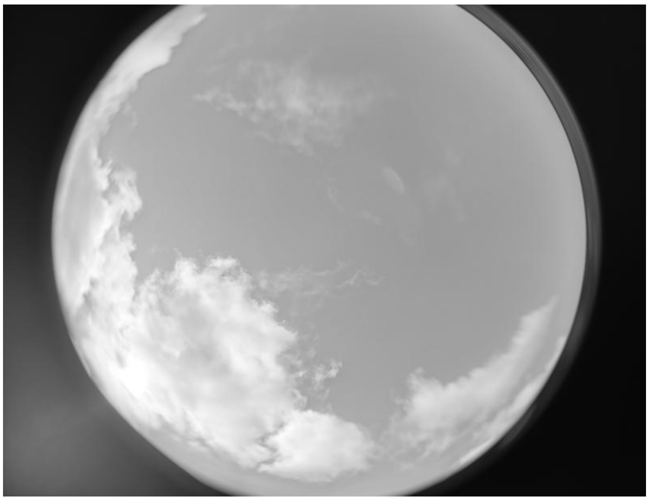

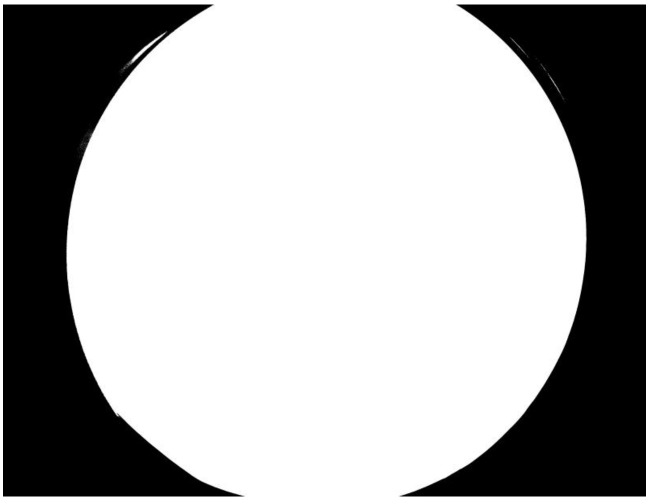

ID: IMG_20260217_084438.jpg | Sky: S (Gamma: 1.0) | Light: 92.79%

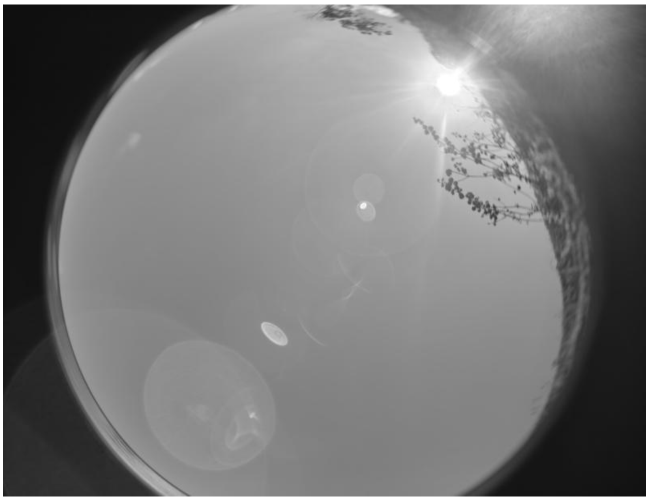

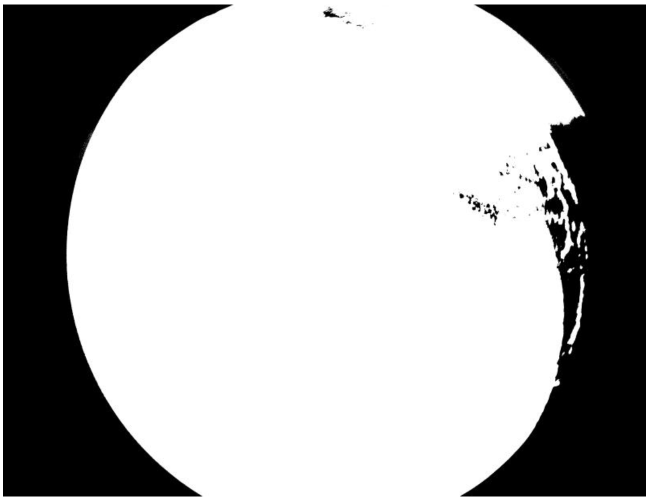

ID: IMG_20260217_085511.jpg | Sky: S (Gamma: 1.0) | Light: 91.48%

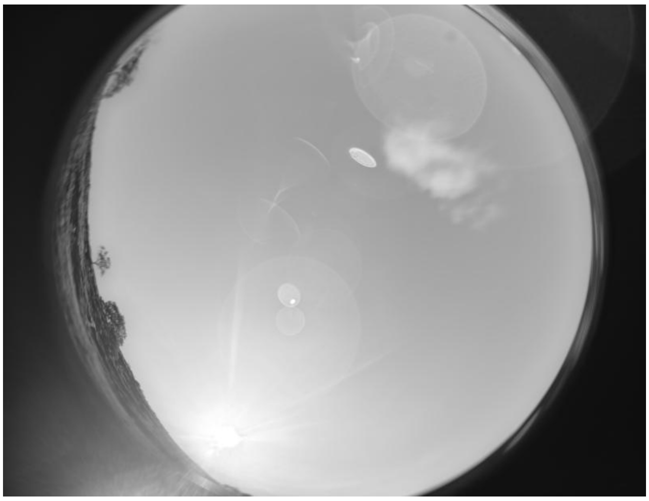

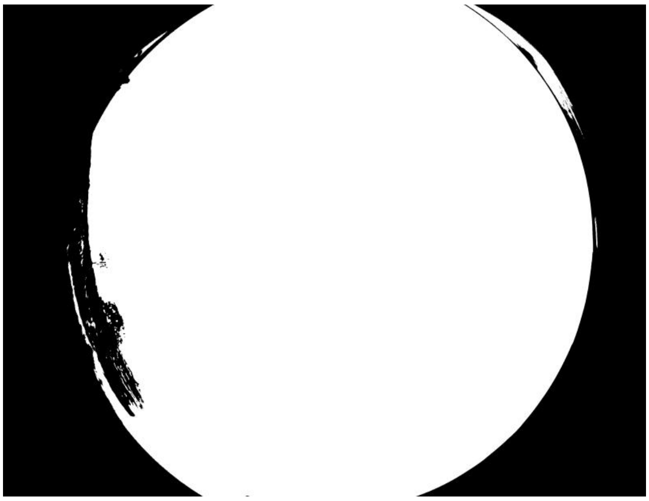

**ID: IMG_20260217_090650.jpg | Sky: S (Gamma: 1.0) | Light: 85.19%**

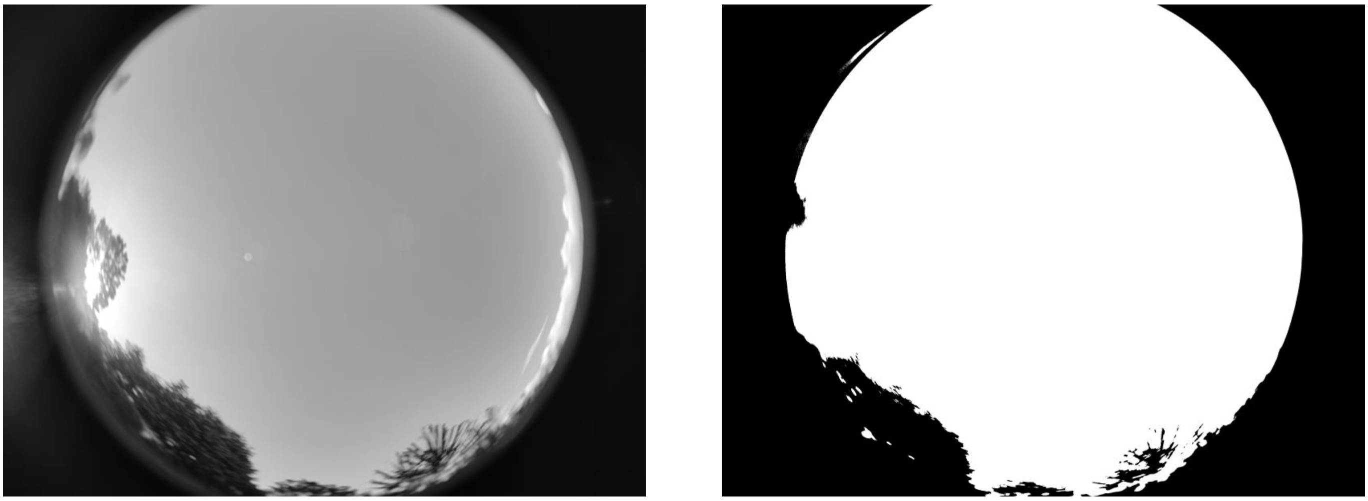

**ID: IMG_20260217_093256.jpg | Sky: S (Gamma: 1.0) | Light: 91.05%**

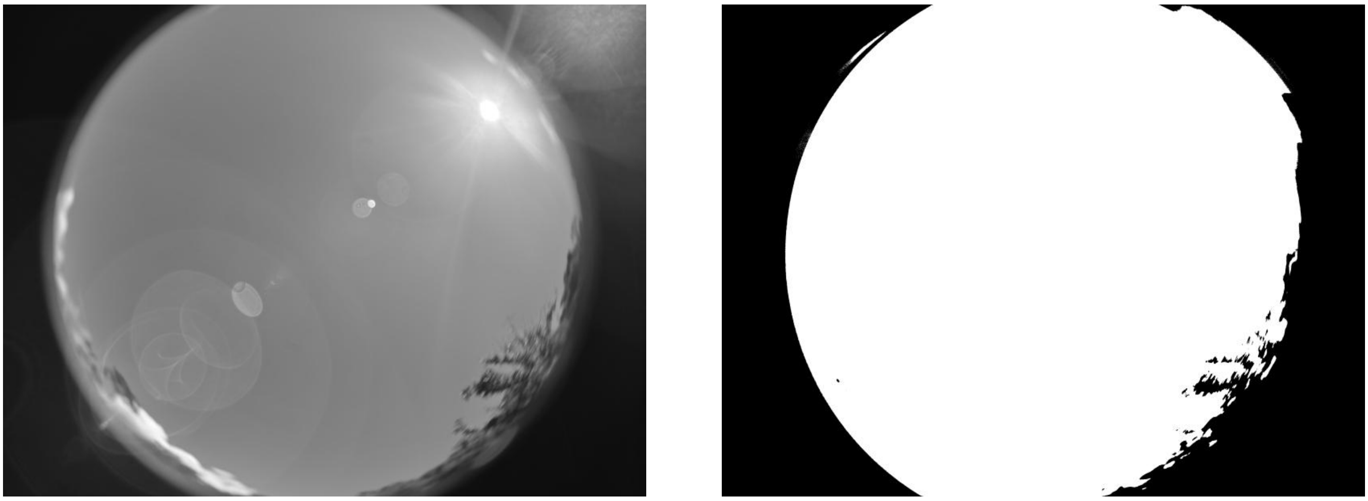

**ID: IMG_20260217_094515.jpg | Sky: S (Gamma: 1.0) | Light: 79.97%**

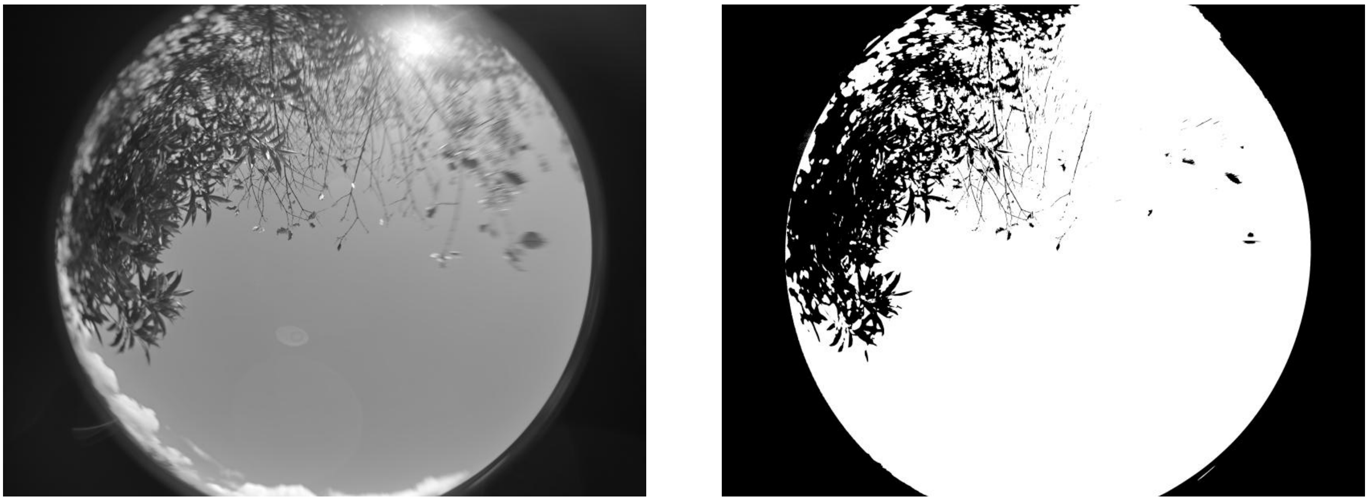

**ID: IMG_20260217_100300.jpg | Sky: S (Gamma: 1.0) | Light: 63.36%**

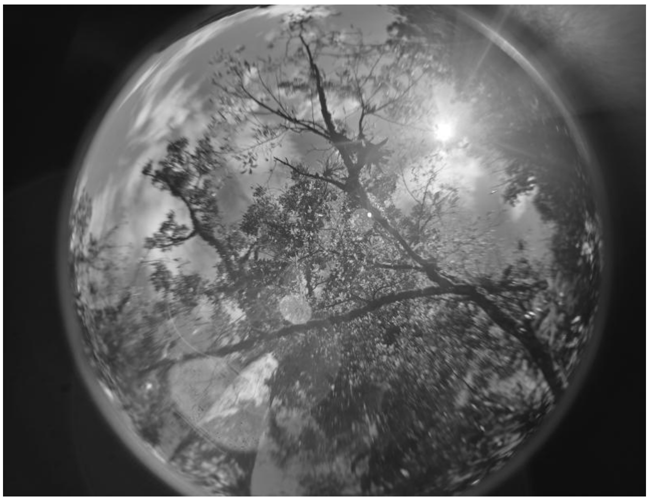

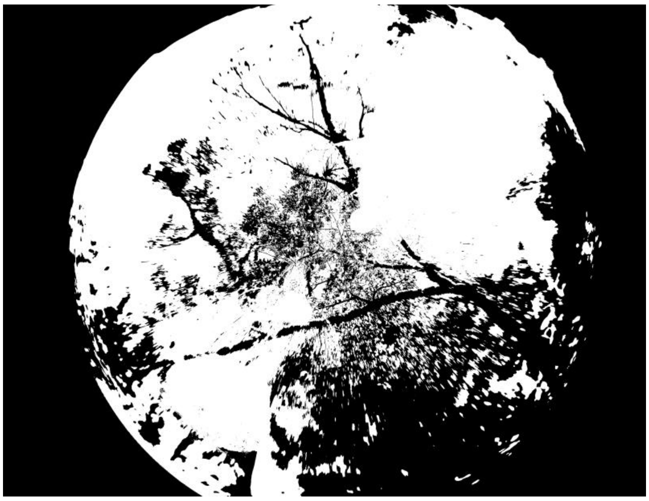

**ID: IMG_20260217_101746.jpg | Sky: S (Gamma: 1.0) | Light: 93.75%**

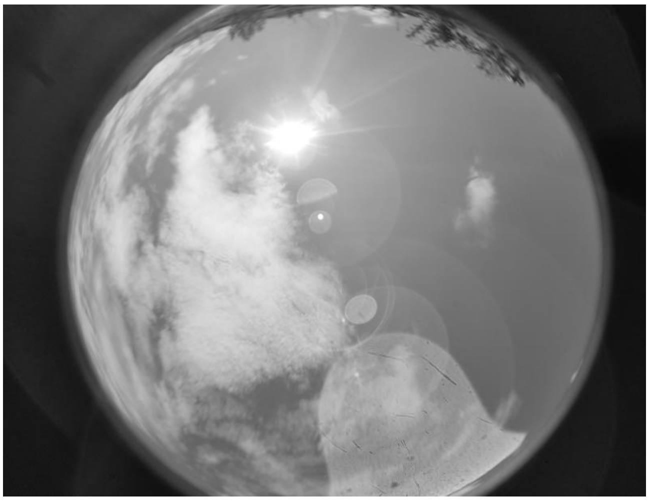

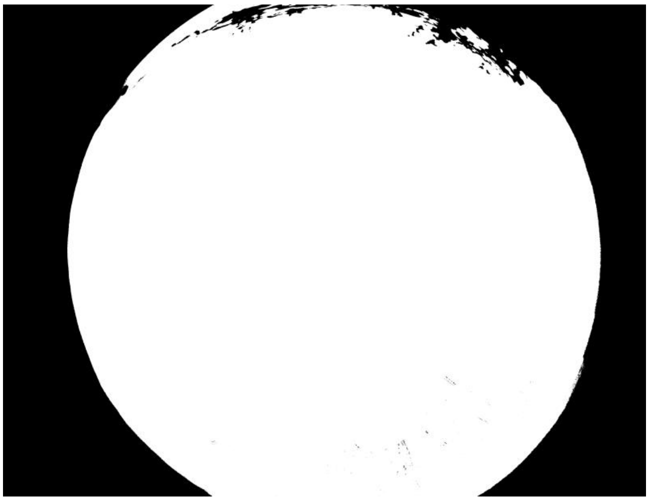

**ID: IMG_20260217_104109.jpg | Sky: S (Gamma: 1.0) | Light: 47.69%**

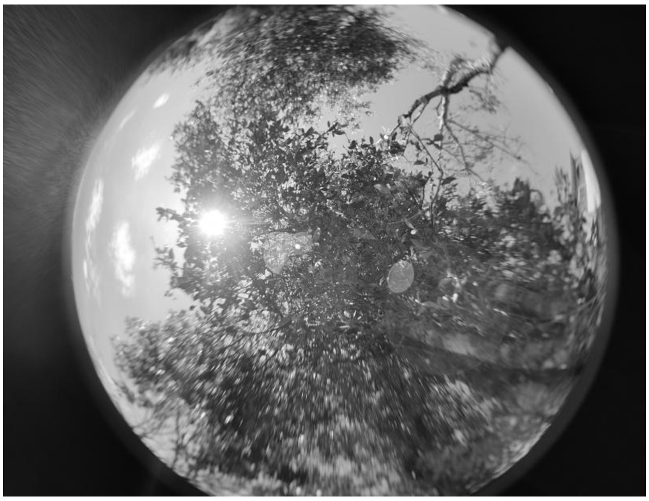

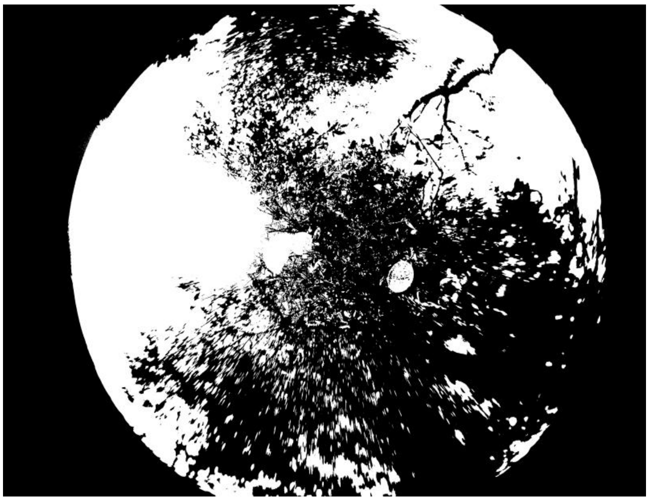

**ID: IMG_20260217_105701.jpg | Sky: S (Gamma: 1.0) | Light: 84.78%**

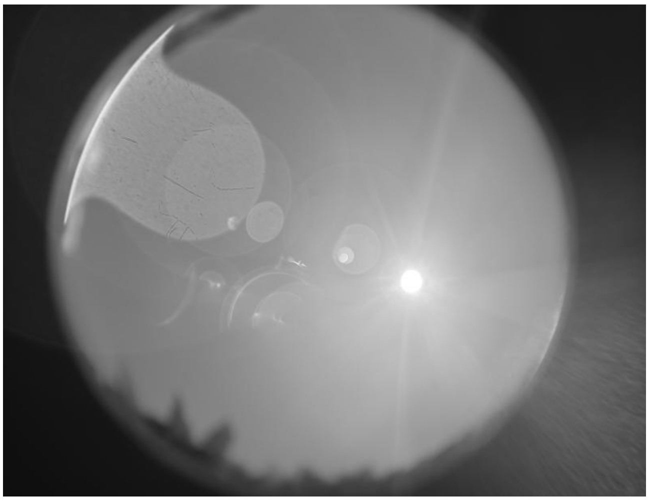

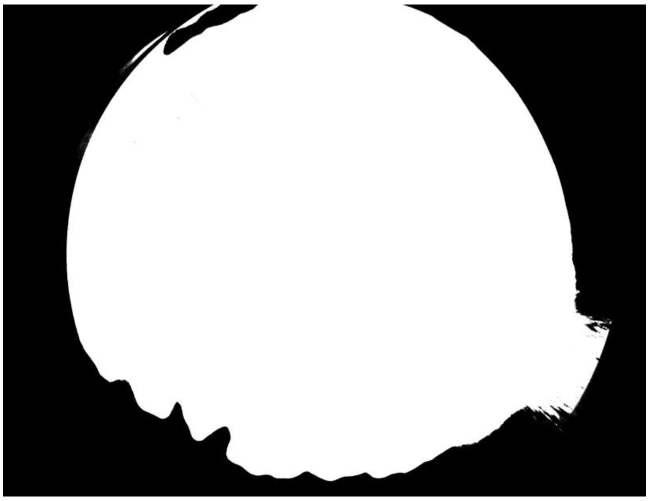

**ID: IMG_20260217_110718.jpg | Sky: S (Gamma: 1.0) | Light: 95.9%**

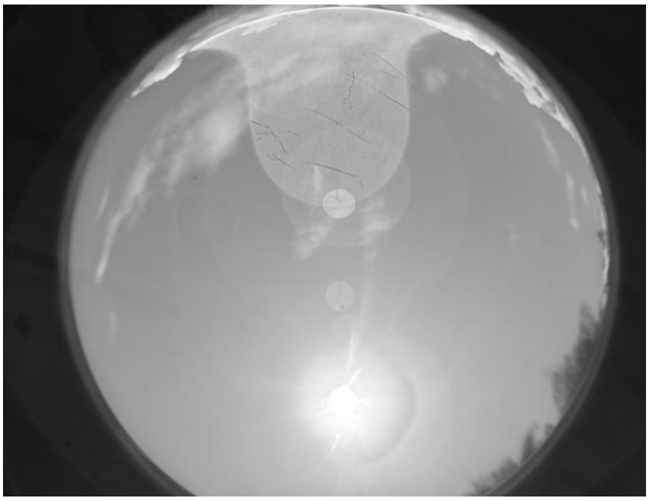

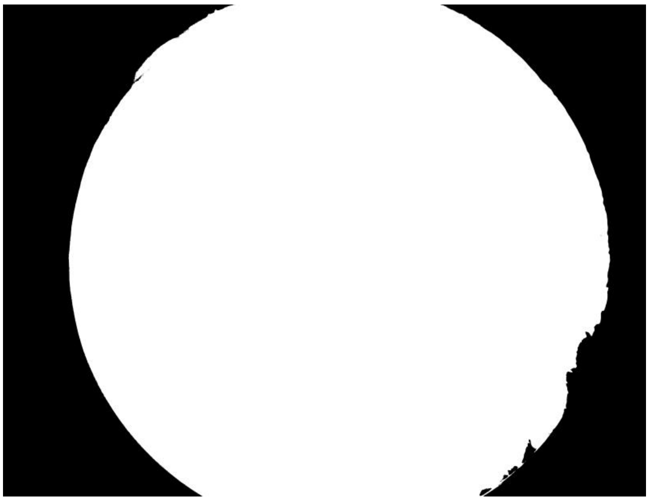

**ID: IMG_20260217_112046.jpg | Sky: S (Gamma: 1.0) | Light: 54.6%**

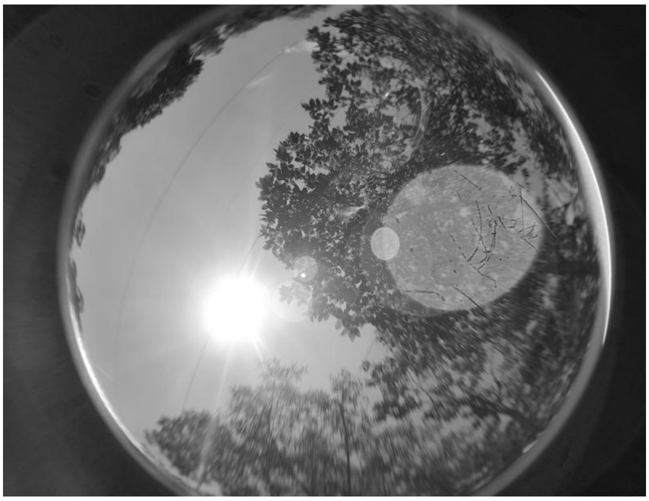

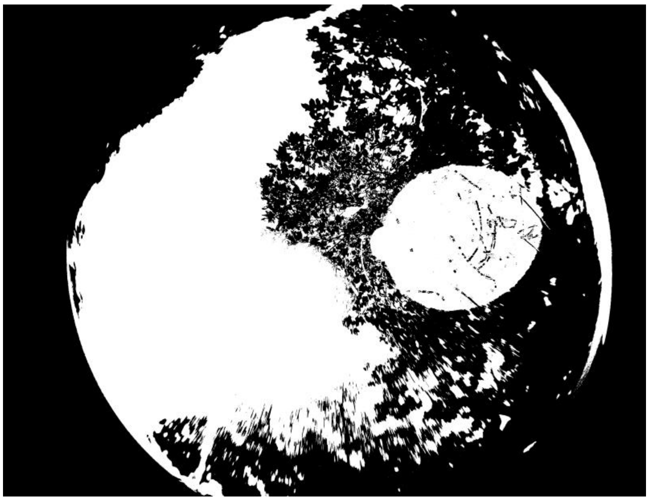

ID: IMG_20260217_113652.jpg | Sky: S (Gamma: 1.0) | Light: 51.35%

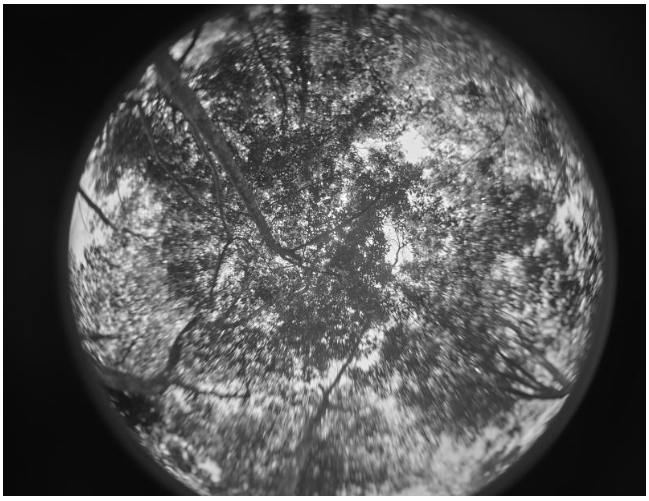

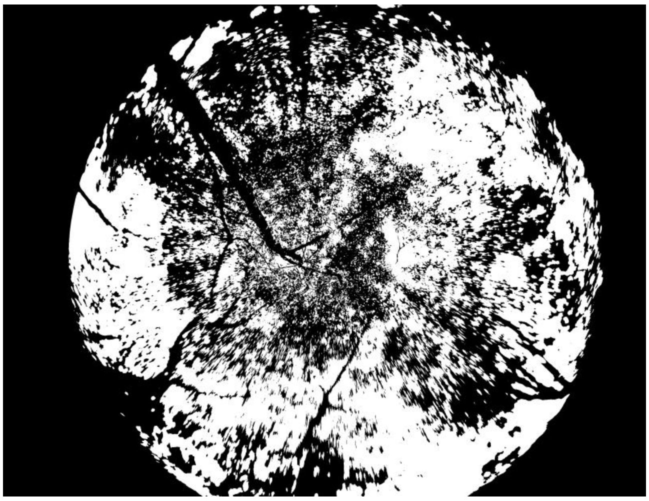

ID: IMG_20260222_083337.jpg | Sky: C (Gamma: 2.0) | Light: 78.77%

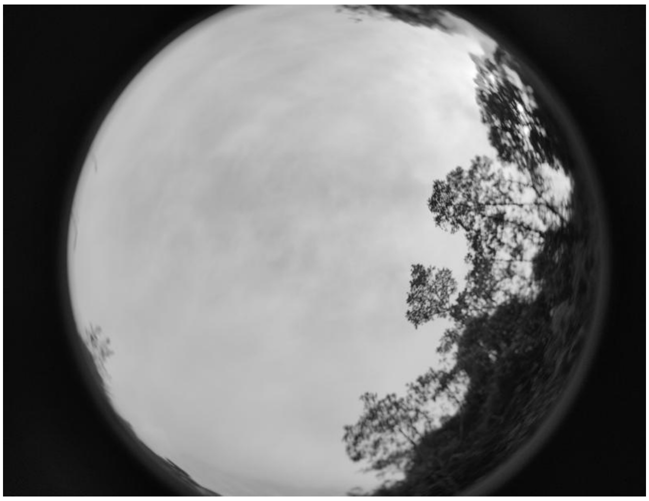

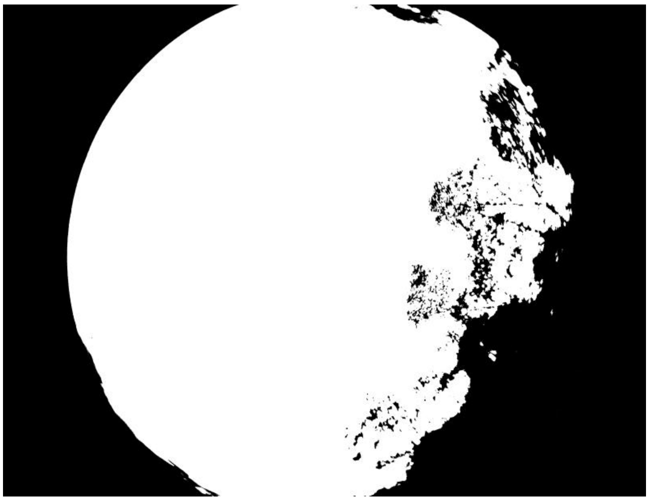

ID: IMG_20260222_084721.jpg | Sky: C (Gamma: 2.0) | Light: 34.62%

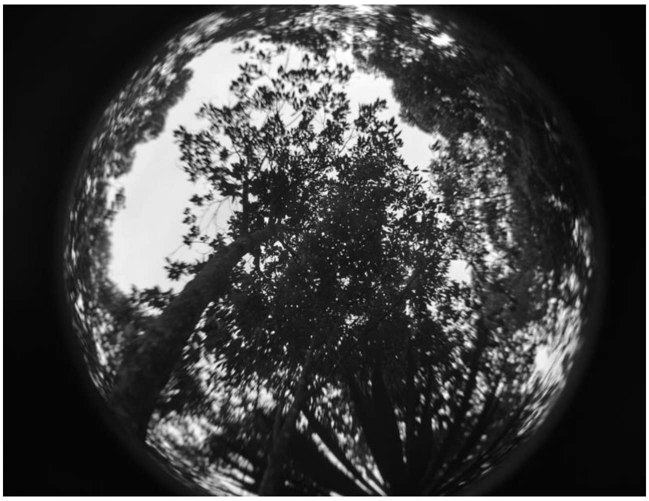

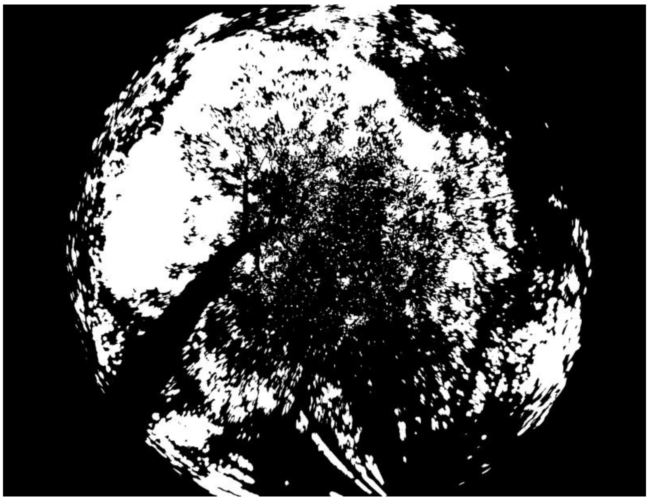

**ID: IMG_20260222_090231.jpg | Sky: C (Gamma: 2.0) | Light: 62.5%**

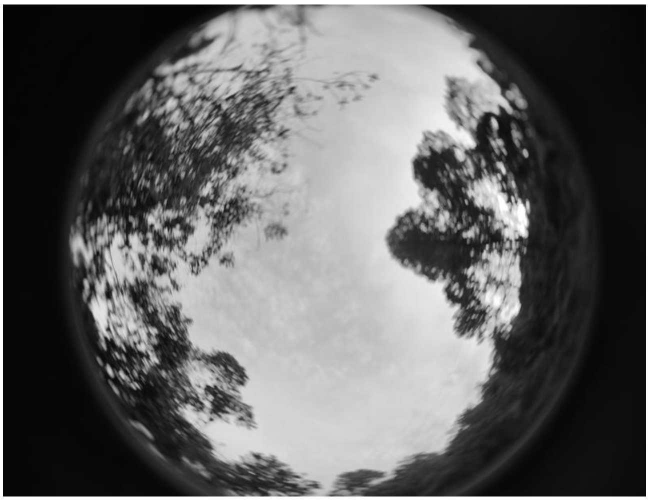

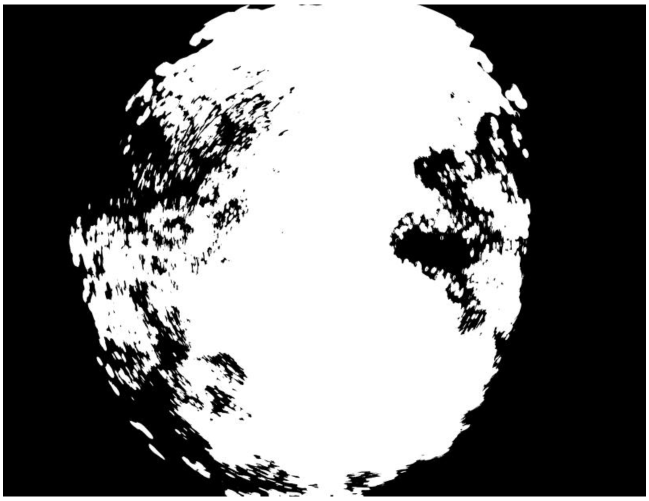

**ID: IMG_20260222_092308.jpg | Sky: P (Gamma: 1.5) | Light: 67.06%**

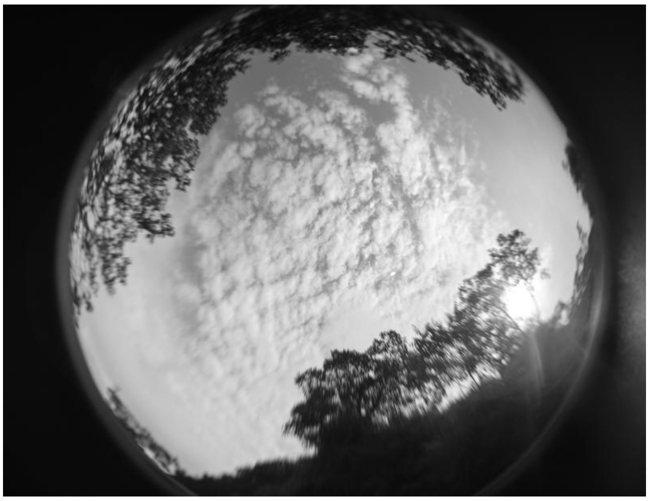

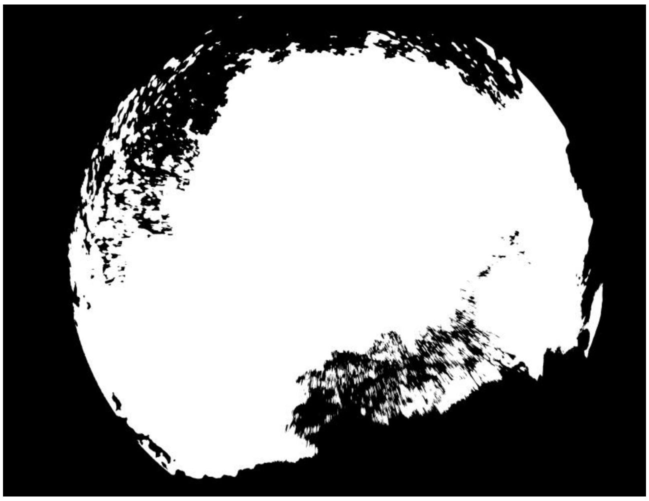

**ID: IMG_20260222_094008.jpg | Sky: P (Gamma: 1.5) | Light: 73.92%**

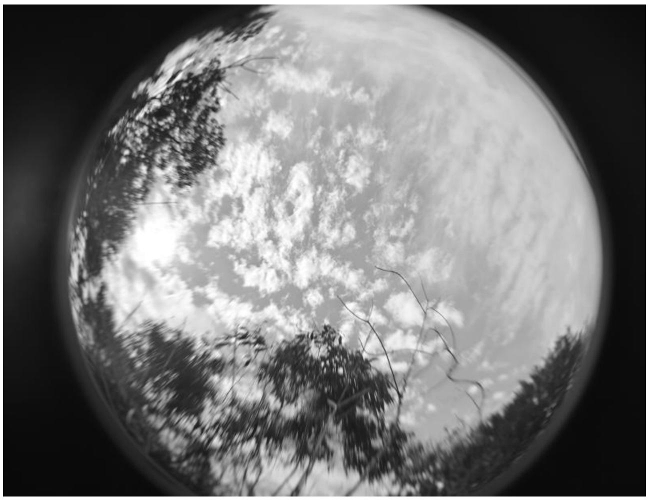

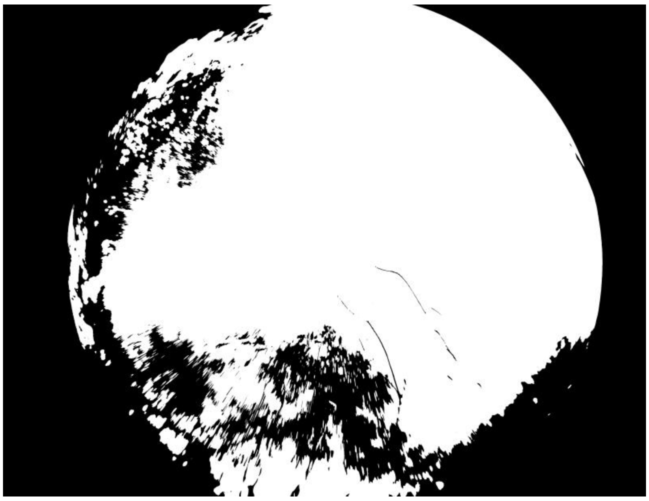

ID: IMG_20260222_095308.jpg | Sky: P (Gamma: 1.5) | Light: 94.32%

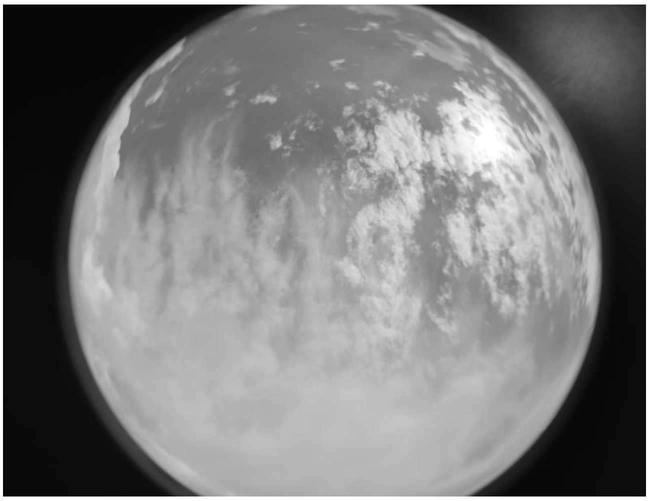

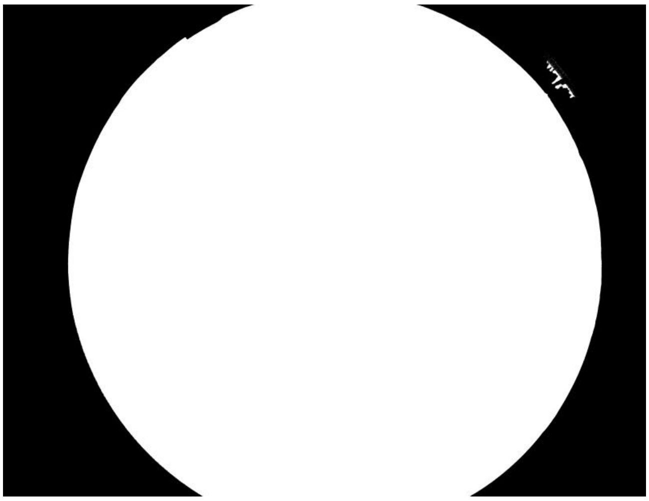

ID: IMG_20260222_101031.jpg | Sky: P (Gamma: 1.5) | Light: 94.33%

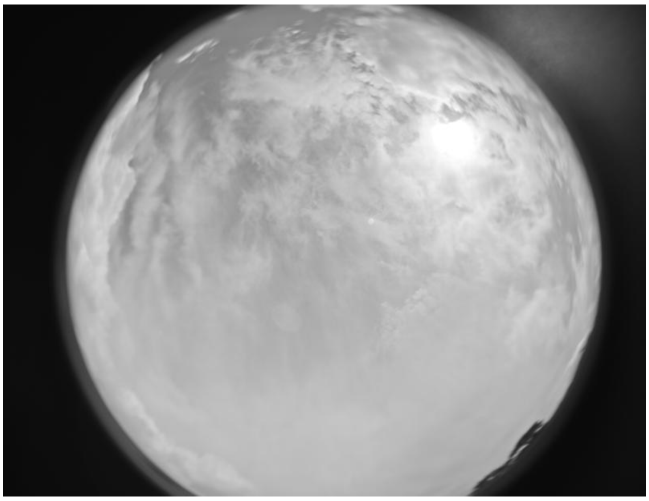

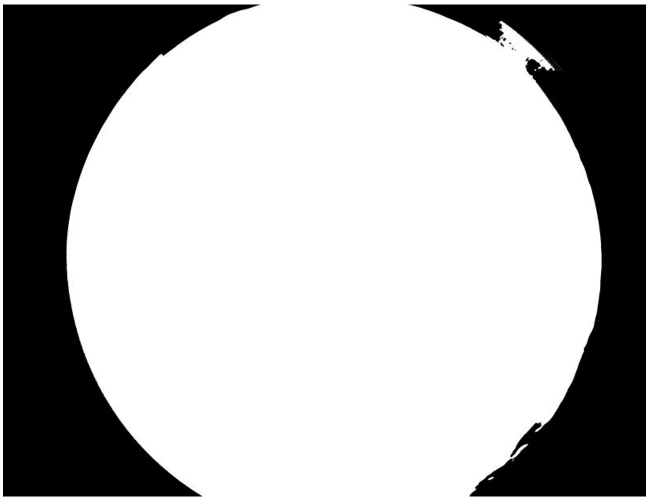

ID: IMG_20260222_102139.jpg | Sky: P (Gamma: 1.5) | Light: 91.22%

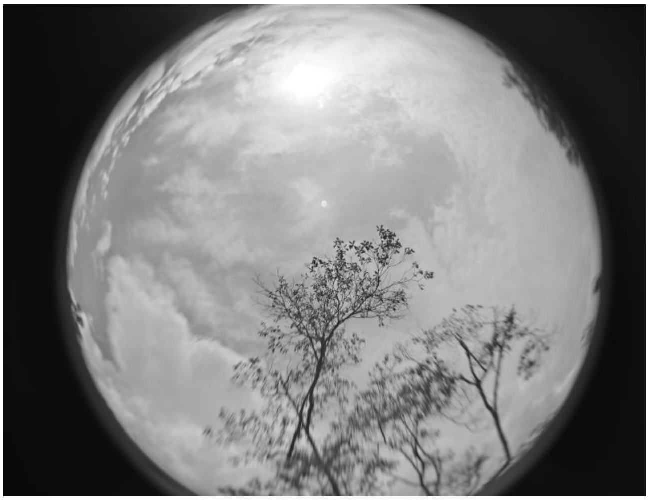

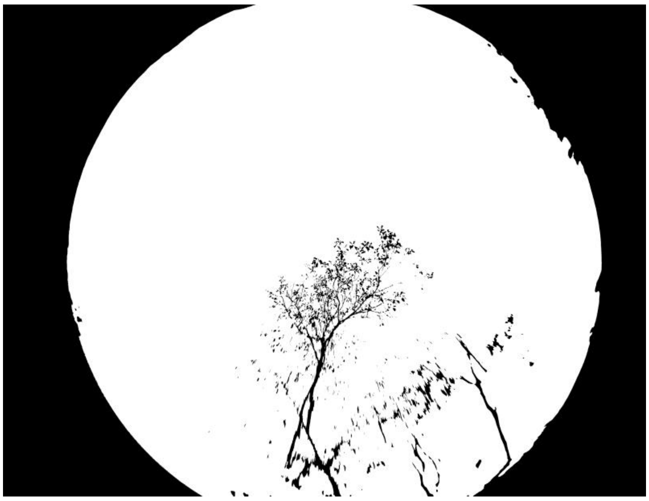

ID: IMG_20260222_103013.jpg | Sky: P (Gamma: 1.5) | Light: 82.88%

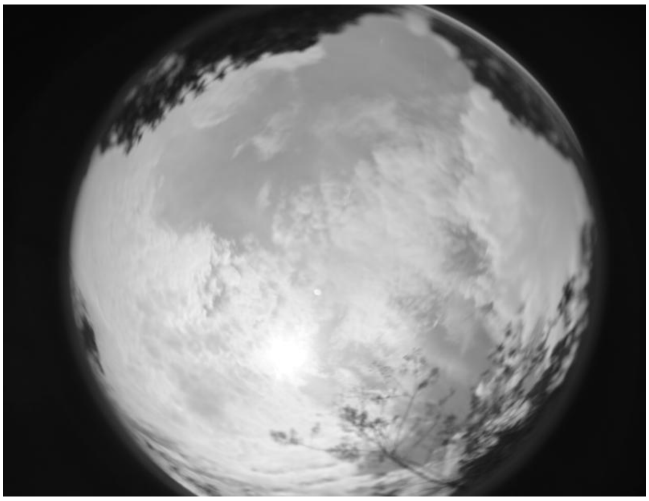
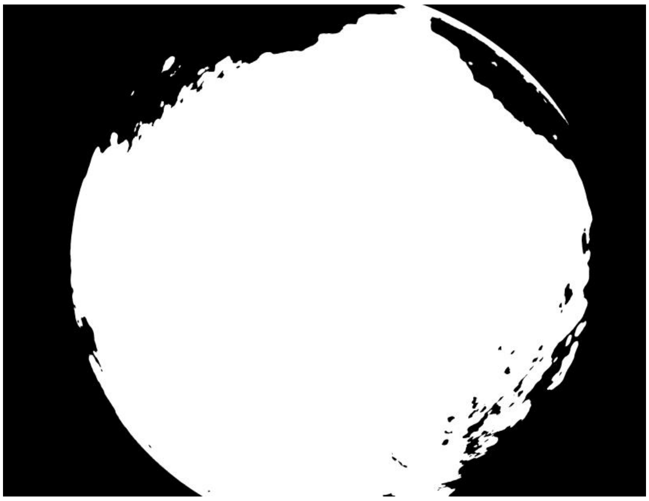

ID: IMG_20260222_104518.jpg | Sky: S (Gamma: 1.0) | Light: 32.73%

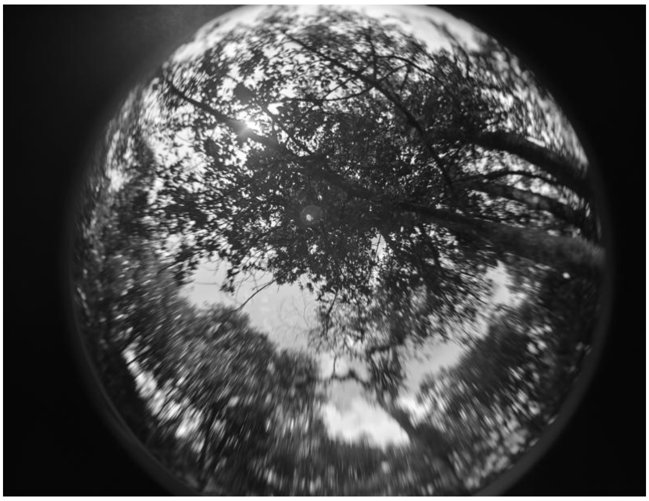
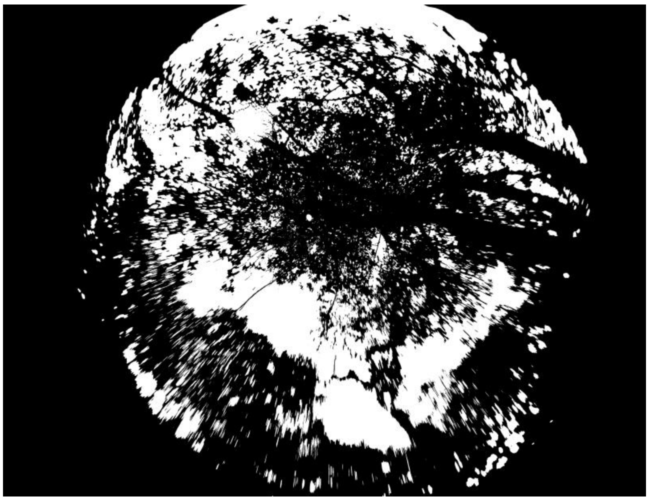

ID: IMG_20260222_110409.jpg | Sky: S (Gamma: 1.0) | Light: 47.61%

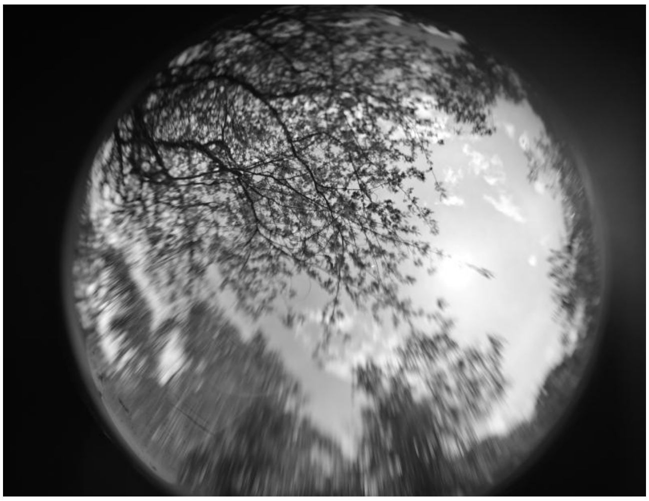
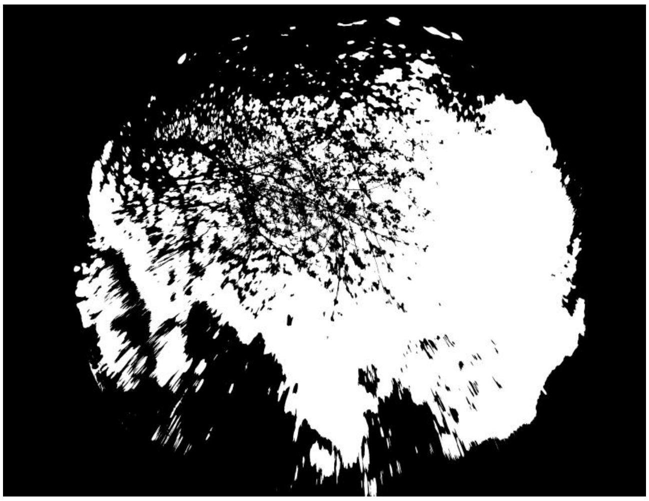

**ID: IMG_20260222_111403.jpg | Sky: S (Gamma: 1.0) | Light: 63.23%**

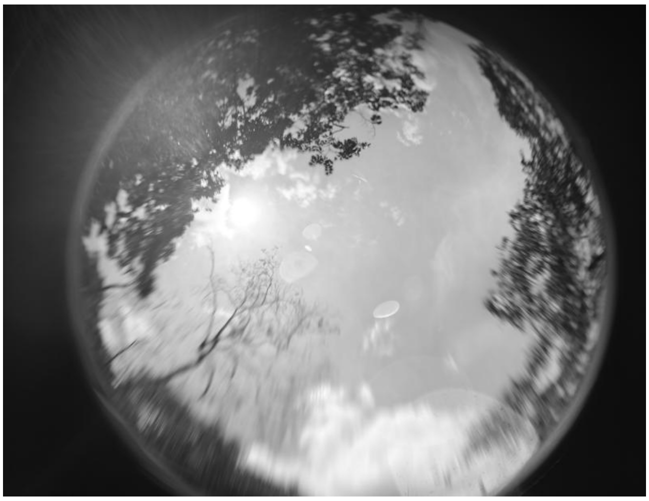
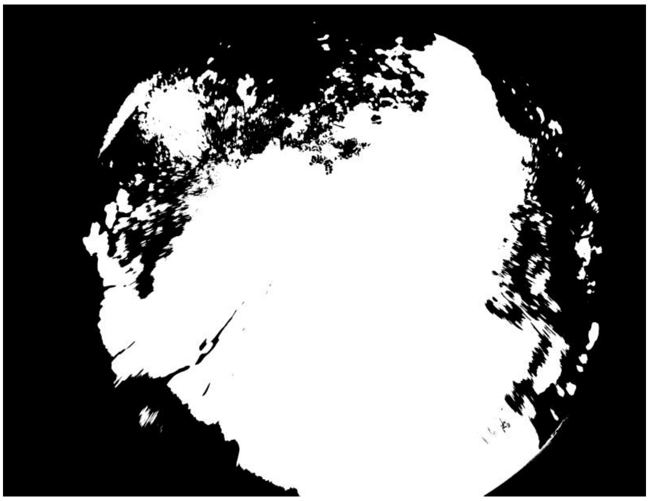

**ID: IMG_20260222_134256.jpg | Sky: P (Gamma: 1.5) | Light: 71.11%**

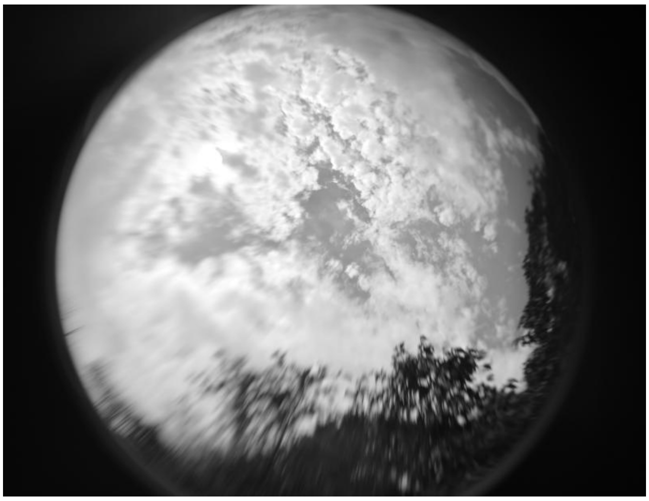
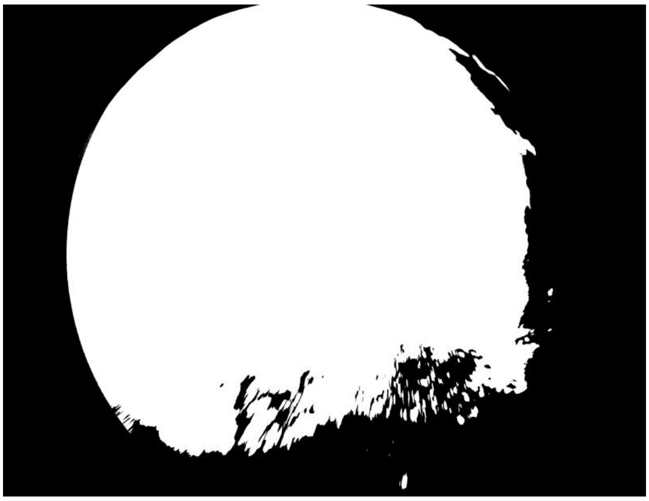

**ID: IMG_20260222_135558.jpg | Sky: S (Gamma: 1.0) | Light: 36.95%**

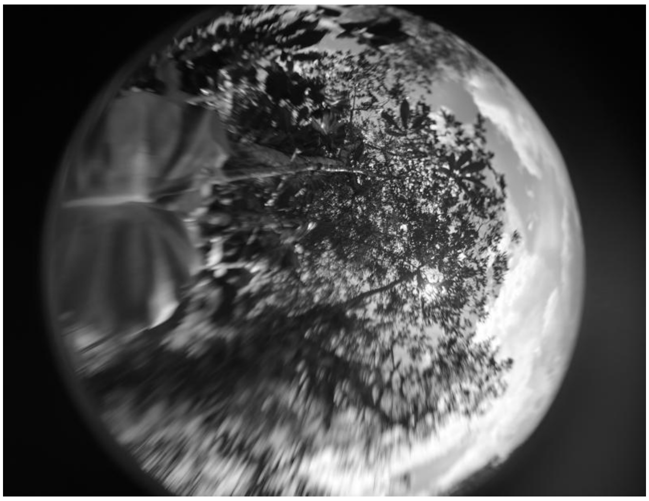
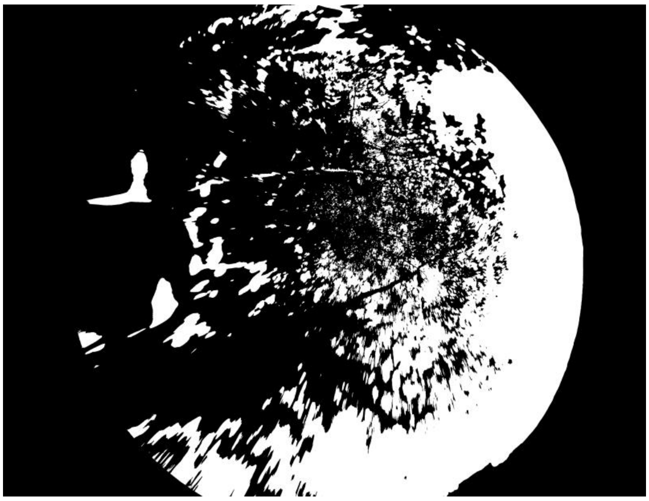

**ID: IMG_20260222_141211.jpg | Sky: S (Gamma: 1.0) | Light: 47.22%**

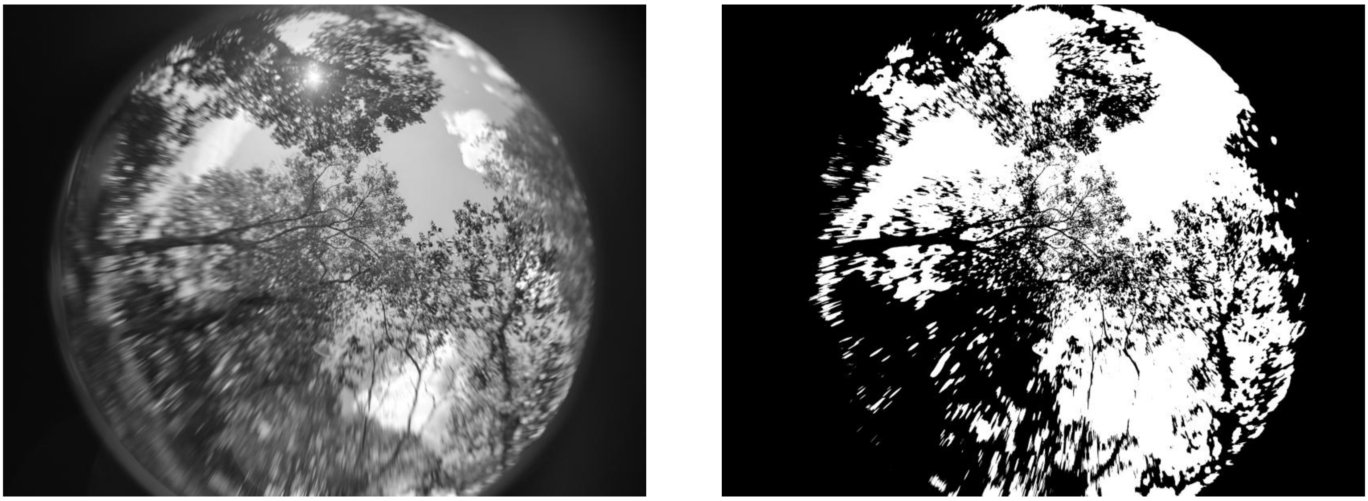

**ID: IMG_20260222_142227.jpg | Sky: S (Gamma: 1.0) | Light: 58.33%**

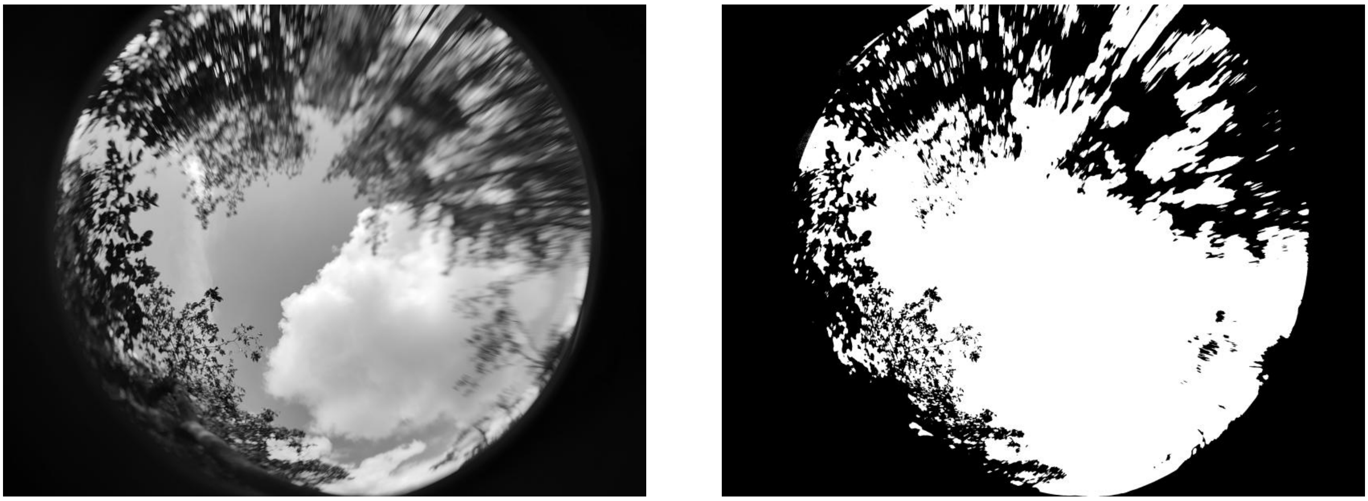

**ID: IMG_20260222_145629.jpg | Sky: C (Gamma: 2.0) | Light: 91.73%**

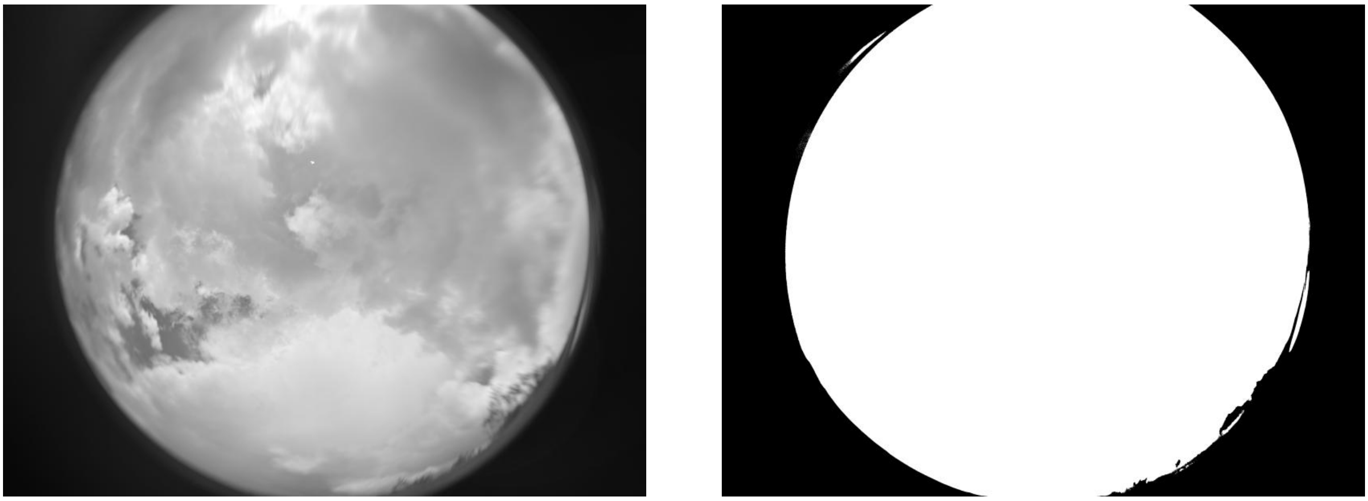
